## Supplementary File 1 for "The formation of the Indo-Pacific montane avifauna"

**Supplementary File 1.** Figs. S1-S15; Tables S1-S7.

### 1. Detailed results of population studies of montane supercolonizers

#### 1.1. Indo-Pacific *Phylloscopus* leaf warblers

Most of the Indo-Pacific *Phylloscopus* leaf warblers, as defined here, form a monophyletic group that is well supported across all analyses (Figs S4–S7). This clade includes all populations from the Philippines, Sulawesi, the Moluccas, New Guinea, and islands further east. Support for the broader group, which additionally includes *P. trivirgatus* (Greater and Lesser Sundas) and *P. presbytes* (Lesser Sundas), is slightly more equivocal (pp=0.97 in the supermatrix analysis). This broadly defined Indo-Pacific *Phylloscopus* leaf warbler clade has an estimated divergence date of 3.9 Ma (95% HPD 3.3–4.4 Ma). Estimated clade ages in our dated *Phylloscopus* tree are slightly older than those in previous studies (Price, 2010; Alström et al., 2018).

The results of the different phylogenetic analyses vary in some details, but are broadly similar. We first discuss the highly resolved tree generated from the genomic ML analysis. *P. presbytes* is sister to the rest of the clade. *P. trivirgatus* is next to diverge; *P. t. trivirgatus* appears to be paraphyletic with respect to *P. t. parvirostris* (Peninsular Malaysia) and *P. t. kinabaluensis* (n Borneo). Next to diverge is *P. nigrorum* from the Philippines, and then *P. sarasinorum nesophilus* (Sulawesi). The relationships recovered between Sulawesi and Moluccan taxa are less well supported than the rest of the topology (Fig. S6 Node A, bootstrap=80; Fig. S6 Node B, bootstrap=83). *P. maforensis waterstradti* (Obi and Bacan) is shown to be polyphyletic. Taxa from New Guinea and nearby islands, Kai Besar (Moluccas), the Bismarcks, and the Solomons form a well-supported group. However, *P. maforensis* is paraphyletic with respect to *P. makirensis* (Makira, Solomons), which is embedded deep within the clade as sister to *P. maforensis becki* (Guadalcanal and Malaita, Solomons).

In the supermatrix analysis, the early branching structure of the tree matches that of the genomic ML analysis, though with slightly lower support. However, the supermatrix analysis breaks up the Moluccan clade, recovering Seram (*P. maforensis ceramensis*) and Obi (*P. maforensis waterstradti*) populations as sister to the Philippines clade. Within the Philippines clade, an undescribed population from Mt. Busa (s Mindanao) does not group with other taxa from Mindanao, but rather has a well-supported sister relationship to the Panay population of *P. n. nigrorum*. This relationship was not commented upon in Jones & Kennedy (2008), from which the ND2 sequence derives. The topology recovered for Sulawesi taxa and the remaining Moluccan taxa (*P. nesophilus* and some *P. maforensis* subspecies) is quite different from the genomic ML analysis. This grouping unites taxa from Sulawesi, the Banggai and Sula Islands, and Buru, Bacan and Halmahera (Moluccas) in a well-supported clade. As in the genomic ML analysis, *P. maforensis waterstradti* (Obi and Bacan) is found to be polyphyletic. Taxa from New Guinea and nearby islands, Kai Besar (Moluccas), the Bismarcks, and the Solomons form a fairly well-supported group (pp=0.96), with *P. makirensis* embedded within *P. maforensis* as sister to *P. maforensis becki*. This analysis also shows that *P. amoenus*, endemic to Kolombangara in the Solomons, is deeply embedded within *P. maforensis*, clustering with other taxa from the Solomons and Bismarcks.

Our analyses reveal that current taxonomic treatments do not accurately reflect the evolution and diversity of the Indo-Pacific leaf warblers. We therefore made a preliminary revision of species limits in order to produce meaningful units for statistical analysis (Sections 4.1–4.5). We applied temporal banding (Avise & Johns, 2011) to the supermatrix tree to delimit species, setting the threshold for species-level divergence at 1.42 Ma. This corresponds to the divergence date estimate for the two taxa co-occurring on Kolombangara, *P. amoenus* and *P. maforensis pallescens*; 1.4 Ma is evidently adequate time to establish reproductive isolation between populations in this complex. We delimited 16 species within the complex (Fig. S4); the unsampled *P. rotiensis* likely represents an additional species. This treatment is entirely congruent with our genomic ML analysis, except for the inclusion of *P. s. nesophilus* (Sulawesi) in the species that also contains Halmahera and Bacan taxa. We note that further refinement of species limits will ultimately require the incorporation of vocalization and morphological data.

The Indo-Pacific leaf warbler radiation originated some 4 Ma, and appears to have reached as far as the Philippines and New Guinea around the start of the Pleistocene 2.6 Ma (Fig. S4, Supplementary File 4). Diversification surged in the early Pleistocene and continued through most of the epoch, though inter-island diversification seems to have tapered off slightly within the last 200–300 Ka. The complex evolved from Palearctic/Indomalayan ancestors (Supplementary File 4), and the genomic ML tree indicates a west-to-east stepping-stone expansion out of continental Asia. This includes an ambiguous double colonization of the Lesser Sundas, an expansion into the Philippines, and a broad eastward colonization proceeding through Sulawesi, the Moluccas, New Guinea, and the Bismarcks and Solomons. Indo-Pacific leaf warblers are overwhelmingly montane, but reach the lowlands of Rote and Timor (Lesser Sundas), Numfor and Biak/Supiori (off northern New Guinea), and Mussau (Bismarcks). Except for Timor, these are all small, low-lying islands. The *Phylloscopus* elevational range reconstruction (Supplementary File 4) and the results of the phylogenetic analyses indicate that these represent 3-4 separate ecological shifts from the ancestral 'montane' state. All members of the complex are essentially sedentary. The *Phylloscopus* migratory behavior reconstruction (Supplementary File 4) indicates that the last common ancestor (3.9 Ma) was sedentary, but that ancestors from 4.2 Ma and earlier were short-distance migrants.

#### 1.2. Snowy-browed Flycatcher *Ficedula hyperythra*

*Ficedula hyperythra* as currently delimited is paraphyletic, as our results reveal that Damar Flycatcher *F. henrici* is embedded within it (Figs. S8–S11). All analyses except the mitogenome analysis recover a well-supported sister relationship between *F. henrici* and *F. h. audacis* from the nearby small island of Babar in the Lesser Sundas. All analyses additionally recover a deep divergence between *F. h. annamensis* of southern Vietnam and the rest of *F. hyperythra*, dated at 3.4 Ma (95% HPD 2.4–4.7 Ma) in the supermatrix analysis. While this strongly indicates species-level divergence, we continue to treat subspecies *annamensis* as part of *F. hyperythra* pending an integrative taxonomic assessment. The supermatrix analysis shows that *F. hyperythra* individuals from northern Vietnam are not closely related to *F. h. annamensis*, but a recently discovered population from the Cardamom Mountains of Cambodia is considered to belong to this subspecies (Eames et al., 2002). The supermatrix analysis recovers a weakly supported (posterior probability = 0.72) sister relationship between *F. hyperythra* and the Sino-Himalayan *F. tricolor*, with a divergence date of 14 Ma (not shown); this relationship was strongly supported by Moyle et al. (2015) and Hooper et al. (2016). The *F. hyperythra* crown group (excluding *F. h. annamensis*) began to diversify at 1.4 Ma (95% HPD 1.1–1.8 Ma).

The supermatrix analysis shows explosive radiation of the group through the Greater Sundas and Wallacean islands beginning at around 1 Ma. It appears that most modern island populations formed during this initial burst, and that inter-island diversification slowed significantly thereafter. Given these temporal diversification dynamics, it is unsurprising that this tree is rather poorly resolved beyond the earliest branches. The genomic ML tree is reasonably well resolved, but receives lower overall support than similar analyses for the other montane supercolonizers. The geographic range reconstruction for the genus *Ficedula* (Supplementary File 4) strongly indicates that *F. hyperythra* evolved from Palearctic ancestors. The finding that *F. h. annamensis* from south Vietnam is a well-diverged sister to the rest of *F. hyperythra* blurs this picture slightly, but the next branch of the tree represents populations spanning the Palearctic and Indomalayan regions, indicating that this is the ancestral range of the species. From here it appears to have expanded into the Greater Sundas. From the Greater Sundas, the genomic ML analysis indicates that *F. hyperythra* made two separate eastward expansions: one via Sulawesi into the Moluccas, and one through the Lesser Sundas. Two populations on very small islands in the eastern Lesser Sundas occur down to low elevations. On Babar it has been recorded down to 200 masl (Trainor & Verbelen, 2013), and on Damar down to 60 masl (Trainor, 2007). Examined together with the *Ficedula* elevational range reconstruction (Supplementary File 4), the results of the phylogenetic analyses strongly suggest that Babar and Damar populations, and *F. hyperythra* as a whole, evolved from montane ancestors. *F. hyperythra* appears to have evolved from short-distance migrants, based on the *Ficedula* migration reconstructions (Supplementary File 4) and inclusion of the altitudinal migrant Himalayan population in an early branching clade of *F. hyperythra* (Fig S8).

#### 1.3. Mountain Tailorbird *Phyllergates cucullatus*

The supermatrix analysis recovered a monophyletic *Phyllergates cucullatus* (Figs. S12–S15). Divergence from its sister species, *P. heterolaemus* from Mindanao, was estimated at 4.1 Ma (95% HPD 1.8–6.9 Ma). Populations of *P. cucullatus* were estimated to have begun diverging 1.3 Ma (95% HPD 1.0–1.7 Ma). The mainland *P. c. coronatus* may be paraphyletic, based on the highly supported topology from the genomic ML analysis. This analysis groups north and south Vietnamese *P. c. coronatus* populations together as sister to the rest of *P. cucullatus.* Next to diverge are Myanmar and northeast India *P. c. coronatus* populations, which are sister to the remaining taxa. Among Moluccan taxa, all analyses recover a sister relationship between *P. c. dumasi* (Seram) and *P. c. batjanensis* (Bacan), which are in turn sister to *P. c. dumasi* (Buru). These relationships are well supported in all but the supermatrix analysis, indicating that *P. c. dumasi* is paraphyletic.

*Phyllergates cucullatus* underwent a recent archipelagic radiation beginning c. 1 Ma, similar to *F. hyperythra*. Inter-island diversification seems to have proceeded at a somewhat slower rate than in *F. hyperythra*, occurring from 1 to 0.3 Ma and even more recently. As a consequence of this slower rate of diversification, the supermatrix tree is slightly better supported, and the genomic ML tree is well resolved. The geographic range reconstruction for Cettiidae (Supplementary File 4) provides somewhat equivocal results as to the ancestral origin of *P. cucullatus*, due to geographic signal from its Philippines-endemic sister species; *Phyllergates* belongs to an otherwise Palearctic/Indomalayan clade. The genomic ML analysis results suggest that *P. cucullatus* evolved from a Palearctic/Indomalayan ancestor, as populations occurring there represent early branches of the tree. From this probable ancestral range, *P. cucullatus* underwent a range expansion similar to that of *F. hyperythra*. It appears to have expanded through the Greater Sundas, and from there to the northern Philippines; east into the Moluccas via Sulawesi; and east into the Lesser Sundas. The elevational range reconstructions coupled with the phylogenetic analyses provide a picture of an entirely montane radiation. The migratory behavior of the species' last common ancestor remains unclear, but the altitudinal migrant population in northeast India belongs to one of the early branching groups (Fig. S12).

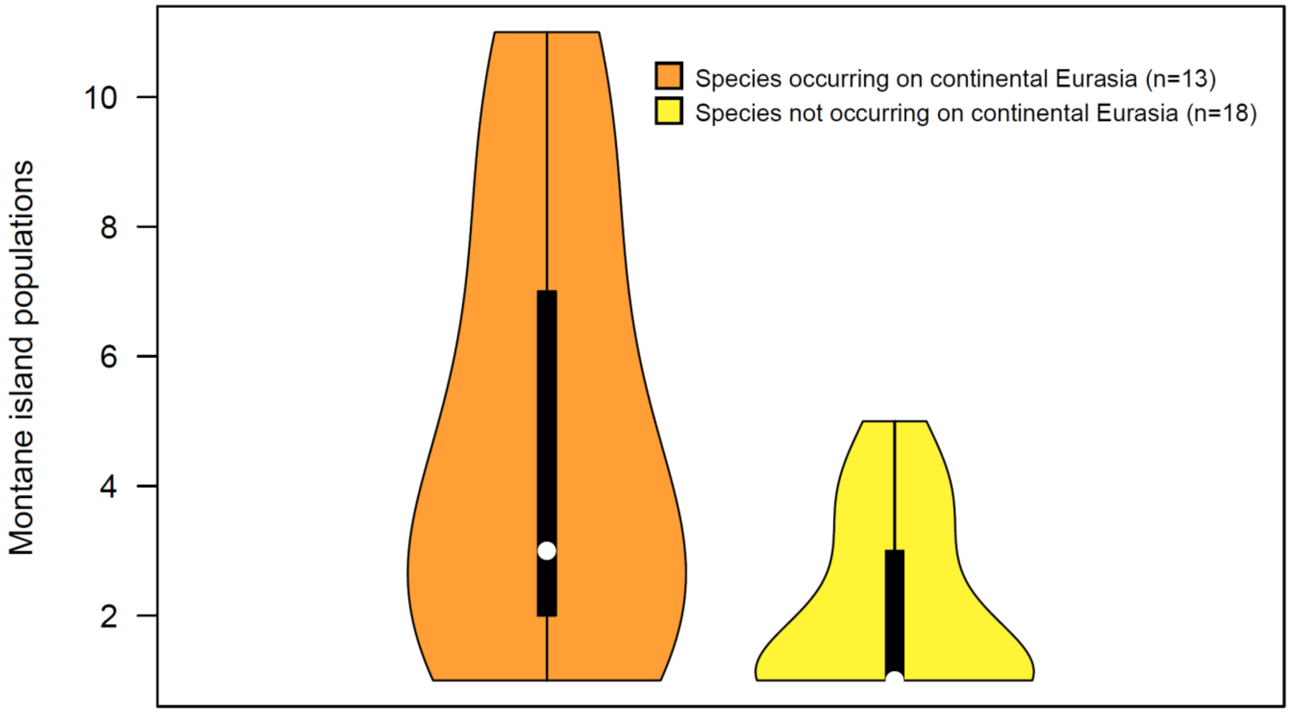


**Fig. S1.** Number of MIPs per Eurasian-origin species which respectively do, or do not, have ranges including continental Eurasia. The fact that species with continental occurrence have more MIPs may indicate that lineages' colonization capacities decline with longer residence in tropical archipelagos.

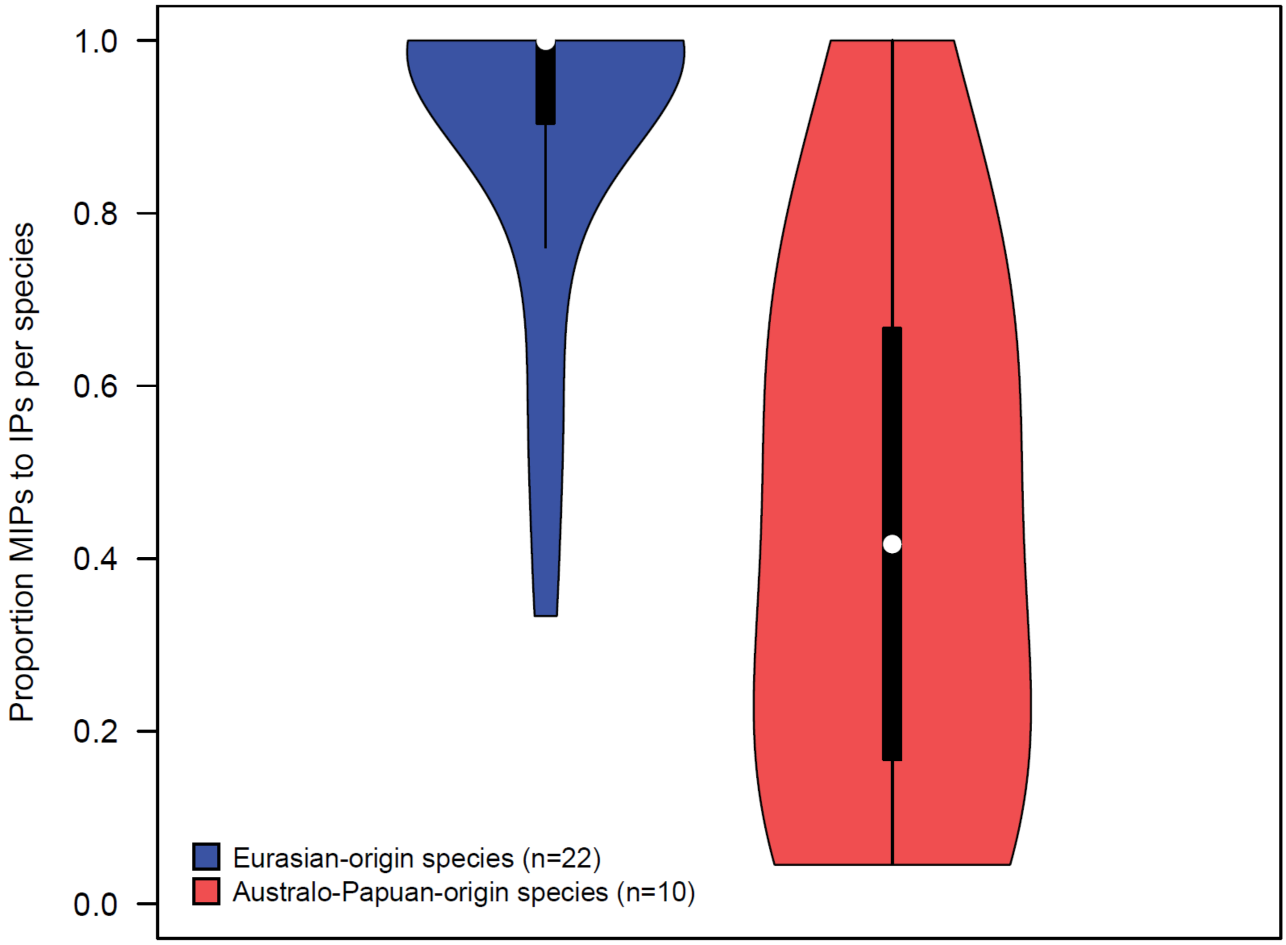


**Fig. S2.** Proportion of MIPs to total island populations (IPs) per species, for Eurasian-origin versus Australo-Papuan-origin species. Species with one MIP and no LIPs are excluded. Eurasian-origin species with MIPs are consistently montane across their archipelagic ranges, but Australo-Papuan-origin species are not.

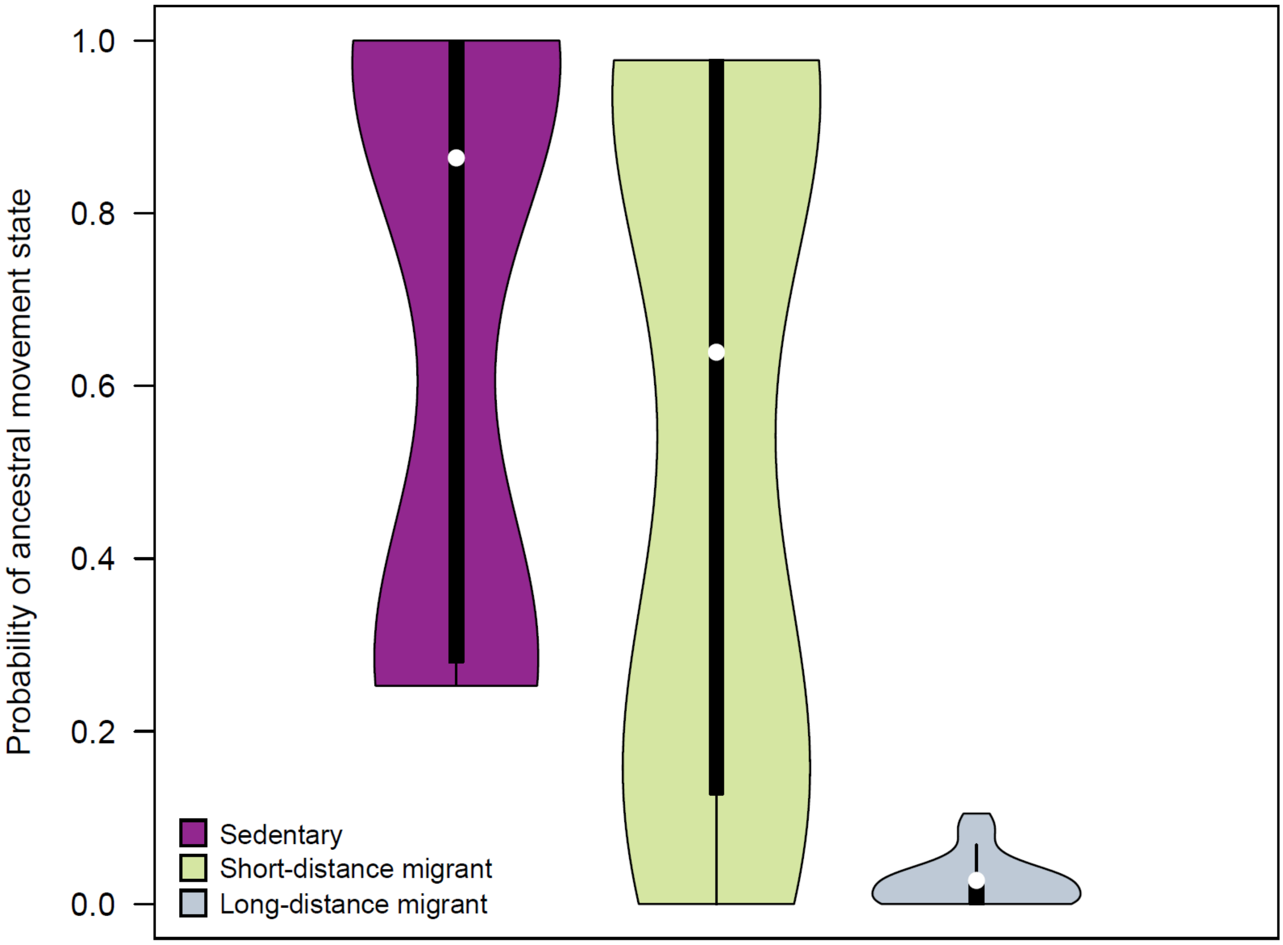


**Fig. S3.** Probabilities of ancestral migratory behavior states for Eurasian-origin species with MIPs. Values are derived from the probabilities at the Ancestral Source Nodes of all species, as described in Section 4.5.4. Note that migratory behavior classes are not mutually exclusive for species or ancestral nodes, as different populations within a single species can show different migratory behavior. Australo-Papuan-origin species are not shown because their continental ancestors were almost entirely sedentary.

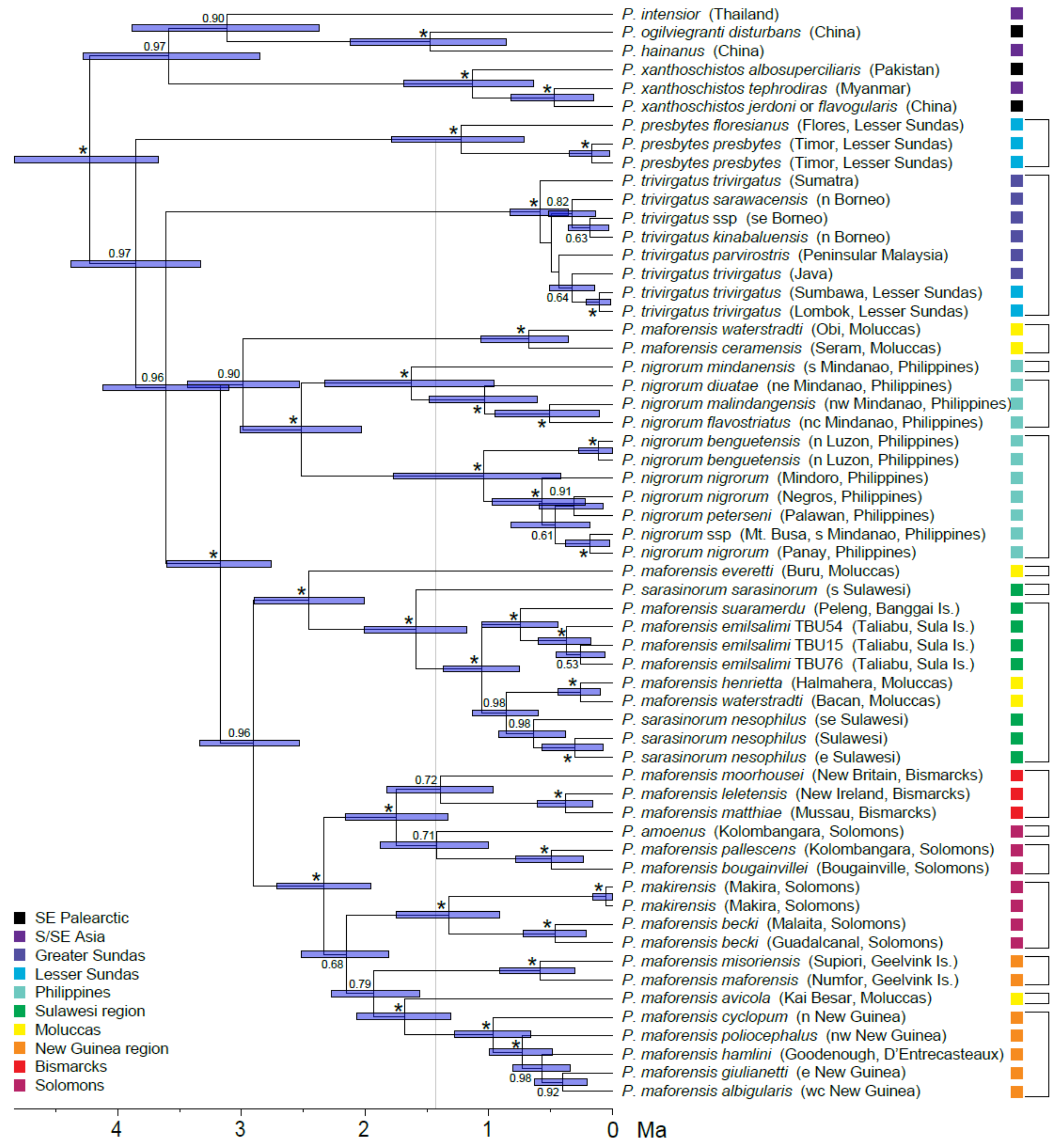


**Fig. S4.** Indo-Pacific *Phylloscopus* leaf warblers: Time-calibrated Bayesian consensus tree from supermatrix analysis of three nuclear and two mitochondrial loci. Posterior probabilities ≥ 0.50 are indicated at the nodes; asterisks denote nodes with PP ≥ 0.99. Error bars indicate 95% highest posterior density (HPD) intervals. A four-species sister clade is shown, but other outgroups are not. Colored cells indicate areas of occurrence (breeding ranges). We used temporal banding to make a preliminary revision of species limits for use in this study. The threshold for species-level divergence was set at 1.42 Ma (indicated with a gray line) based on the divergence date estimate for the two sympatric species on Kolombangara, *P. amoenus* and *P. maforensis pallescens*. Species membership as defined for this study is indicated with brackets.

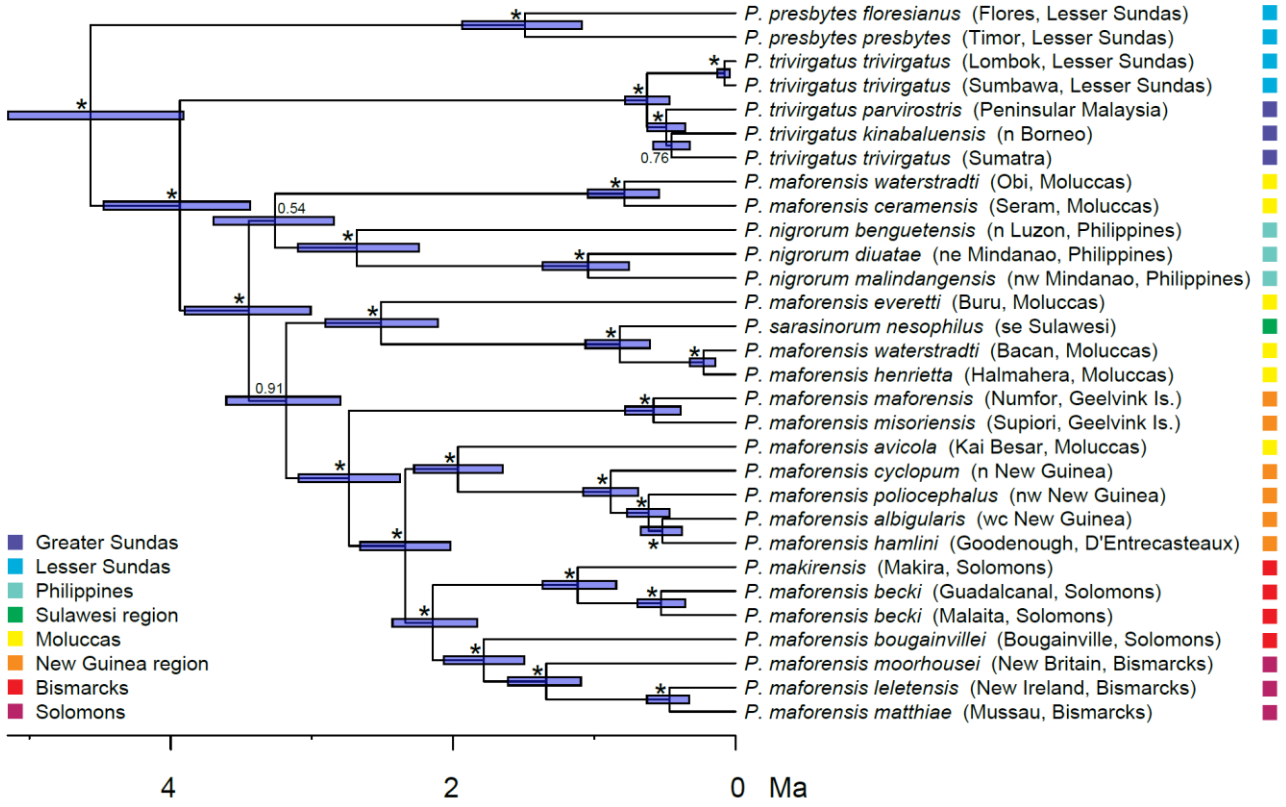


**Fig. S5.** Indo-Pacific *Phylloscopus* leaf warblers: Time-calibrated Bayesian consensus tree from analysis of mitochondrial genomes. Posterior probabilities ≥ 0.50 are indicated at the nodes; asterisks denote nodes with PP ≥ 0.99. Error bars indicate 95% highest posterior density (HPD) intervals. The single outgroup taxon is not shown. Colored cells indicate areas of occurrence (breeding ranges).

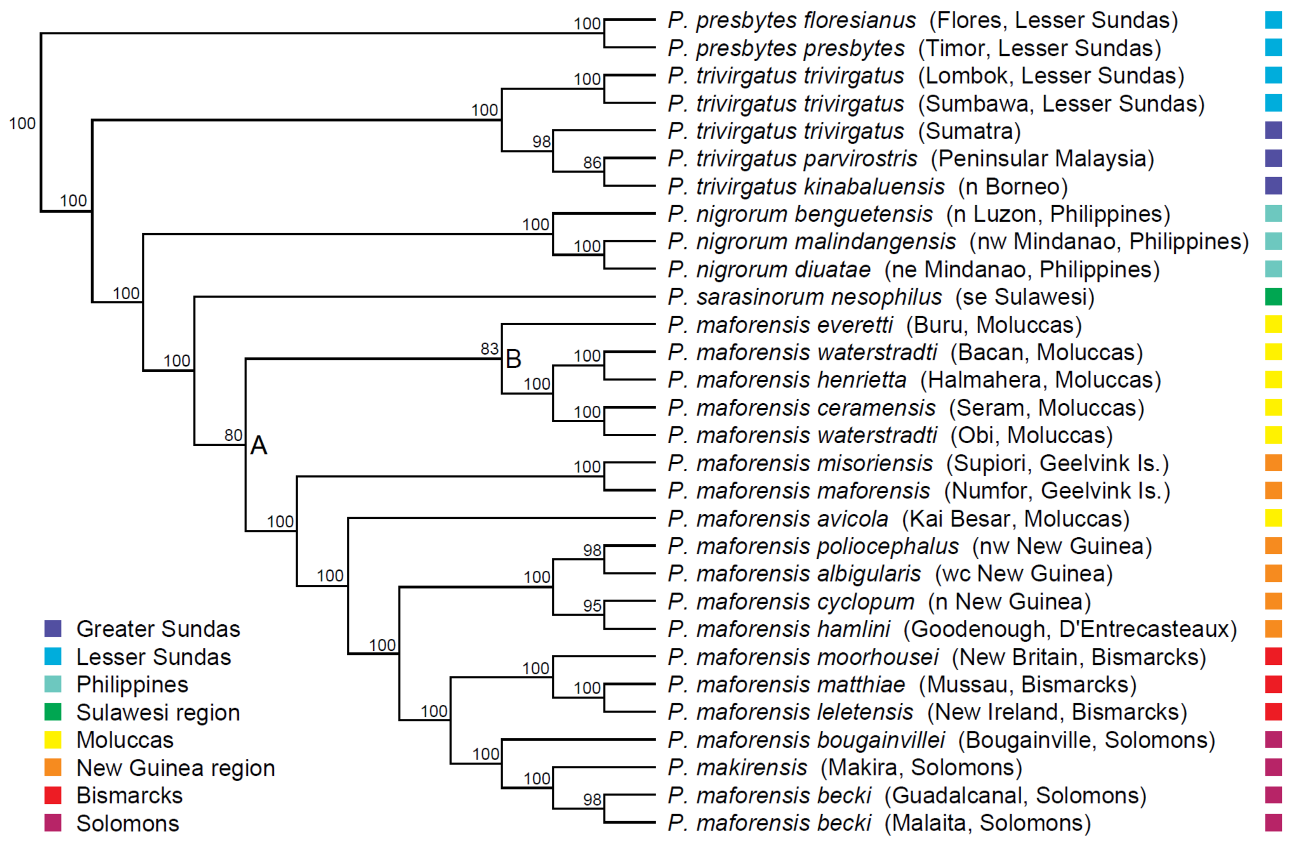


**Fig. S6.** Indo-Pacific *Phylloscopus* leaf warblers: Concatenated genomic maximum likelihood tree. ML bootstrap supports ≥ 50 are indicated at the nodes. Nodes labelled with letters are discussed in the text. Outgroup taxa are not shown. Colored cells indicate areas of occurrence (breeding ranges).

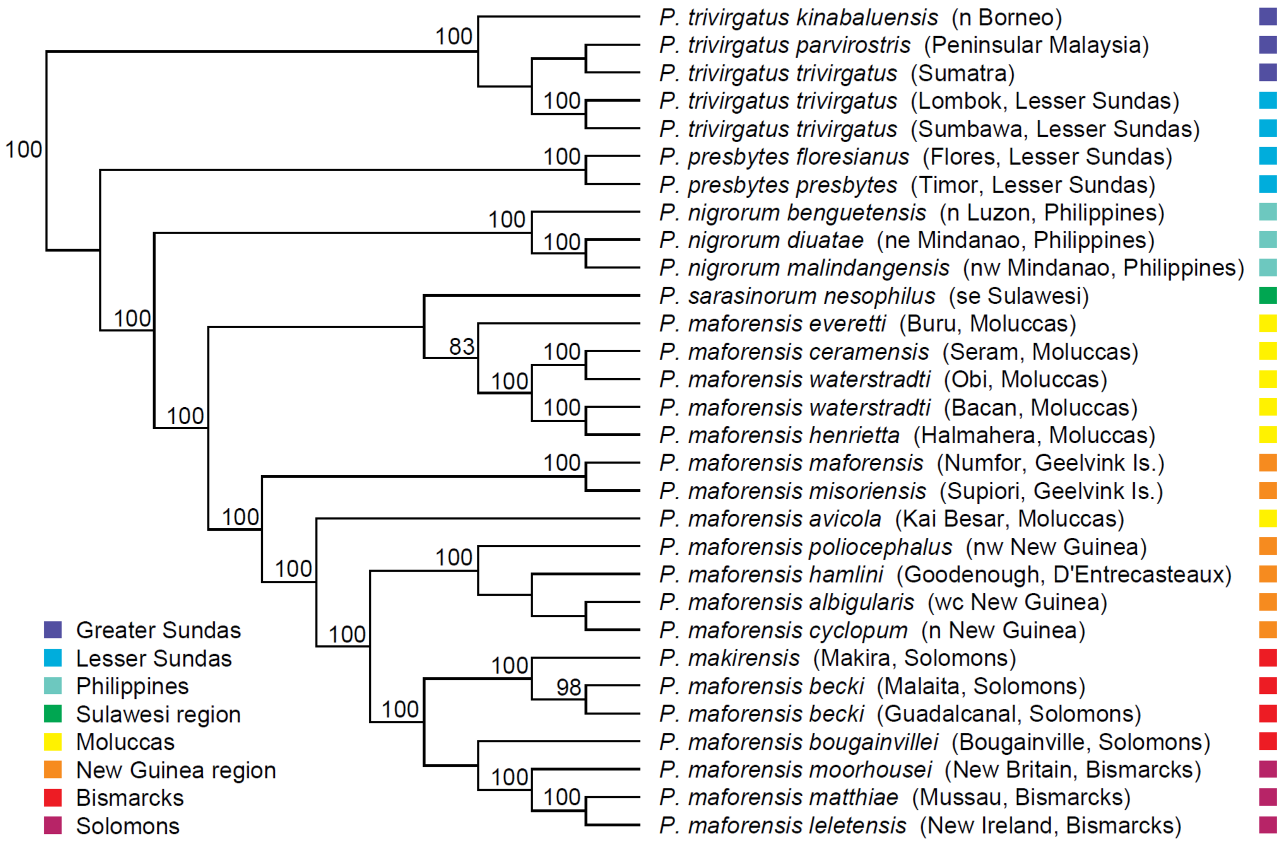


**Fig. S7.** Indo-Pacific *Phylloscopus* leaf warblers: Genomic species tree with maximum likelihood bootstrap values. ML bootstrap supports ≥ 50 are indicated at the nodes. Outgroup taxa are not shown. Colored cells indicate areas of occurrence (breeding ranges).

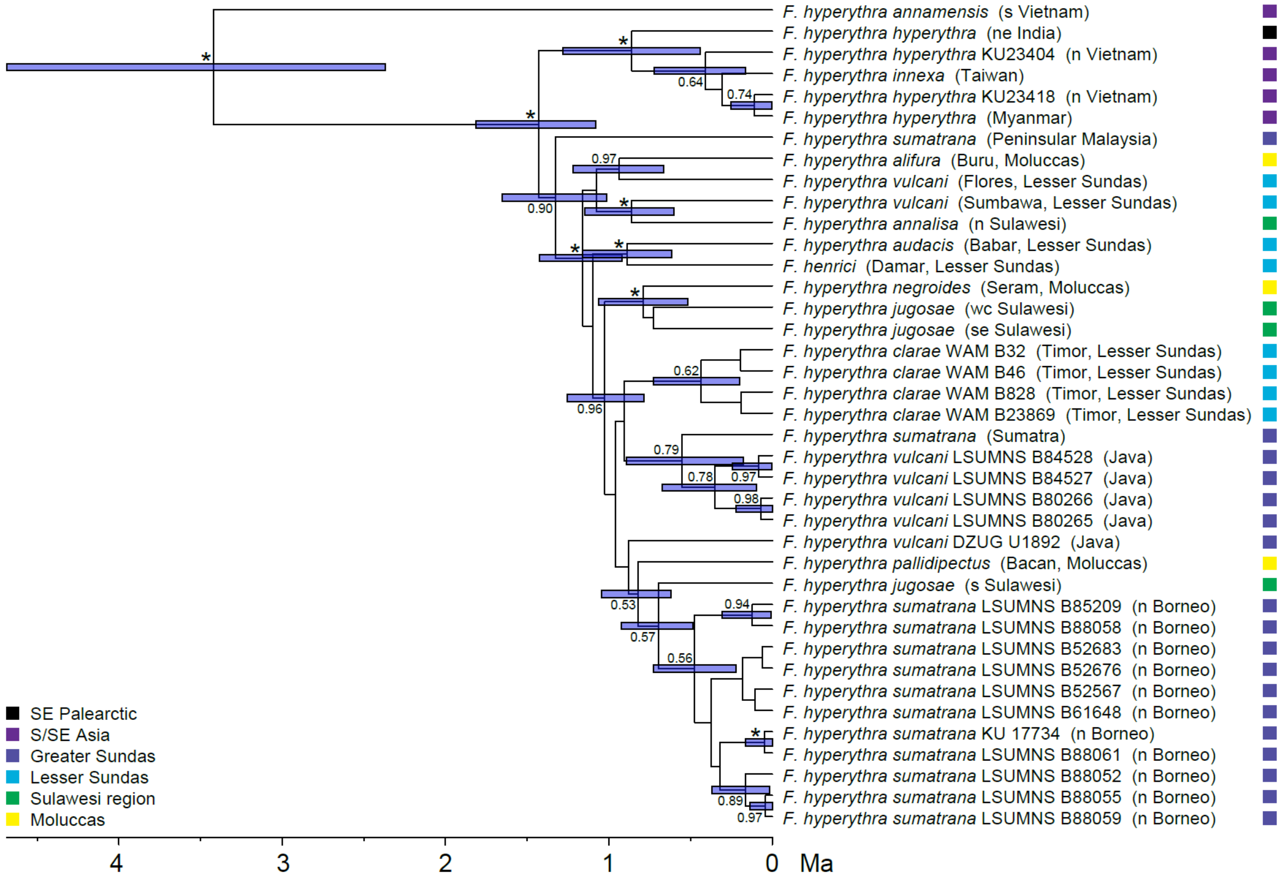


**Fig. S8.** Snowy-browed Flycatcher *Ficedula hyperythra*: time-calibrated Bayesian consensus tree from supermatrix analysis of three nuclear and three mitochondrial loci. Posterior probabilities ≥ 0.50 are indicated at the nodes; asterisks denote nodes with PP ≥ 0.99. Error bars indicate 95% highest posterior density (HPD) intervals. Outgroup taxa are not shown. Colored cells indicate areas of occurrence (breeding ranges).

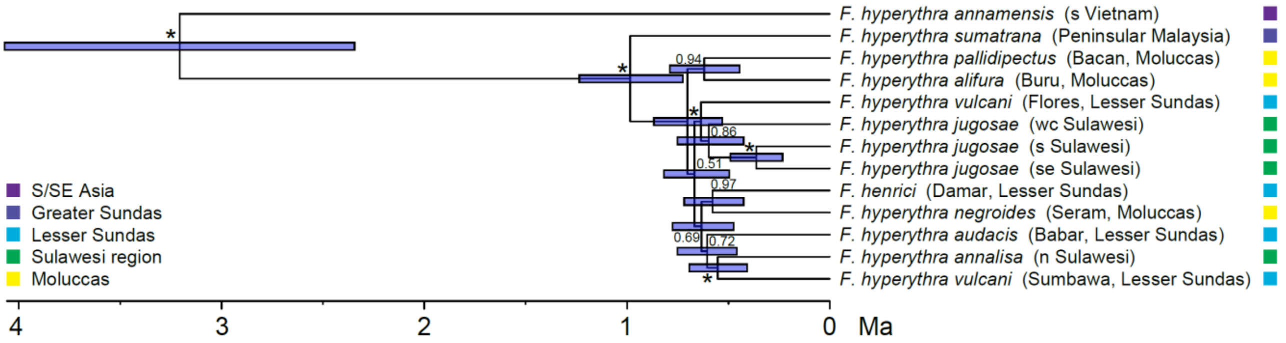


**Fig. S9.** Snowy-browed Flycatcher *Ficedula hyperythra*: Time-calibrated Bayesian consensus tree from analysis of mitochondrial genomes. Posterior probabilities ≥ 0.50 are indicated at the nodes; asterisks denote nodes with PP ≥ 0.99. Error bars indicate 95% highest posterior density (HPD) intervals. Outgroup taxa are not shown. Colored cells indicate areas of occurrence (breeding ranges).

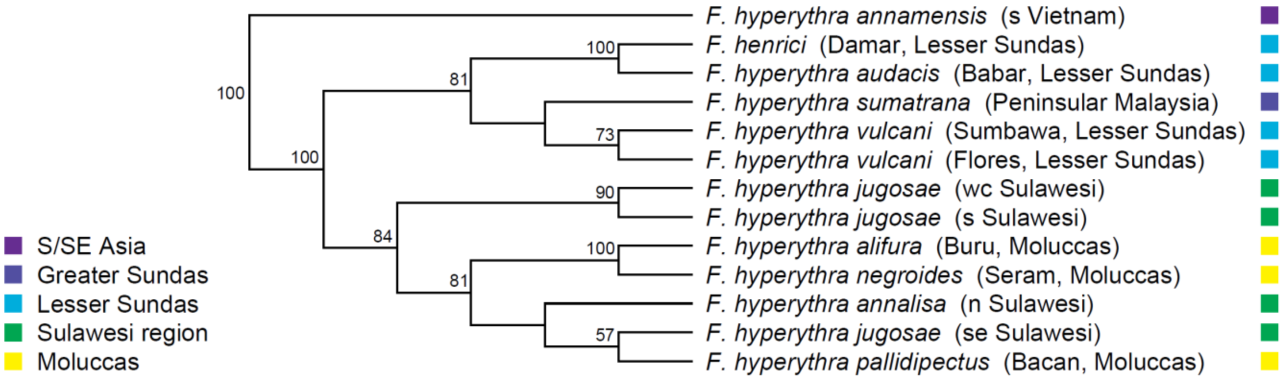


**Fig. S10.** Snowy-browed Flycatcher *Ficedula hyperythra*: Concatenated genomic maximum likelihood tree. ML bootstrap supports ≥ 50 are indicated at the nodes. Outgroup taxa are not shown. Colored cells indicate areas of occurrence (breeding ranges).

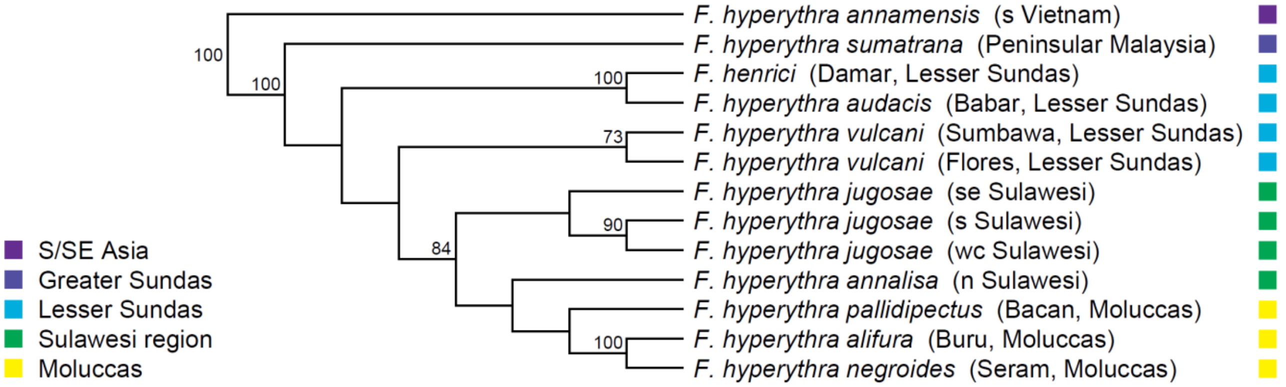


**Fig. S11.** Snowy-browed Flycatcher *Ficedula hyperythra*: Genomic species tree with maximum likelihood bootstrap values. ML bootstrap supports ≥ 50 are indicated at the nodes. Outgroup taxa are not shown. Colored cells indicate areas of occurrence (breeding ranges).

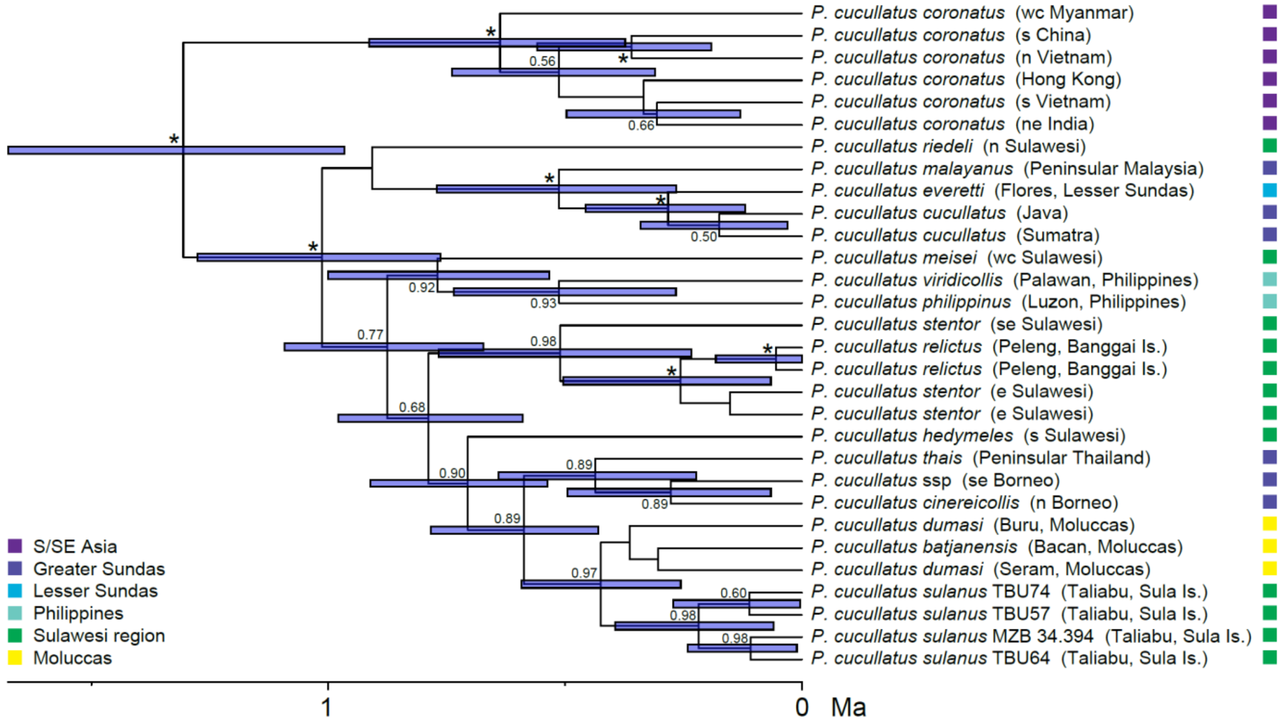


**Fig. S12.** Mountain Tailorbird *Phyllergates cucullatus*: time-calibrated Bayesian consensus tree from supermatrix analysis of five nuclear and two mitochondrial loci. Posterior probabilities ≥ 0.50 are indicated at the nodes; asterisks denote nodes with PP ≥ 0.99. Error bars indicate 95% highest posterior density (HPD) intervals. Outgroup taxa are not shown. Colored cells indicate areas of occurrence (breeding ranges).

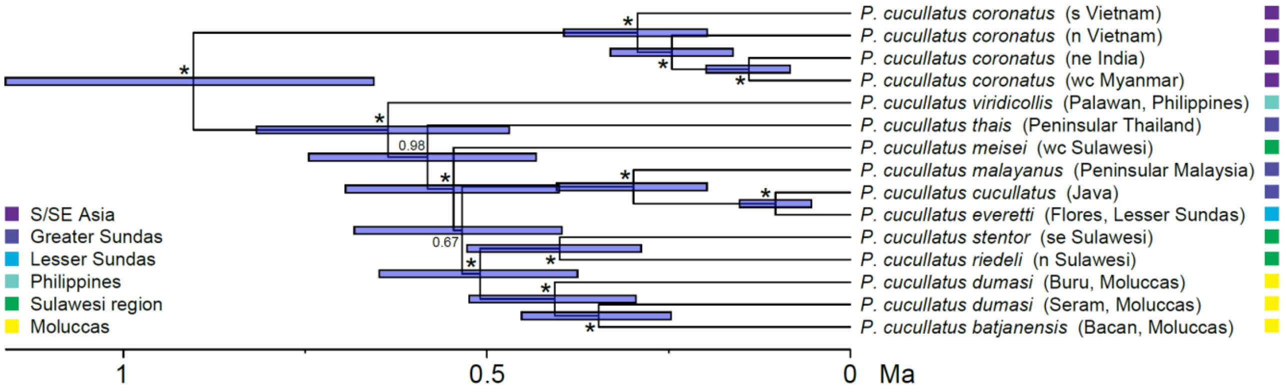


**Fig. S13.** Mountain Tailorbird *Phyllergates cucullatus*: Time-calibrated Bayesian consensus tree from analysis of mitochondrial genomes. Posterior probabilities ≥ 0.50 are indicated at the nodes; asterisks denote nodes with PP ≥ 0.99. Error bars indicate 95% highest posterior density (HPD) intervals. The single outgroup taxon is not shown. Colored cells indicate areas of occurrence (breeding ranges).

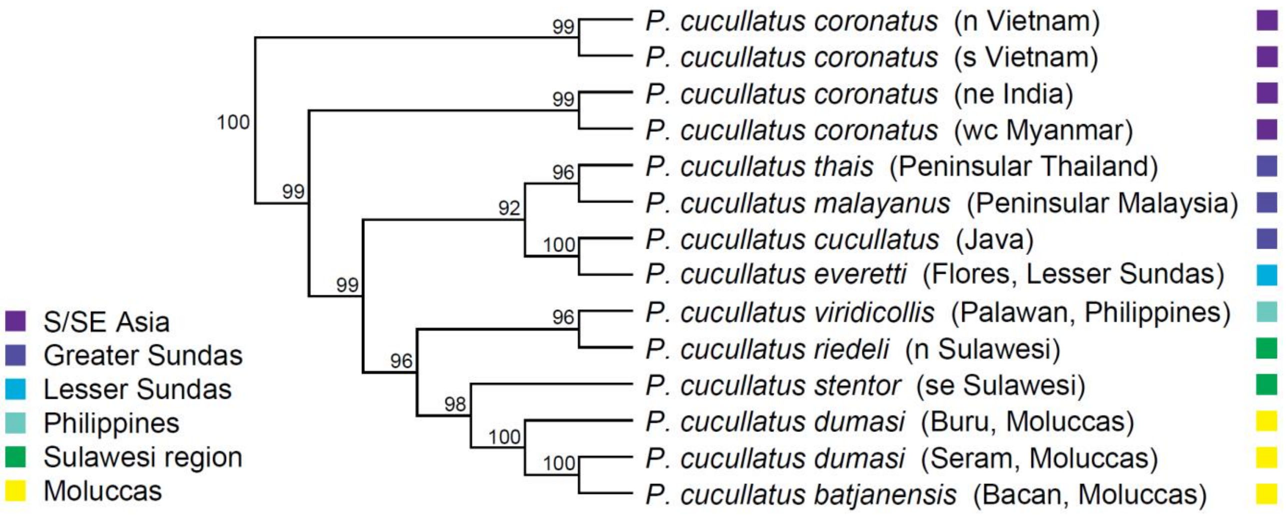


**Fig. S14.** Mountain Tailorbird *Phyllergates cucullatus*: Concatenated genomic maximum likelihood tree. ML bootstrap supports ≥ 50 are indicated at the nodes. Outgroup taxa are not shown. Colored cells indicate areas of occurrence (breeding ranges).

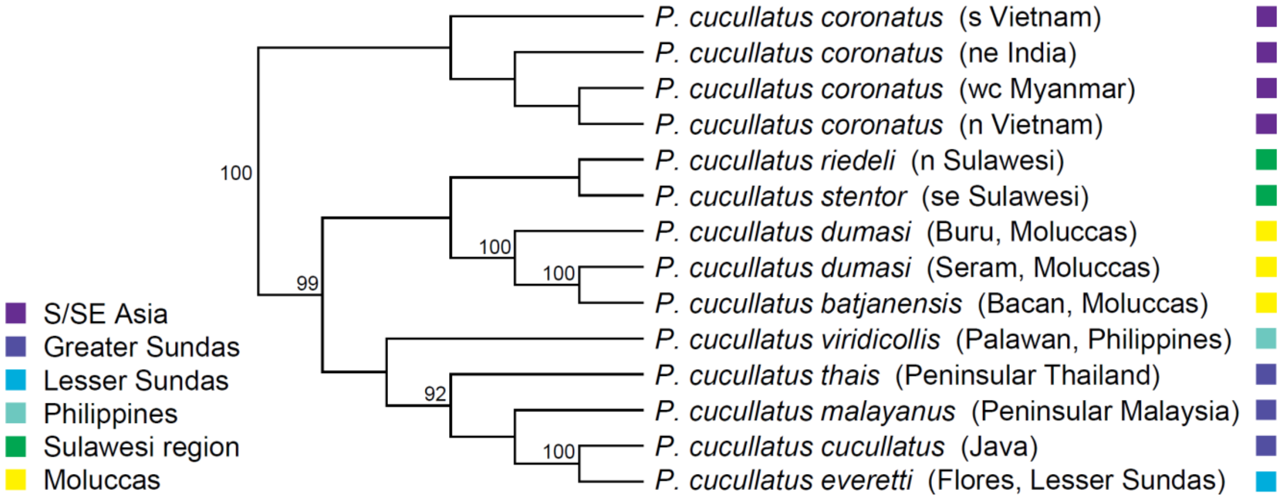


**Fig. S15.** Mountain Tailorbird *Phyllergates cucullatus*: Genomic species tree with maximum likelihood bootstrap values. ML bootstrap supports ≥ 50 are indicated at the nodes. Outgroup taxa are not shown. Colored cells indicate areas of occurrence (breeding ranges).

**Table S1.** Taxonomic diversity of species with montane island population (MIPs).

| | Genera | Families |
|---|---|---|
| Groups identified | 47 | 22 |
| Groups placed in our trees | 37 | 17 |
| representation by Eurasian-origin species | 11 | 7 |
| representation by Australo-Papuan-origin species | 14 | 5 |
| representation by other species | 14 | 11 |

**Table S2.** Models used for ancestral state reconstructions. AIC values are shown for DEC and DECj models for the analyses of geographic range, elevational range, and migration across all clades. The DECj model was used if its AIC value was > 2 under the AIC value of the DEC model. A "-" in a cell indicates that no analysis was performed because all species had the same character state.

| Clade | Geographic Range | | Elevational Range | | Migration | |
|---|---|---|---|---|---|---|
| | DEC | DECj | DEC | DECj | DEC | DECj |
| Meliphagidae | 475.50 | 382.10 | 311.30 | 258.30 | 203.90 | 205.90 |
| Campephagidae | 412.90 | 713.10 | 142.90 | 120.80 | 102.70 | 104.70 |
| Pachycephalidae | 258.20 | 229.50 | 127.63 | 107.78 | 35.85 | 37.85 |
| Rhipiduridae A (*Rhipidura*) | 87.83 | 77.48 | 36.51 | 34.91 | 28.34 | 30.34 |
| Rhipiduridae B (Lamproliidae) | 15.12 | 11.72 | 11.17 | 9.66 | - | - |
| Corvidae | 141.90 | 136.00 | 16.97 | 15.82 | 74.83 | 75.70 |
| Petroicidae A (*Microeca*) | 21.08 | 21.06 | 21.88 | 17.37 | - | - |
| Petroicidae B (*Petroica*) | 37.25 | 34.80 | 28.88 | 26.88 | 30.38 | 32.38 |
| Stenostiridae | 37.59 | 39.56 | 21.66 | 23.66 | 15.30 | 17.30 |
| Pnoepygidae | 14.84 | 16.77 | - | - | 7.64 | 9.64 |
| Cettiidae | 190.30 | 175.30 | 78.28 | 67.01 | 84.43 | 86.43 |
| Phylloscopidae | 145.61 | 151.80 | 56.82 | 59.08 | 96.53 | 94.53 |
| Locustellidae | 266.60 | 650.80 | 124.90 | 99.71 | 156.00 | 153.00 |
| Sturnidae | 146.90 | 133.10 | 40.49 | 35.28 | 26.34 | 28.34 |
| Turdidae A (*Geokichla*) | 35.62 | 36.99 | 34.81 | 25.84 | 16.88 | 18.88 |
| Turdidae B (*Zoothera*) | 116.20 | 108.70 | 34.30 | 36.30 | 62.55 | 121.40 |
| Turdidae C (*Turdus*) | 55.14 | 57.14 | 60.28 | 55.30 | 103.90 | 105.90 |
| Muscicapidae A (*Eumyias*) | 32.31 | 31.12 | 16.28 | 17.49 | 21.56 | 23.56 |
| Muscicapidae B (*Brachypteryx*) | 46.11 | 60.50 | - | - | 15.66 | 20.27 |
| Muscicapidae C (*Ficedula*) | 103.20 | 101.80 | 87.96 | 67.18 | 79.58 | 79.75 |
| Dicaeidae | 102.26 | 88.08 | 31.48 | 27.05 | 23.39 | 25.39 |
| Motacillidae | 41.03 | 34.93 | 26.04 | 26.95 | 44.80 | 43.19 |
| Fringillidae | 54.73 | 54.26 | 70.78 | 61.75 | 122.30 | 124.30 |

**Table S3.** Establishment of multiple montane island populations (MIPs) by Eurasian-origin vs. Australo-Papuan-origin species.

| | All archipelagos | | Wallacea | | Bismarcks/Solomons | |
|---|---|---|---|---|---|---|
| No. MIPs | mean | median | mean | median | mean | median |
| Eurasian-origin species | 3.23 | 2 | 3.29 | 2 | 2.63 | 2.5 |
| Australo-Papuan-origin species | 1.32 | 1 | 1.55 | 1 | 1.14 | 1 |
| Proportion of spp with >1 MIP | | | | | | |
| Eurasian-origin species | 0.65 | | 0.67 | | 0.63 | |
| Australo-Papuan-origin species | 0.2 | | 0.36 | | 0.07 | |

**Table S4.** Statistical testing of differences in Table S3: multiple MIP establishment by Eurasian-origin vs. Australo-Papuan-origin species. Results of Mann-Whitney U-tests and chi-squared tests are given. Tests were repeated across different regions: all archipelagos (Wal/Bis/Sol); Wallacea (Wal); and the Bismarcks and Solomons (Bis/Sol).

| **Measure** | **Region** | **Test** | **W** | **Chi-sq** | **df** | **p** |
|---|---|---|---|---|---|---|
| No. MIPs | Wal/Bis/Sol | M-W U-Test | 581.5 | NA | NA | 0.0004472 |
| | Wal | | 188 | | | 0.0385 |
| | Bis/Sol | | 87.5 | | | 0.00692 |
| Frequency of spp with >1 MIPs | Wal/Bis/Sol | Chi-sq Test | NA | 9.3692 | 1 | 0.002207 |
| | Wal | | | 1.7262 | 1 | 0.1889 |
| | Bis/Sol | | | 5.322 | 1 | 0.02106 |

**Table S5.** Results for comparisons of empirical vs phylogenetic null datasets. Abbreviations: MIPs = montane island populations; IPs = total island populations; EUR = Eurasian-origin species; AP = Australo-Papuan-origin species; SDM = short-distance migration; SED = sedentary; CON = continental occurrence; ISL = no continental occurrence. Numbers in parentheses refer to the methods and results sections where the tests of empirical data and the results are described. F statistics from one-way ANOVA tests of empirical data are given, as well as the 5% upper quantiles of the distributions of the simulated F statistics.

| Analysis | Empirical Value of F | Null upper 5% | p |
|---|---|---|---|
| Montane vs. lowland ancestry: EUR vs. AP (2.4.2, 4.4.2) | 107.1 | 36.91108 | 0.001 |
| No. MIPs (total): EUR vs. AP (2.4.3, 4.4.3) | 11 | 14.37757 | 0.071 |
| No. MIPs (Wallacea): EUR vs. AP (2.4.3, 4.4.3) | 4.224 | 8.359259 | 0.158 |
| No. MIPs (Bismarcks/Solomons): EUR vs. AP (2.4.3, 4.4.3) | 8.741 | 11.51451 | 0.076 |
| MIPs/IPs: EUR vs. AP (2.4.4, 4.4.4) | 25.96 | 14.82485 | 0.012 |
| Migration ancestry: EUR vs. AP (2.4.5, 4.4.5) | 49.01 | 40.22154 | 0.029 |
| No. MIPs (total): EUR SDM vs. SED (2.4.5, 4.4.5) | 7.866 | 5.979078 | 0.024 |
| No. MIPs (total): EUR CON vs. ISL (2.4.6, 4.4.6) | 9.761 | 7.742936 | 0.025 |

**Table S6.** Taxon sampling of montane supercolonizers.

| Clade | No. taxa | No. taxa our sampling | No. individuals our sampling | No. taxa Genbank | No. individuals Genbank | No. taxa supermatrix | No. individuals supermatrix |
|---|---|---|---|---|---|---|---|
| Indo-Pacific *Phylloscopus* | 38 | 26 (68%) | 30 | 16 | 23 | 37 (97%) | 53 |
| *Phylloscopus amoenus* | 1 | 0 (0%) | 0 | 1 | 1 | 1 (100%) | 1 |
| *Phylloscopus maforensis* | 20 | 16 (80%) | 18 | 4 | 6 | 20 (100%) | 24 |
| *Phylloscopus makirensis* | 1 | 1 (100%) | 1 | 1 | 1 | 1 (100%) | 2 |
| *Phylloscopus nigrorum* | 7 | 3 (43%) | 3 | 5 | 8 | 7 (100%) | 11 |
| *Phylloscopus presbytes* | 2 | 2 (100%) | 2 | 1 | 1 | 2 (100%) | 3 |
| *Phylloscopus rotiensis* | 1 | 0 (0%) | 0 | 0 | 0 | 0 (0%) | 0 |
| *Phylloscopus sarasinorum* | 2 | 1 (50%) | 1 | 2 | 3 | 2 (100%) | 4 |
| *Phylloscopus trivirgatus* | 4 | 3 (75%) | 5 | 2 | 3 | 4 (100%) | 8 |
| *Ficedula hyperythra* | 14 | 9 (64%) | 12 | 6 | 26 | 13 (93%) | 38 |
| *Phyllergates cucullatus* | 16 | 11 (69%) | 15 | 8 | 15 | 16 (100%) | 30 |

**Table S7.** Reference sequences for gene extraction.

| Study taxon | Reference sequences |
|---|---|
| *Phylloscopus* spp | mitochondrial genome: *Phylloscopus examinandus* LR026996 |
| | Myo: *Phylloscopus claudiae* MH079093 |
| | GAPDH: *Phylloscopus claudiae* MH042994 |
| | ODC: *Phylloscopus claudiae* MH037419 |
| *Ficedula* spp | mitochondrial genome: *Ficedula zanthopygia* JN018411 |
| | Myo: *Ficedula tricolor* KJ931278 |
| | ODC: *Ficedula tricolor* KJ931305 |
| | PEPCK: *Ficedula mugimaki* KJ931247 |
| *Phyllergates cucullatus* | mitochondrial genome: *Aegithalos glaucogularis* KF951090 |
| | Myo: *Tickellia hodgsoni* DQ008565 |
| | GAPDH: *Tickellia hodgsoni* HQ121538 |
| | ODC: *Tickellia hodgsoni* EU680774 |
| | TGFb2: *Phyllergates heterolaemus* JX006177 |
| | MUSK: *Phyllergates heterolaemus* JX006218 |
| *Horornis parens* | mitochondrial genome: *Horonis fortipes* MK051002 |
| *Poodytes albolimbatus* | mitochondrial genome: *Poodytes punctatus* NC_029138 |
| *Aplonis mystacea* | mitochondrial genome: *Acridotheres tristis* HQ915864 |
| *Dicaeum pectorale* | mitochondrial genome: *Aethopyga gouldiae* NC_027241 |
