## Supplementary File 3 for "The formation of the Indo-Pacific montane avifauna"

**Supplementary File 3.** Species-level trees. Black circles at nodes indicate posterior probabilities of 0.99-1.00; gray circles indicate PP 0.95-0.98; and white circles indicate PP 0.90-0.94. Nodes without circles have PP < 0.90.

**Contents:**

2. Pachycephalidae - p. 2
3. Petroicidae A (*Microeca*) - p. 3
4. Petroicidae B (*Petroica*) - p. 4
5. Stenostiridae - p. 5
6. Pnoepygidae - p. 6
7. Cettiidae - p. 7
8. Locustellidae - p. 8
9. Sturnidae - p. 9
10. Turdidae A (*Geokichla*) - p. 10
11. Turdidae B (*Zoothera*) - p. 11
12. Turdidae C (*Turdus*) - p. 12
13. Muscicapidae A (*Eumyias*) - p. 13
14. Muscicapidae B (*Brachypteryx*) - p. 14
15. Muscicapidae C (*Ficedula*) - p. 15
16. Dicaeidae - p. 16
17. Fringillidae - p. 17

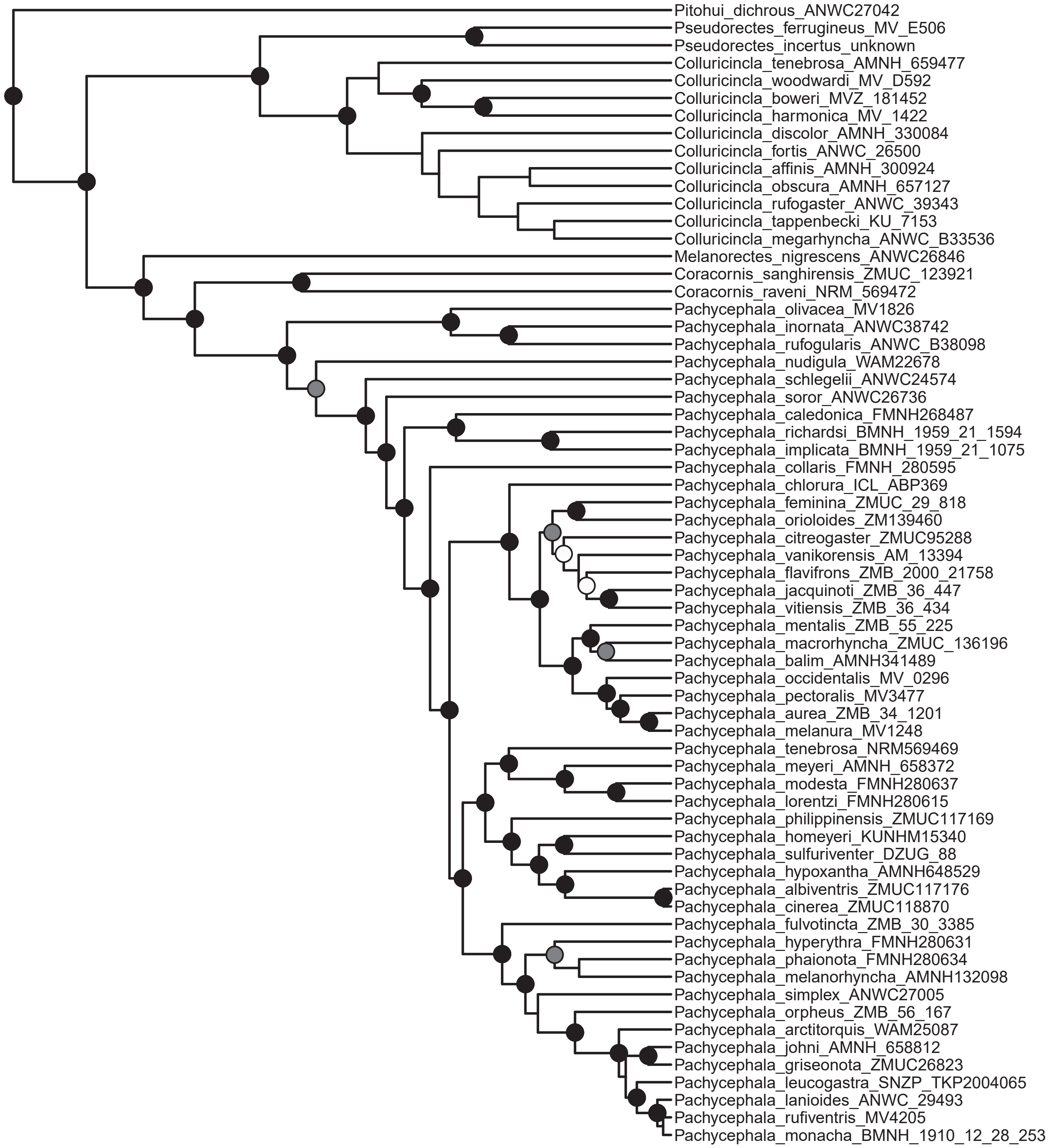
Pitohui_dichrous_ANWC27042
Pseudorectes_ferrugineus_MV_E506
Pseudorectes_incertus_unknown
Colluricincla_tenebrosa_AMNH_659477
Colluricincla_woodwardi_MV_D592
Colluricincla_boweri_MVZ_181452
Colluricincla_harmonica_MV_1422
Colluricincla_discolor_AMNH_330084
Colluricincla_fortis_ANWC_26500
Colluricincla_affinis_AMNH_300924
Colluricincla_obscura_AMNH_657127
Colluricincla_rufogaster_ANWC_39343
Colluricincla_tappenbecki_KU_7153
Colluricincla_megarhyncha_ANWC_B33536
Melanorectes_nigrescens_ANWC26846
Coracornis_sanghirensis_ZMUC_123921
Coracornis_raveni_NRM_569472
Pachycephala_olivacea_MV1826
Pachycephala_inornata_ANWC38742
Pachycephala_rufogularis_ANWC_B38098
Pachycephala_nudigula_WAM22678
Pachycephala_schlegelii_ANWC24574
Pachycephala_soror_ANWC26736
Pachycephala_caledonica_FMNH268487
Pachycephala_richardsi_BMNH_1959_21_1594
Pachycephala_implicata_BMNH_1959_21_1075
Pachycephala_collaris_FMNH_280595
Pachycephala_chlorura_ICL_ABP369
Pachycephala_feminina_ZMUC_29_818
Pachycephala_orioloides_ZM139460
Pachycephala_citreogaster_ZMUC95288
Pachycephala_vanikorensis_AM_13394
Pachycephala_flavifrons_ZMB_2000_21758
Pachycephala_jacquinoti_ZMB_36_447
Pachycephala_vitiensis_ZMB_36_434
Pachycephala_mentalis_ZMB_55_225
Pachycephala_macrorhyncha_ZMUC_136196
Pachycephala_balim_AMNH341489
Pachycephala_occidentalis_MV_0296
Pachycephala_pectoralis_MV3477
Pachycephala_aurea_ZMB_34_1201
Pachycephala_melanura_MV1248
Pachycephala_tenebrosa_NRM569469
Pachycephala_meyeri_AMNH_658372
Pachycephala_modesta_FMNH280637
Pachycephala_lorentzi_FMNH280615
Pachycephala_philippinensis_ZMUC117169
Pachycephala_homeyeri_KUNHM15340
Pachycephala_sulfuriventer_DZUG_88
Pachycephala_hypoxantha_AMNH648529
Pachycephala_albiventris_ZMUC117176
Pachycephala_cinerea_ZMUC118870
Pachycephala_fulvotincta_ZMB_30_3385
Pachycephala_hyperythra_FMNH280631
Pachycephala_phaionota_FMNH280634
Pachycephala_melanorhyncha_AMNH132098
Pachycephala_simplex_ANWC27005
Pachycephala_orpheus_ZMB_56_167
Pachycephala_arctitorquis_WAM25087
Pachycephala_johni_AMNH_658812
Pachycephala_griseonota_ZMUC26823
Pachycephala_leucogastra_SNZP_TKP2004065
Pachycephala_lanioides_ANWC_29493
Pachycephala_rufiventris_MV4205
Pachycephala_monacha_BMNH_1910_12_28_253

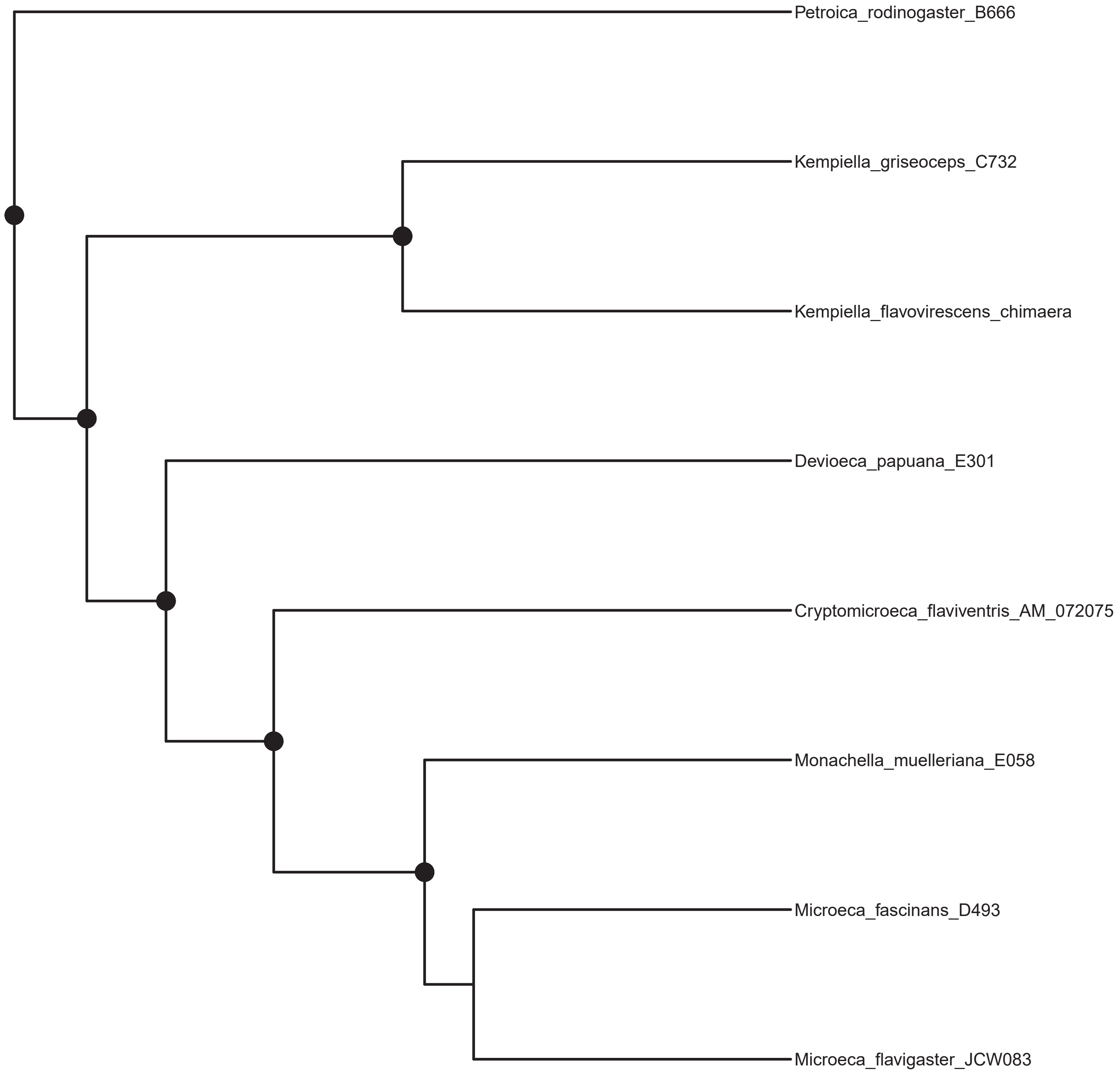
Petroica_rodinogaster_B666
Kempiella_griseoceps_C732
Kempiella_flavovirescens_chimaera
Devioeca_papuana_E301
Cryptomicroeca_flaviventris_AM_072075
Monachella_muelleriana_E058
Microeca_fascinans_D493
Microeca_flavigaster_JCW083

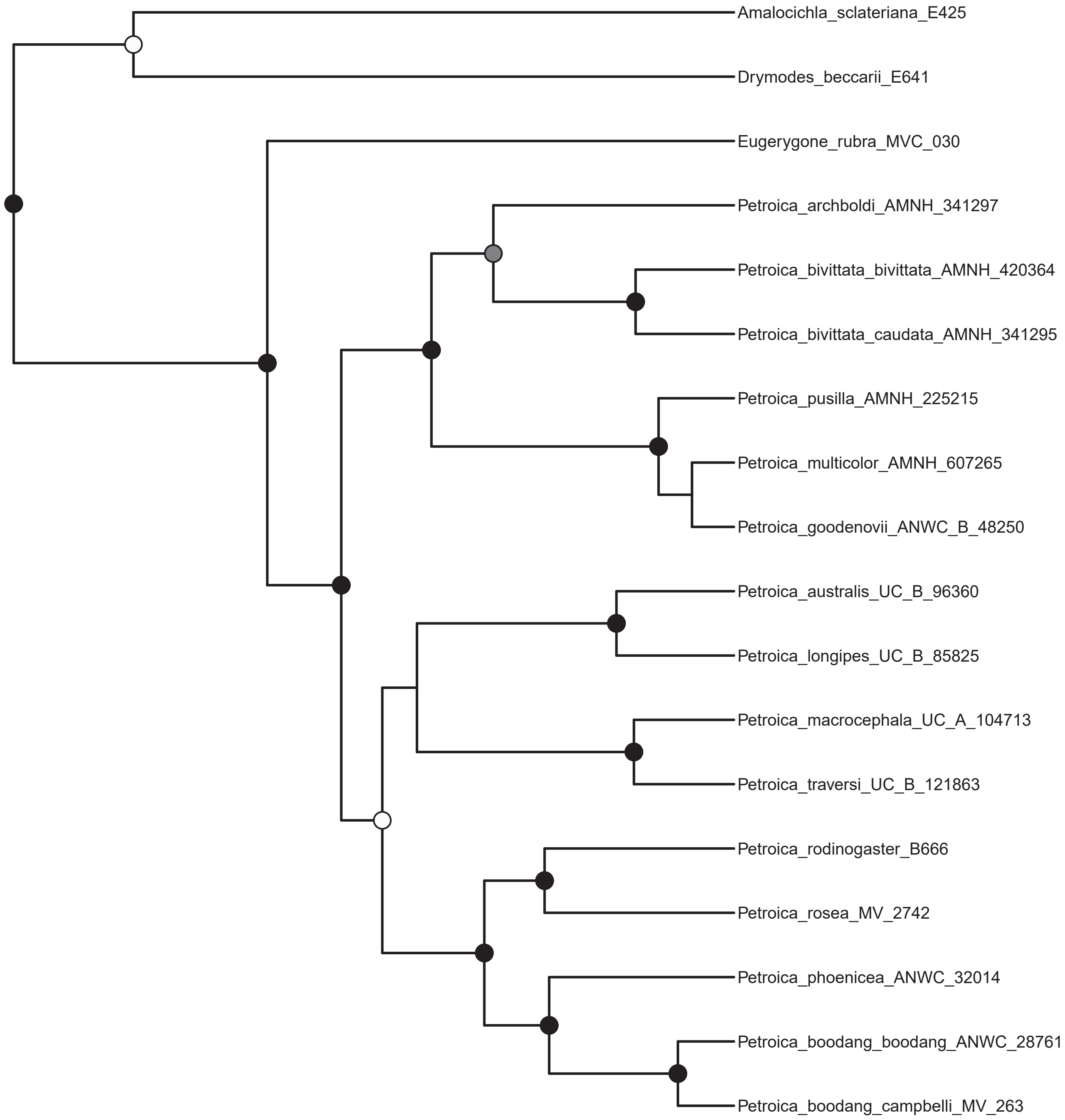
Amalocichla_sclateriana_E425
Drymodes_beccarii_E641
Eugerygone_rubra_MVC_030
Petroica_archboldi_AMNH_341297
Petroica_bivittata_bivittata_AMNH_420364
Petroica_bivittata_caudata_AMNH_341295
Petroica_pusilla_AMNH_225215
Petroica_multicolor_AMNH_607265
Petroica_goodenovii_ANWC_B_48250
Petroica_australis_UC_B_96360
Petroica_longipes_UC_B_85825
Petroica_macrocephala_UC_A_104713
Petroica_traversi_UC_B_121863
Petroica_rodinogaster_B666
Petroica_rosea_MV_2742
Petroica_phoenicea_ANWC_32014
Petroica_boodang_boodang_ANWC_28761
Petroica_boodang_campbelli_MV_263

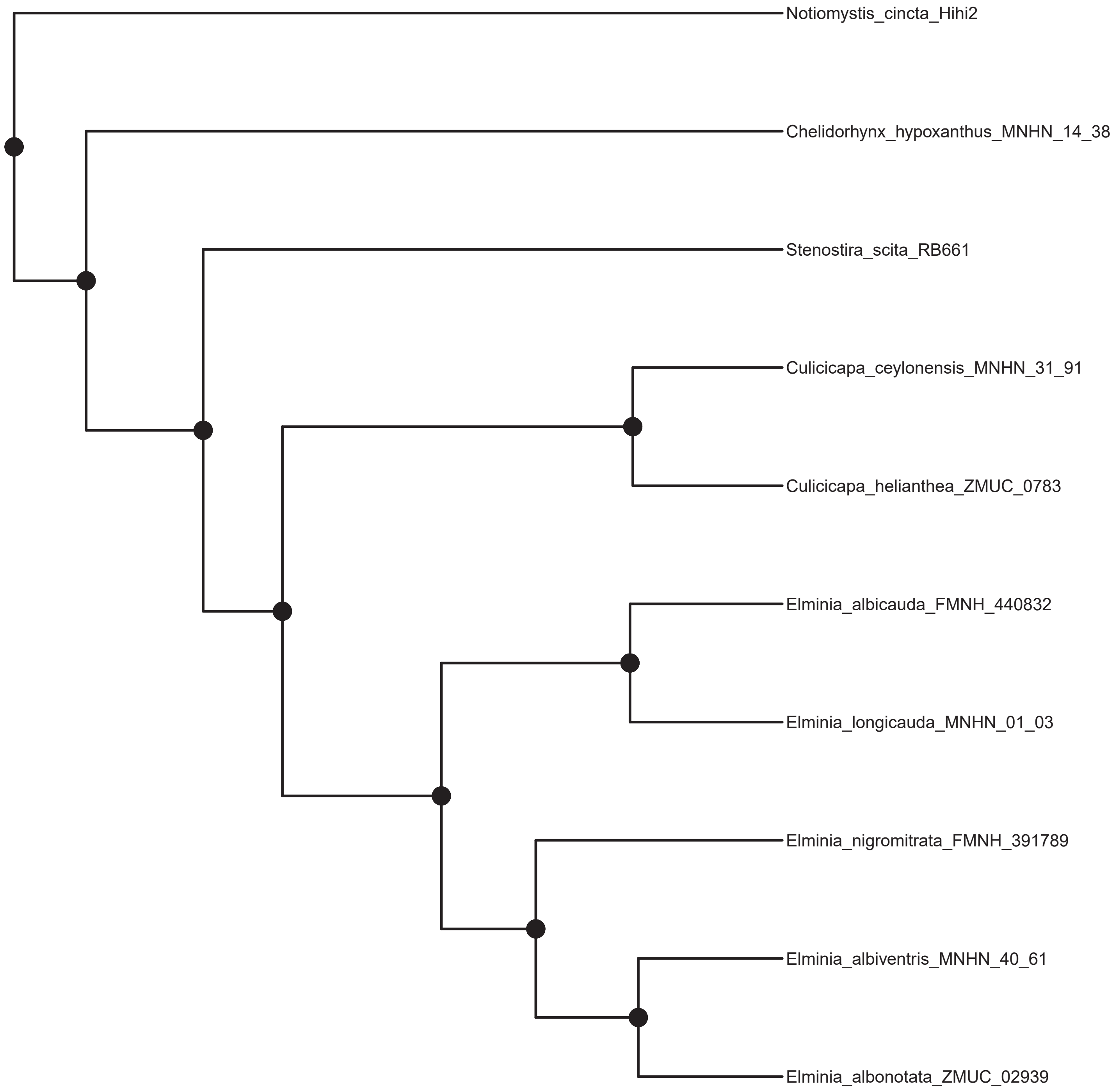
Notiomystis_cincta_Hihi2
Chelidorhynx_hypoxanthus_MNHN_14_38
Stenostira_scita_RB661
Culicicapa_ceylonensis_MNHN_31_91
Culicicapa_helianthea_ZMUC_0783
Elminia_albicauda_FMNH_440832
Elminia_longicauda_MNHN_01_03
Elminia_nigromitrata_FMNH_391789
Elminia_albiventris_MNHN_40_61
Elminia_albonotata_ZMUC_02939

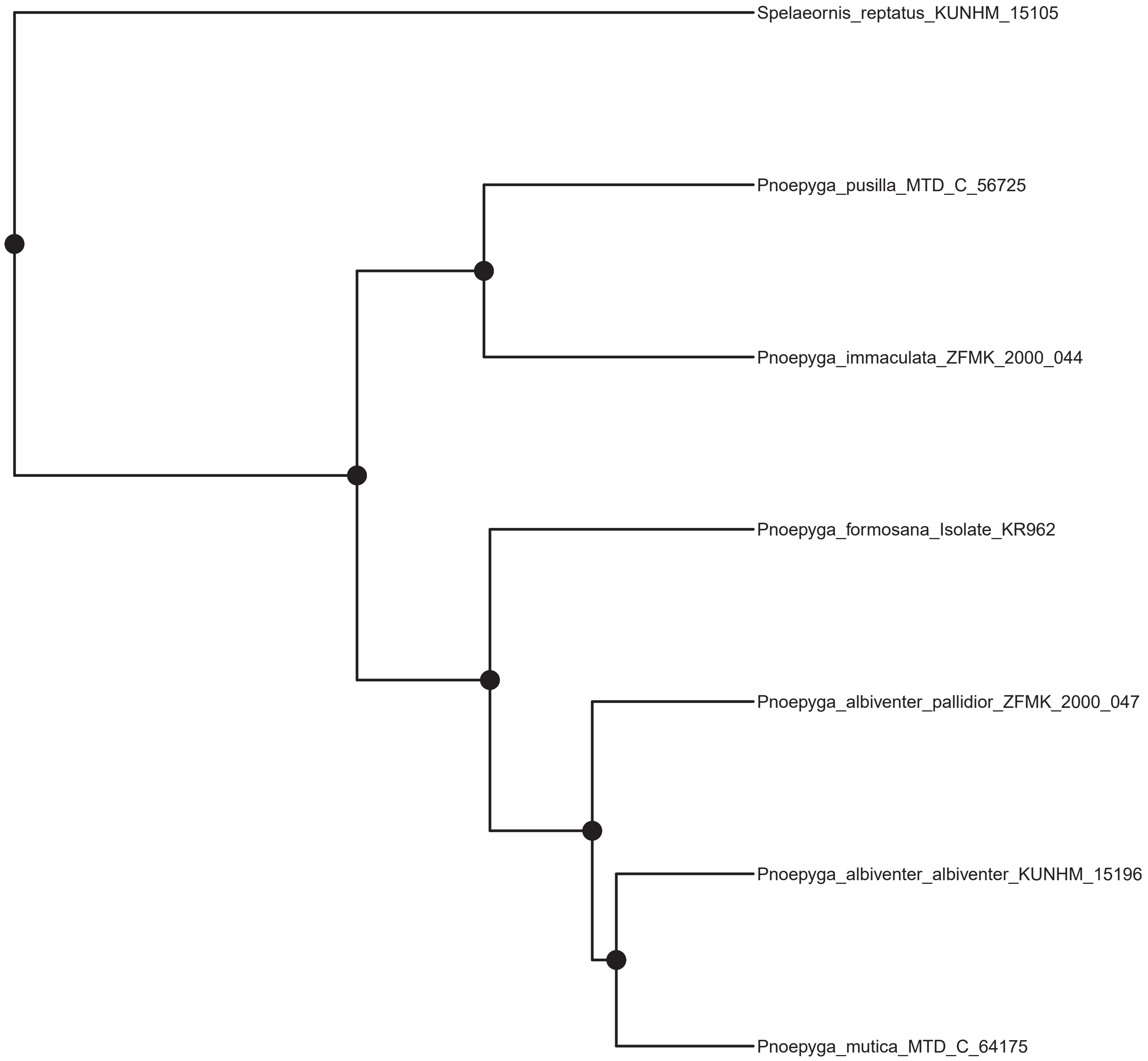
Spelaeornis_reptatus_KUNHM_15105
Pnoepyga_pusilla_MTD_C_56725
Pnoepyga_immaculata_ZFMK_2000_044
Pnoepyga_formosana_Isolate_KR962
Pnoepyga_albiventer_pallidior_ZFMK_2000_047
Pnoepyga_albiventer_albiventer_KUNHM_15196
Pnoepyga_mutica_MTD_C_64175

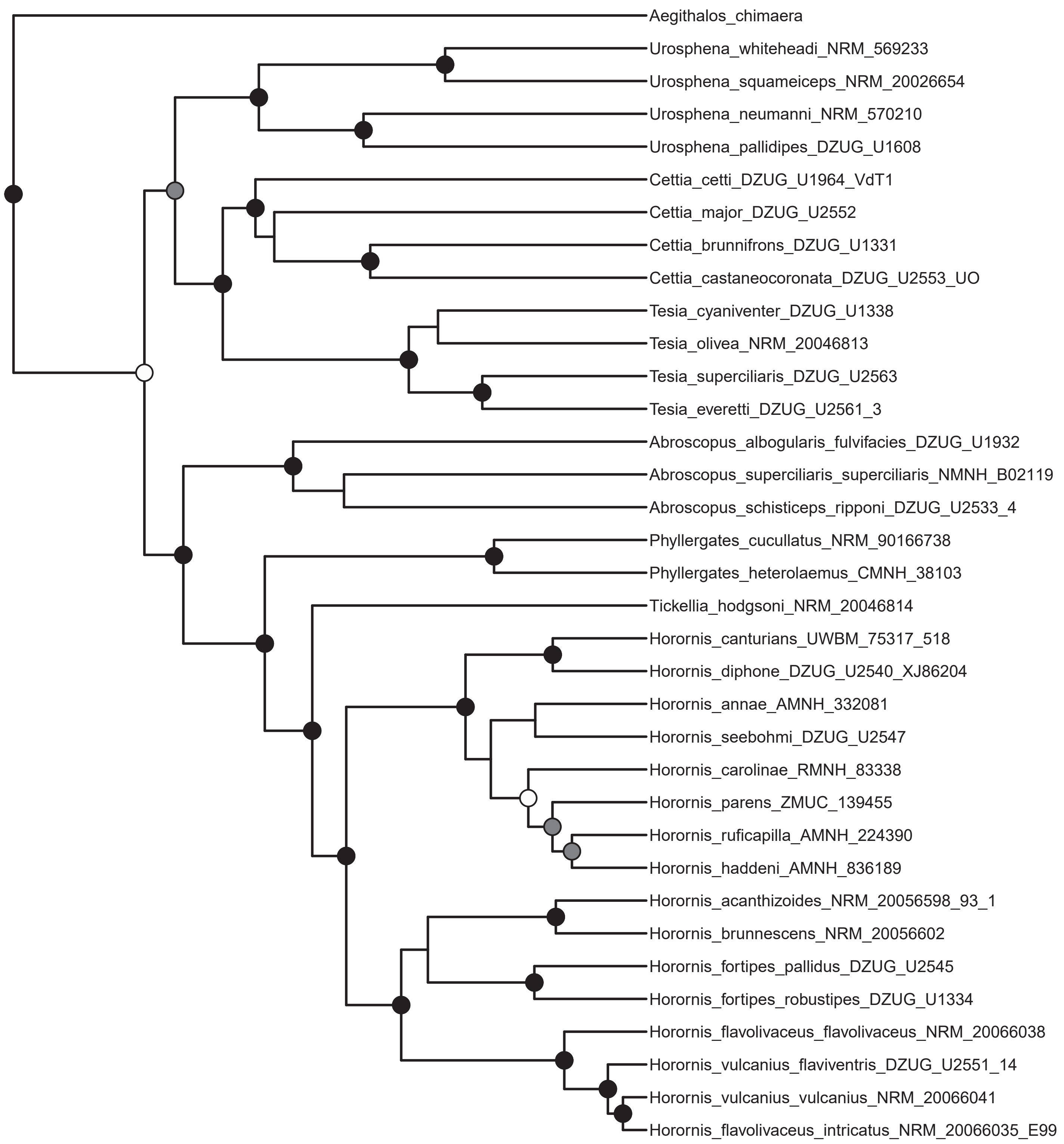
Aegithalos_chimaera
Urosphena_whiteheadi_NRM_569233
Urosphena_squameiceps_NRM_20026654
Urosphena_neumanni_NRM_570210
Urosphena_pallidipes_DZUG_U1608
Cettia_cetti_DZUG_U1964_VdT1
Cettia_major_DZUG_U2552
Cettia_brunnifrons_DZUG_U1331
Cettia_castaneocoronata_DZUG_U2553_UO
Tesia_cyaniventer_DZUG_U1338
Tesia_olivea_NRM_20046813
Tesia_superciliaris_DZUG_U2563
Tesia_everetti_DZUG_U2561_3
Abroscopus_albogularis_fulvifacies_DZUG_U1932
Abroscopus_superciliaris_superciliaris_NMNH_B02119
Abroscopus_schisticeps_ripponi_DZUG_U2533_4
Phyllergates_cucullatus_NRM_90166738
Phyllergates_heterolaemus_CMNH_38103
Tickellia_hodgsoni_NRM_20046814
Horornis_canturians_UWBM_75317_518
Horornis_diphone_DZUG_U2540_XJ86204
Horornis_annae_AMNH_332081
Horornis_seebohmi_DZUG_U2547
Horornis_carolinae_RMNH_83338
Horornis_parens_ZMUC_139455
Horornis_ruficapilla_AMNH_224390
Horornis_haddeni_AMNH_836189
Horornis_acanthizoides_NRM_20056598_93_1
Horornis_brunnescens_NRM_20056602
Horornis_fortipes_pallidus_DZUG_U2545
Horornis_fortipes_robustipes_DZUG_U1334
Horornis_flavolivaceus_flavolivaceus_NRM_20066038
Horornis_vulcanius_flaviventris_DZUG_U2551_14
Horornis_vulcanius_vulcanius_NRM_20066041
Horornis_flavolivaceus_intricatus_NRM_20066035_E99

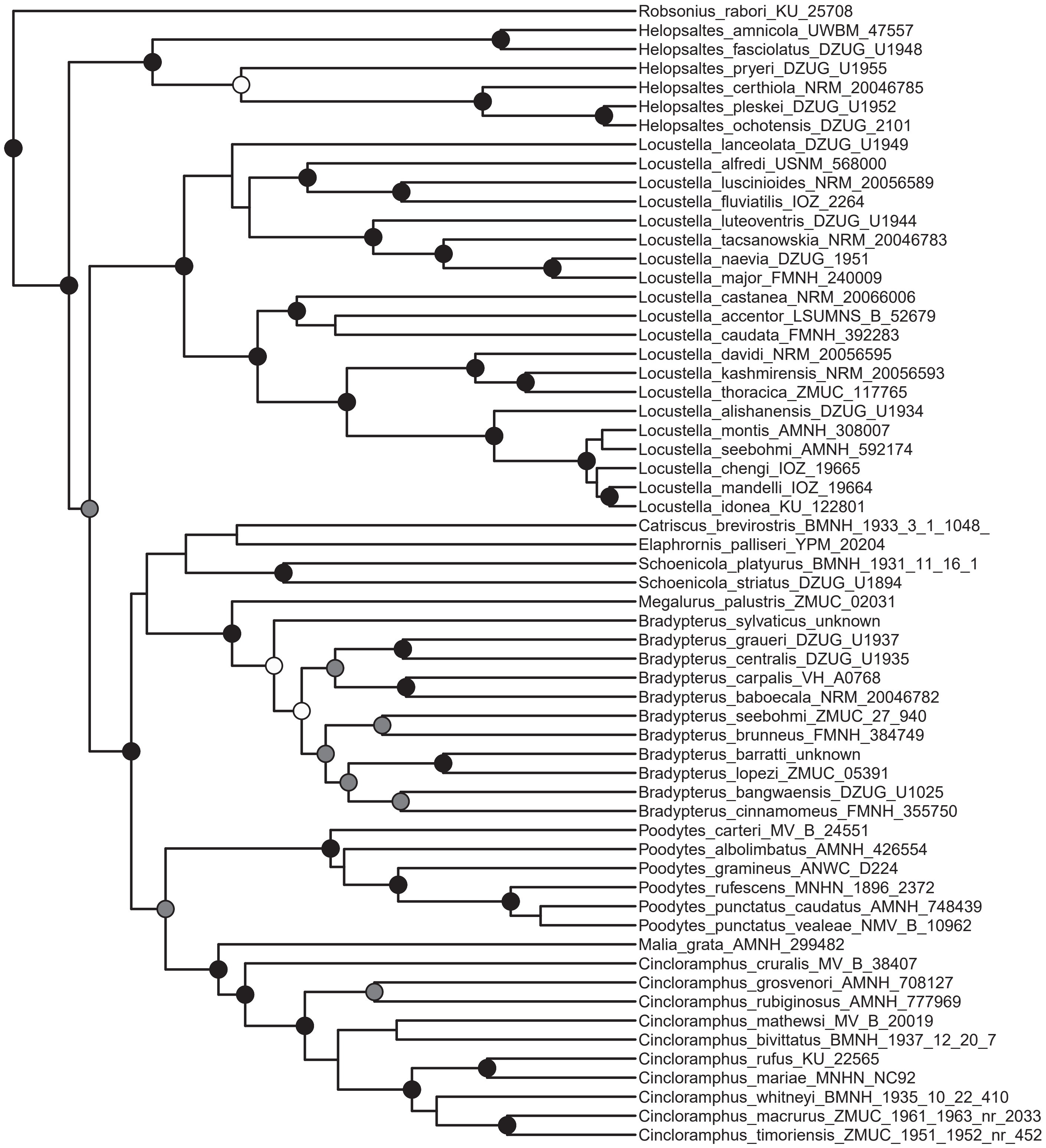
Robsonius_rabori_KU_25708
Helopsaltes_amnicola_UWBM_47557
Helopsaltes_fasciolatus_DZUG_U1948
Helopsaltes_pryeri_DZUG_U1955
Helopsaltes_certhiola_NRM_20046785
Helopsaltes_pleskei_DZUG_U1952
Helopsaltes_ochotensis_DZUG_2101
Locustella_lanceolata_DZUG_U1949
Locustella_alfredi_USNM_568000
Locustella_luscinioides_NRM_20056589
Locustella_fluviatilis_IOZ_2264
Locustella_luteoventris_DZUG_U1944
Locustella_tacsanowskia_NRM_20046783
Locustella_naevia_DZUG_1951
Locustella_major_FMNH_240009
Locustella_castanea_NRM_20066006
Locustella_accentor_LSUMNS_B_52679
Locustella_caudata_FMNH_392283
Locustella_davidi_NRM_20056595
Locustella_kashmirensis_NRM_20056593
Locustella_thoracica_ZMUC_117765
Locustella_alishanensis_DZUG_U1934
Locustella_montis_AMNH_308007
Locustella_seebohmi_AMNH_592174
Locustella_chengi_IOZ_19665
Locustella_mandelli_IOZ_19664
Locustella_idonea_KU_122801
Catriscus_brevirostris_BMNH_1933_3_1_1048_
Elaphrornis_palliseri_YPM_20204
Schoenicola_platyurus_BMNH_1931_11_16_1
Schoenicola_striatus_DZUG_U1894
Megalurus_palustris_ZMUC_02031
Bradypterus_sylvaticus_unknown
Bradypterus_graueri_DZUG_U1937
Bradypterus_centralis_DZUG_U1935
Bradypterus_carpalis_VH_A0768
Bradypterus_baboecala_NRM_20046782
Bradypterus_seebohmi_ZMUC_27_940
Bradypterus_brunneus_FMNH_384749
Bradypterus_barratti_unknown
Bradypterus_lopezi_ZMUC_05391
Bradypterus_bangwaensis_DZUG_U1025
Bradypterus_cinnamomeus_FMNH_355750
Poodytes_carteri_MV_B_24551
Poodytes_albolimbatus_AMNH_426554
Poodytes_gramineus_ANWC_D224
Poodytes_rufescens_MNHN_1896_2372
Poodytes_punctatus_caudatus_AMNH_748439
Poodytes_punctatus_vealeae_NMV_B_10962
Malia_grata_AMNH_299482
Cincloramphus_cruralis_MV_B_38407
Cincloramphus_grosvenori_AMNH_708127
Cincloramphus_rubiginosus_AMNH_777969
Cincloramphus_mathewsi_MV_B_20019
Cincloramphus_bivittatus_BMNH_1937_12_20_7
Cincloramphus_rufus_KU_22565
Cincloramphus_mariae_MNHN_NC92
Cincloramphus_whitneyi_BMNH_1935_10_22_410
Cincloramphus_macrurus_ZMUC_1961_1963_nr_2033
Cincloramphus_timoriensis_ZMUC_1951_1952_nr_452

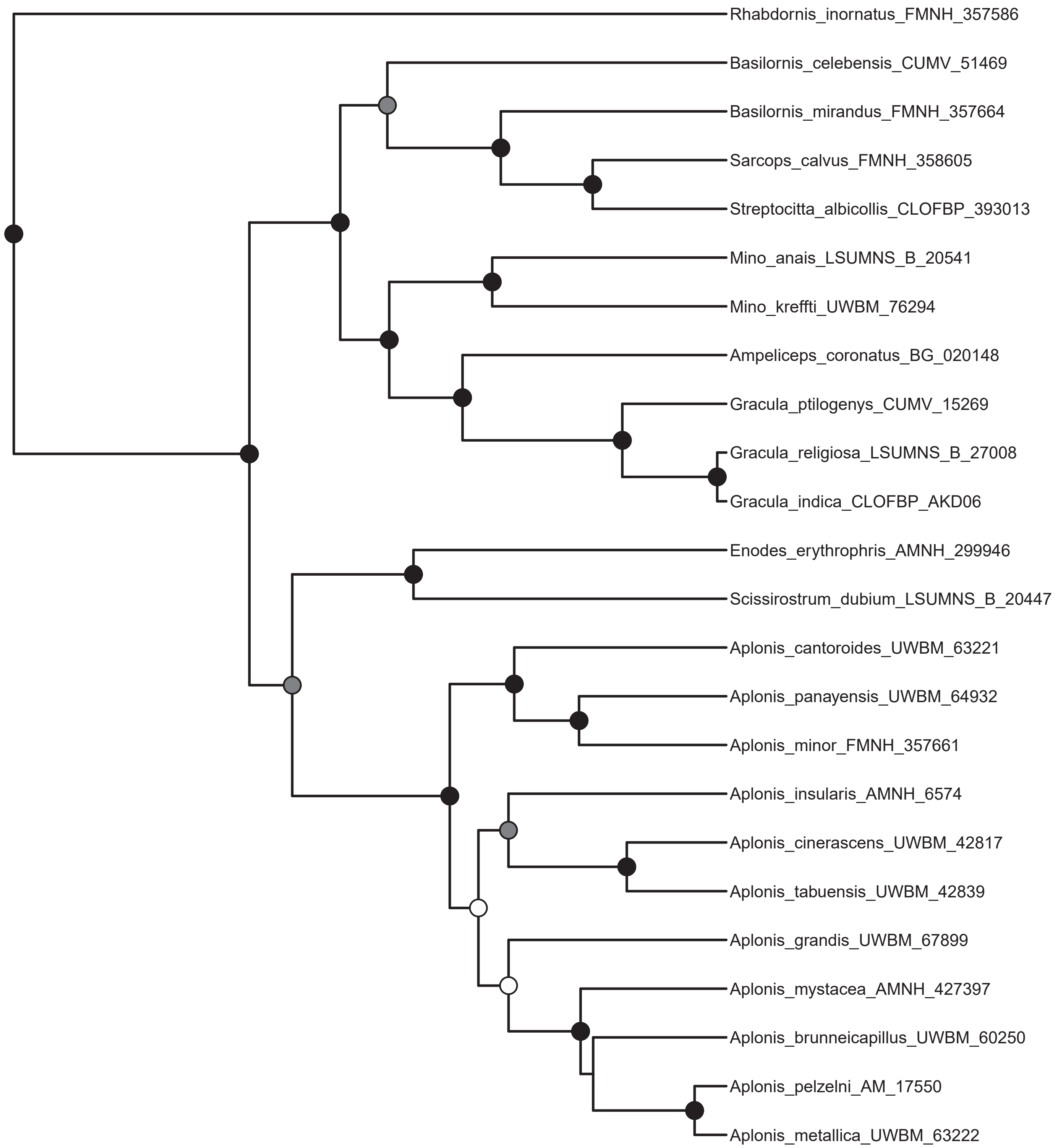
Rhabdornis_inornatus_FMNH_357586
Basilornis_celebensis_CUMV_51469
Basilornis_mirandus_FMNH_357664
Sarcops_calvus_FMNH_358605
Streptocitta_albicollis_CLOFBP_393013
Mino_anais_LSUMNS_B_20541
Mino_kreffti_UWBM_76294
Ampeliceps_coronatus_BG_020148
Gracula_ptilogenys_CUMV_15269
Gracula_religiosa_LSUMNS_B_27008
Gracula_indica_CLOFBP_AKD06
Enodes_erythrophris_AMNH_299946
Scissirostrum_dubium_LSUMNS_B_20447
Aplonis_cantoroides_UWBM_63221
Aplonis_panayensis_UWBM_64932
Aplonis_minor_FMNH_357661
Aplonis_insularis_AMNH_6574
Aplonis_cinerascens_UWBM_42817
Aplonis_tabuensis_UWBM_42839
Aplonis_grandis_UWBM_67899
Aplonis_mystacea_AMNH_427397
Aplonis_brunneicapillus_UWBM_60250
Aplonis_pelzelni_AM_17550
Aplonis_metallica_UWBM_63222

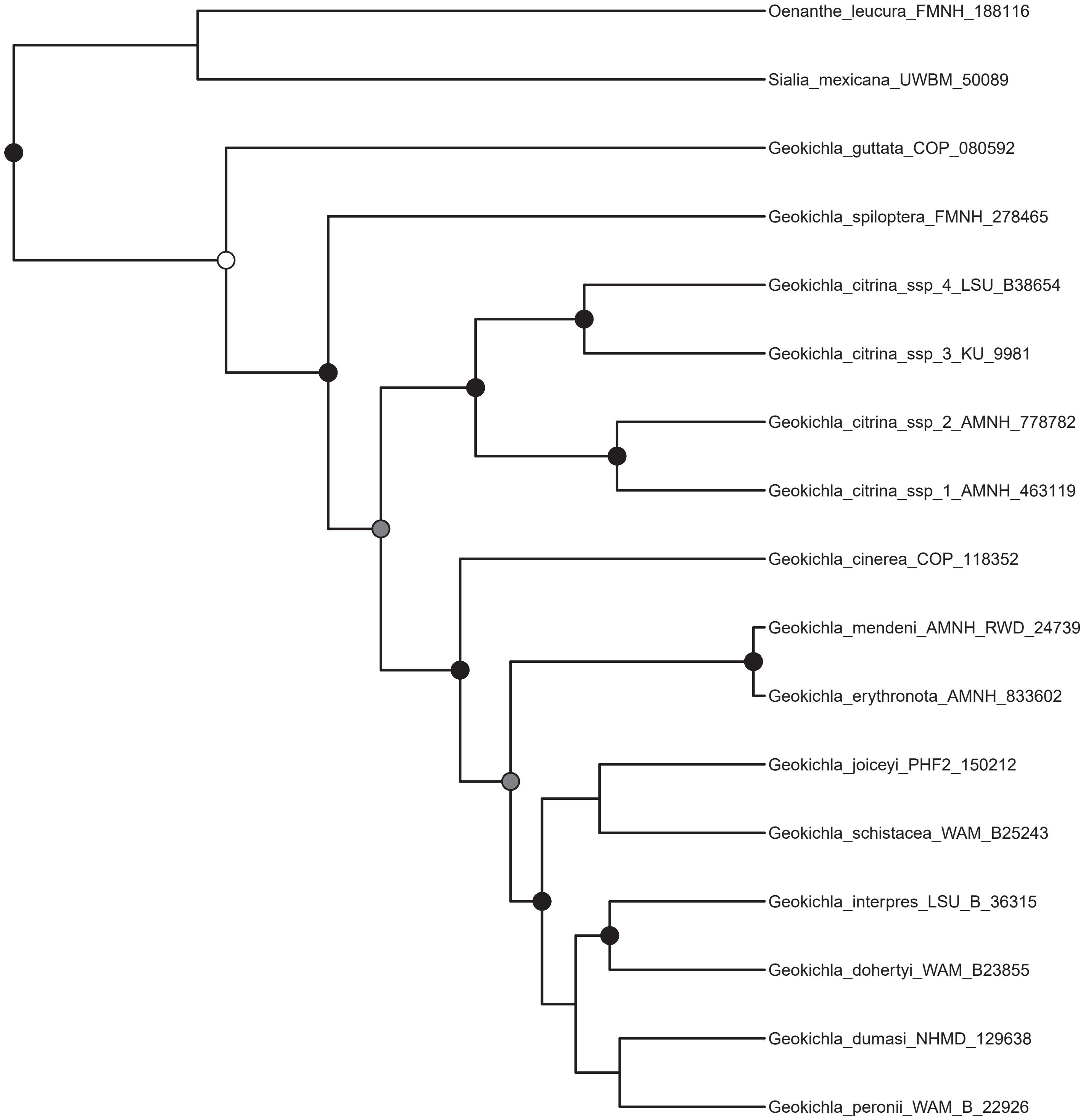
Oenanthe_leucura_FMNH_188116
Sialia_mexicana_UWBM_50089
Geokichla_guttata_COP_080592
Geokichla_spiloptera_FMNH_278465
Geokichla_citrina_ssp_4_LSU_B38654
Geokichla_citrina_ssp_3_KU_9981
Geokichla_citrina_ssp_2_AMNH_778782
Geokichla_citrina_ssp_1_AMNH_463119
Geokichla_cinerea_COP_118352
Geokichla_mendeni_AMNH_RWD_24739
Geokichla_erythronota_AMNH_833602
Geokichla_joiceyi_PHF2_150212
Geokichla_schistacea_WAM_B25243
Geokichla_interpres_LSU_B_36315
Geokichla_dohertyi_WAM_B23855
Geokichla_dumasi_NHMD_129638
Geokichla_peronii_WAM_B_22926

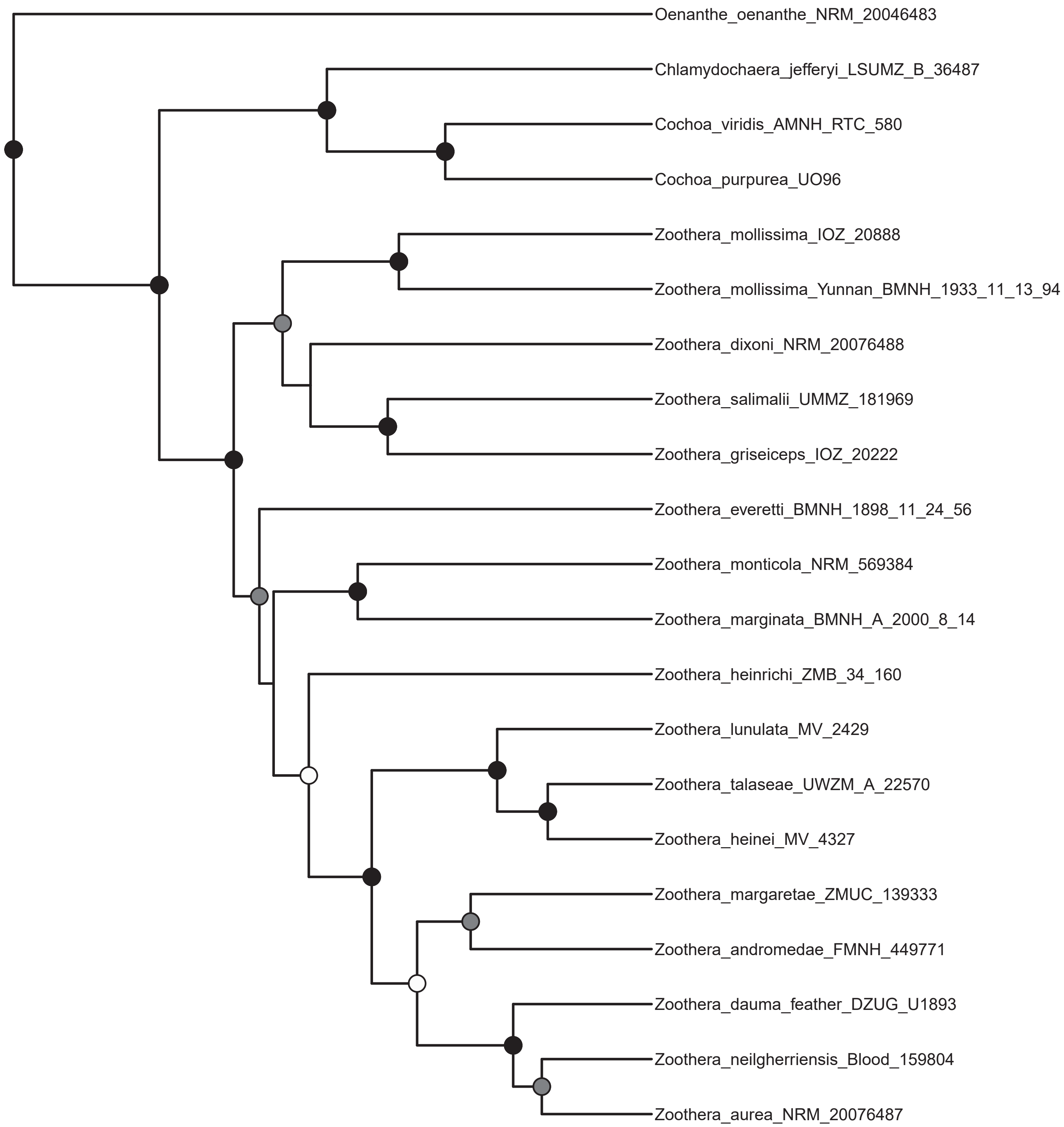
Oenanthe_oenanthe_NRM_20046483
Chlamydochaera_jefferyi_LSUMZ_B_36487
Cochoa_viridis_AMNH_RTC_580
Cochoa_purpurea_UO96
Zoothera_mollissima_IOZ_20888
Zoothera_mollissima_Yunnan_BMNH_1933_11_13_94
Zoothera_dixoni_NRM_20076488
Zoothera_salimalii_UMMZ_181969
Zoothera_griseiceps_IOZ_20222
Zoothera_everetti_BMNH_1898_11_24_56
Zoothera_monticola_NRM_569384
Zoothera_marginata_BMNH_A_2000_8_14
Zoothera_heinrichi_ZMB_34_160
Zoothera_lunulata_MV_2429
Zoothera_talaseae_UWZM_A_22570
Zoothera_heinei_MV_4327
Zoothera_margaretae_ZMUC_139333
Zoothera_andromedae_FMNH_449771
Zoothera_dauma_feather_DZUG_U1893
Zoothera_neilgherriensis_Blood_159804
Zoothera_aurea_NRM_20076487

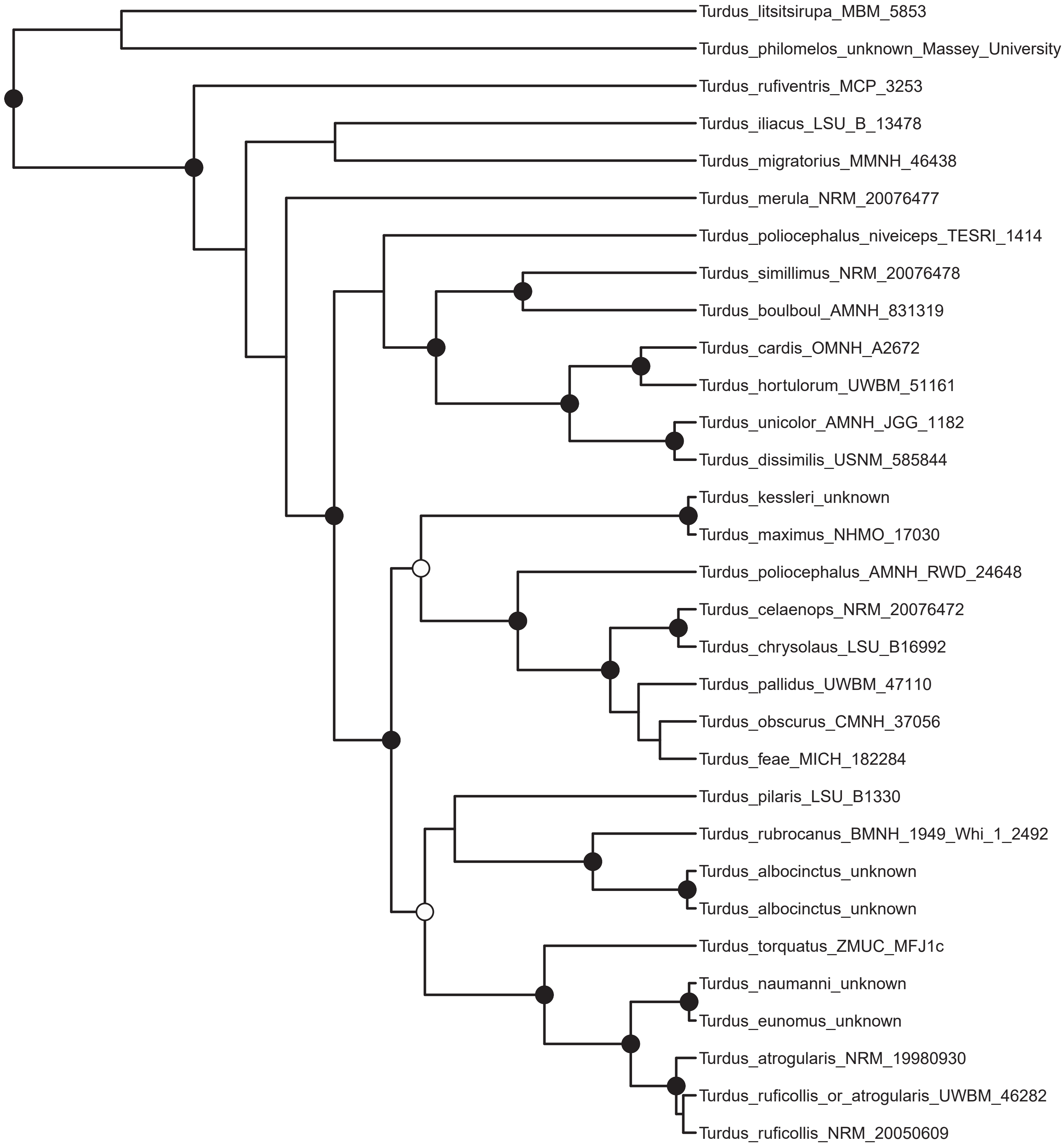
Turdus_litsitsirupa_MBM_5853
Turdus_philomelos_unknown_Massey_University
Turdus_rufiventris_MCP_3253
Turdus_iliacus_LSU_B_13478
Turdus_migratorius_MMNH_46438
Turdus_merula_NRM_20076477
Turdus_poliocephalus_niveiceps_TESRI_1414
Turdus_simillimus_NRM_20076478
Turdus_boulboul_AMNH_831319
Turdus_cardis_OMNH_A2672
Turdus_hortulorum_UWBM_51161
Turdus_unicolor_AMNH_JGG_1182
Turdus_dissimilis_USNM_585844
Turdus_kessleri_unknown
Turdus_maximus_NHMO_17030
Turdus_poliocephalus_AMNH_RWD_24648
Turdus_celaenops_NRM_20076472
Turdus_chrysolaus_LSU_B16992
Turdus_pallidus_UWBM_47110
Turdus_obscurus_CMNH_37056
Turdus_feae_MICH_182284
Turdus_pilaris_LSU_B1330
Turdus_rubrocanus_BMNH_1949_Whi_1_2492
Turdus_albocinctus_unknown
Turdus_albocinctus_unknown
Turdus_torquatus_ZMUC_MFJ1c
Turdus_naumanni_unknown
Turdus_eunomus_unknown
Turdus_atrogularis_NRM_19980930
Turdus_ruficollis_or_atrogularis_UWBM_46282
Turdus_ruficollis_NRM_20050609

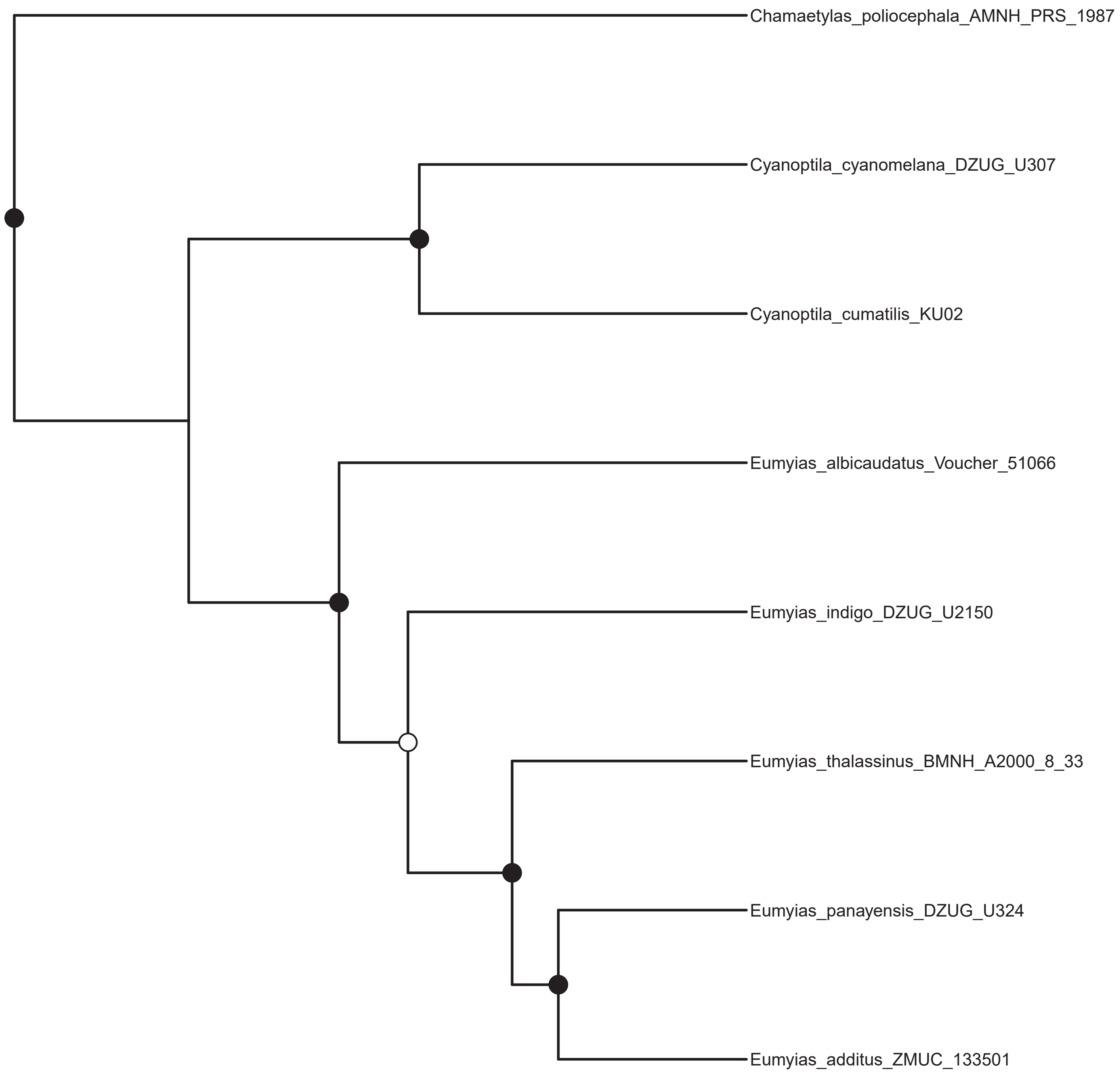
Chamaetylas_poliocephala_AMNH_PRS_1987
Cyanoptila_cyanomelana_DZUG_U307
Cyanoptila_cumatilis_KU02
Eumyias_albicaudatus_Voucher_51066
Eumyias_indigo_DZUG_U2150
Eumyias_thalassinus_BMNH_A2000_8_33
Eumyias_panayensis_DZUG_U324
Eumyias_additus_ZMUC_133501

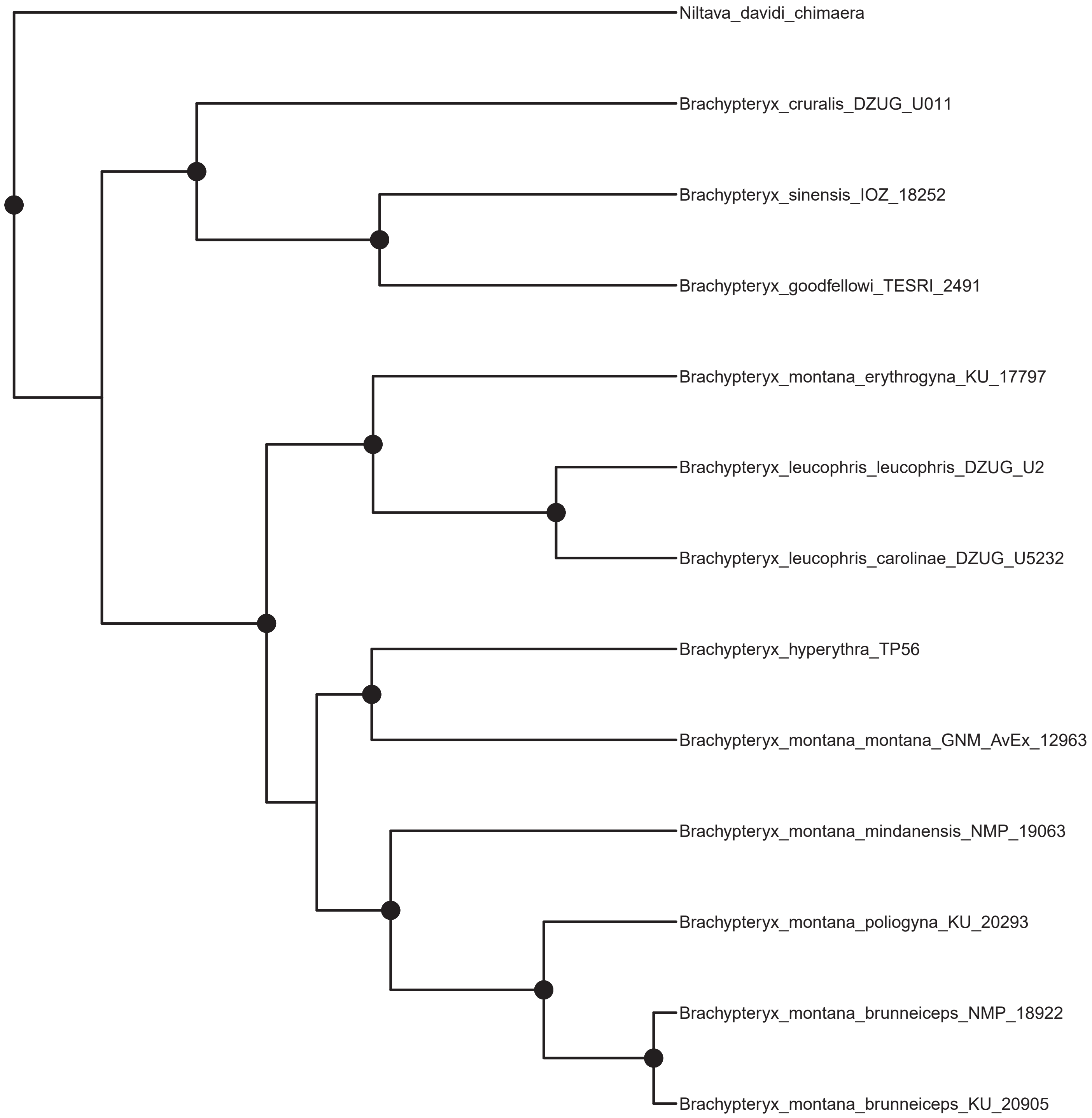
Niltava_davidi_chimaera
Brachypteryx_cruralis_DZUG_U011
Brachypteryx_sinensis_IOZ_18252
Brachypteryx_goodfellowi_TESRI_2491
Brachypteryx_montana_erythrogyna_KU_17797
Brachypteryx_leucophris_leucophris_DZUG_U2
Brachypteryx_leucophris_carolinae_DZUG_U5232
Brachypteryx_hyperythra_TP56
Brachypteryx_montana_montana_GNM_AvEx_12963
Brachypteryx_montana_mindanensis_NMP_19063
Brachypteryx_montana_poliogyna_KU_20293
Brachypteryx_montana_brunneiceps_NMP_18922
Brachypteryx_montana_brunneiceps_KU_20905

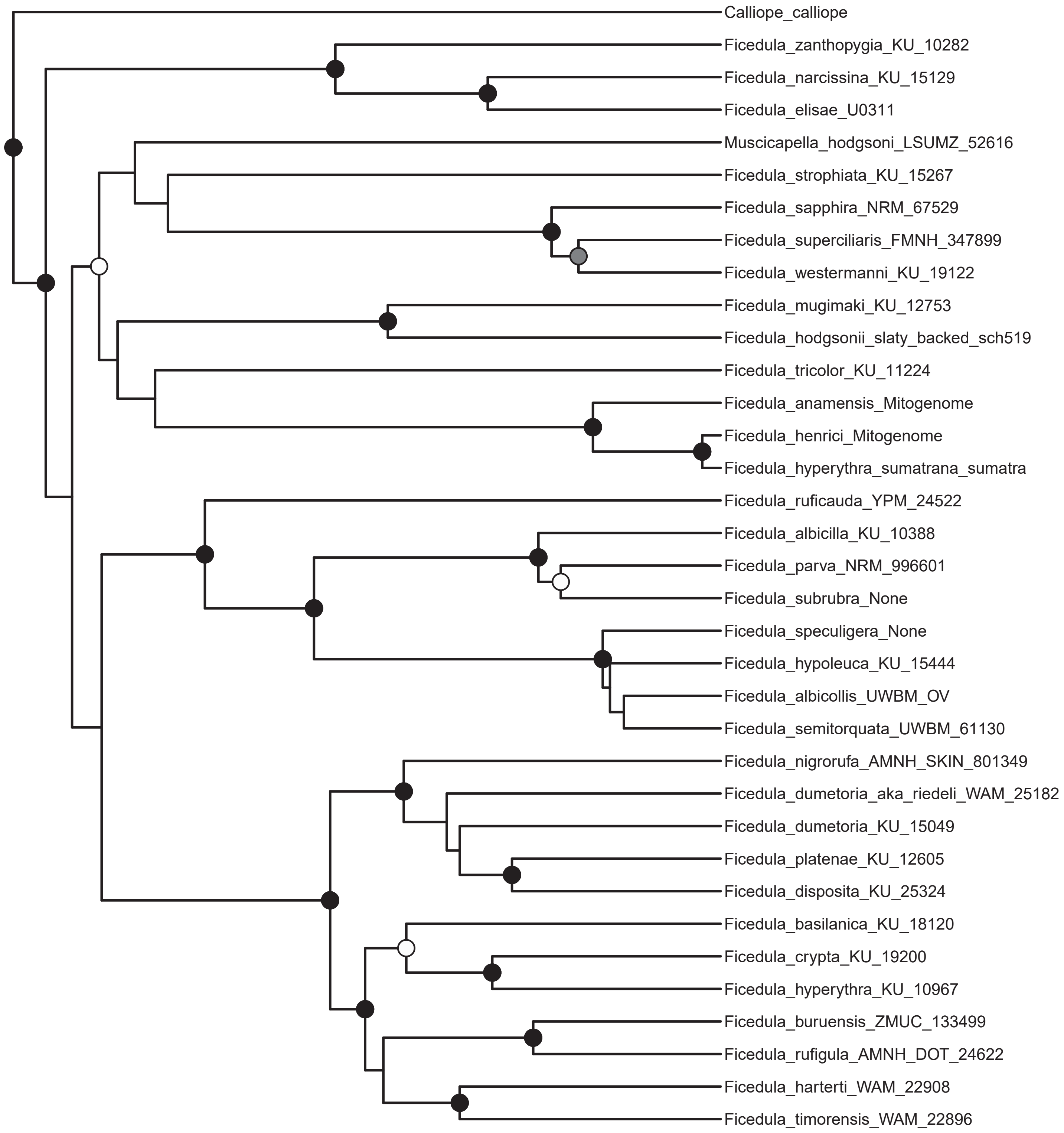
Calliope_calliope
Ficedula_zanthopygia_KU_10282
Ficedula_narcissina_KU_15129
Ficedula_elisae_U0311
Muscicapella_hodgsoni_LSUMZ_52616
Ficedula_strophiata_KU_15267
Ficedula_sapphira_NRM_67529
Ficedula_superciliaris_FMNH_347899
Ficedula_westermanni_KU_19122
Ficedula_mugimaki_KU_12753
Ficedula_hodgsonii_slaty_backed_sch519
Ficedula_tricolor_KU_11224
Ficedula_anamensis_Mitogenome
Ficedula_henrici_Mitogenome
Ficedula_hyperythra_sumatrana_sumatra
Ficedula_ruficauda_YPM_24522
Ficedula_albicilla_KU_10388
Ficedula_parva_NRM_996601
Ficedula_subrubra_None
Ficedula_speculigera_None
Ficedula_hypoleuca_KU_15444
Ficedula_albicollis_UWBM_OV
Ficedula_semitorquata_UWBM_61130
Ficedula_nigrorufa_AMNH_SKIN_801349
Ficedula_dumetoria_aka_riedeli_WAM_25182
Ficedula_dumetoria_KU_15049
Ficedula_platenae_KU_12605
Ficedula_disposita_KU_25324
Ficedula_basilanica_KU_18120
Ficedula_crypta_KU_19200
Ficedula_hyperythra_KU_10967
Ficedula_buruensis_ZMUC_133499
Ficedula_rufigula_AMNH_DOT_24622
Ficedula_harterti_WAM_22908
Ficedula_timorensis_WAM_22896

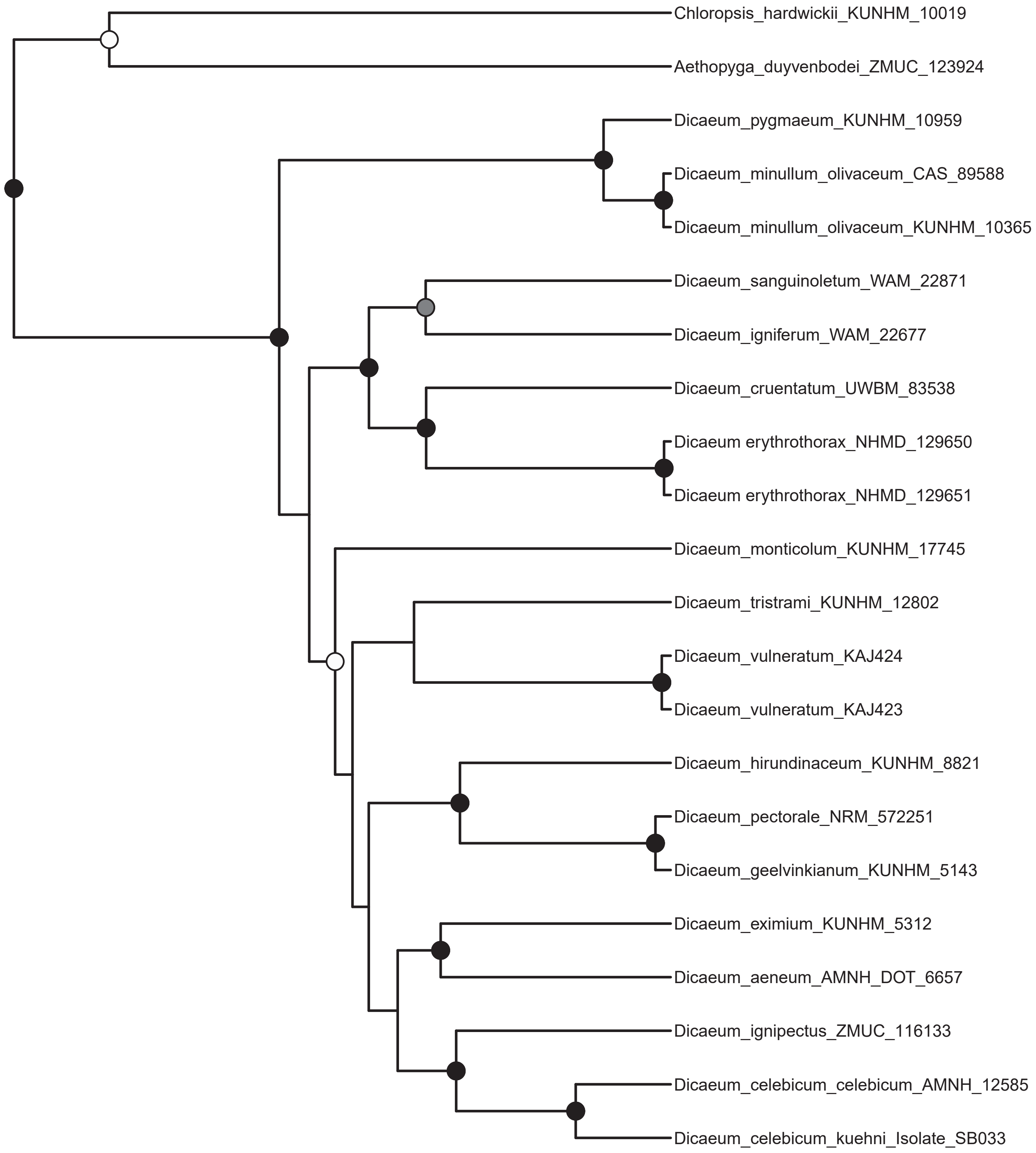
Chloropsis_hardwickii_KUNHM_10019
Aethopyga_duyvenbodei_ZMUC_123924
Dicaeum_pygmaeum_KUNHM_10959
Dicaeum_minullum_olivaceum_CAS_89588
Dicaeum_minullum_olivaceum_KUNHM_10365
Dicaeum_sanguinoletum_WAM_22871
Dicaeum_igniferum_WAM_22677
Dicaeum_cruentatum_UWBM_83538
Dicaeum erythrothorax_NHMD_129650
Dicaeum erythrothorax_NHMD_129651
Dicaeum_monticolum_KUNHM_17745
Dicaeum_tristrami_KUNHM_12802
Dicaeum_vulneratum_KAJ424
Dicaeum_vulneratum_KAJ423
Dicaeum_hirundinaceum_KUNHM_8821
Dicaeum_pectorale_NRM_572251
Dicaeum_geelvinkianum_KUNHM_5143
Dicaeum_eximium_KUNHM_5312
Dicaeum_aeneum_AMNH_DOT_6657
Dicaeum_ignipectus_ZMUC_116133
Dicaeum_celebicum_celebicum_AMNH_12585
Dicaeum_celebicum_kuehni_Isolate_SB033

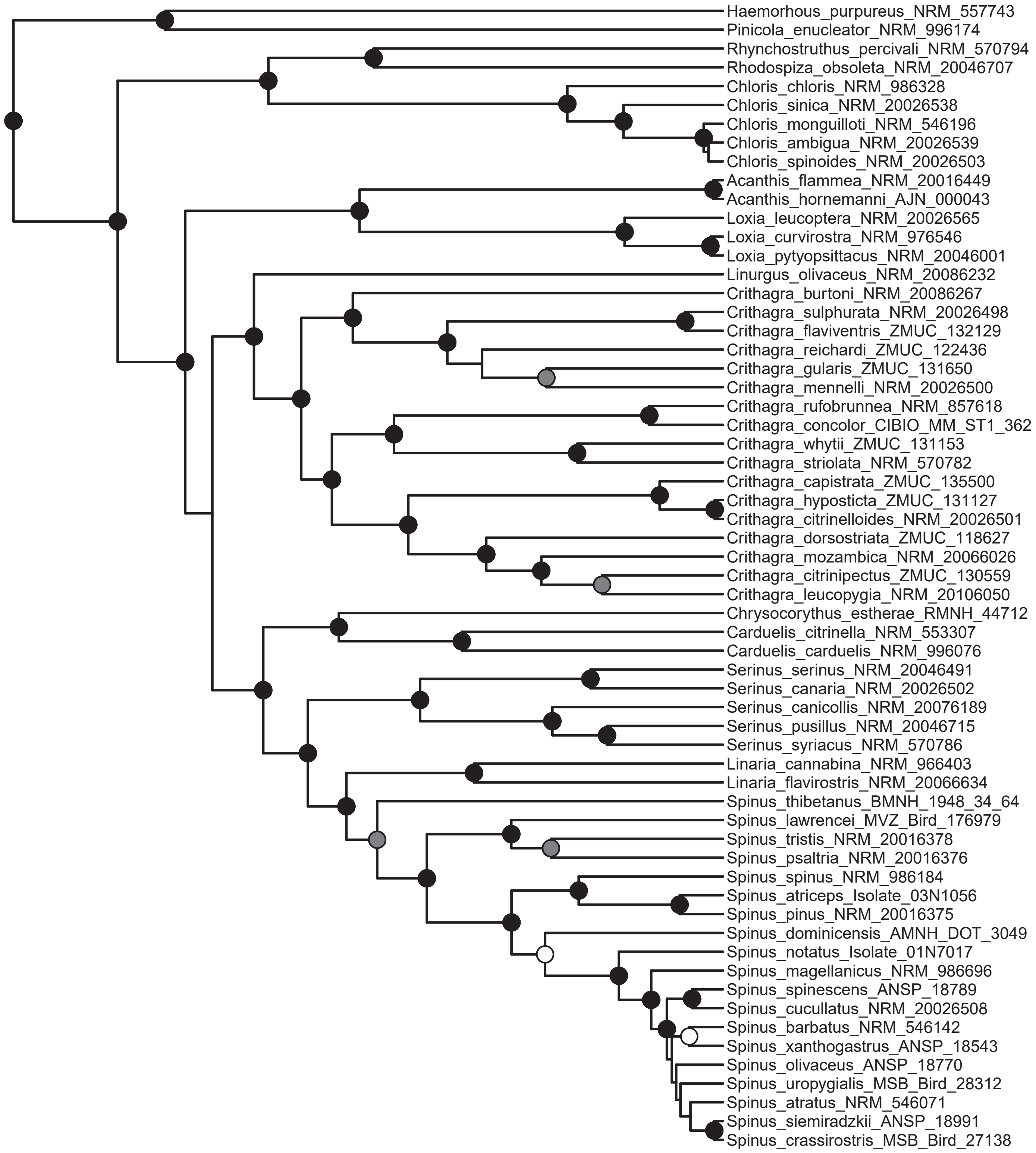
Haemorhous_purpureus_NRM_557743
Pinicola_enucleator_NRM_996174
Rhynchostruthus_percivali_NRM_570794
Rhodospiza_obsoleta_NRM_20046707
Chloris_chloris_NRM_986328
Chloris_sinica_NRM_20026538
Chloris_monguilloti_NRM_546196
Chloris_ambigua_NRM_20026539
Chloris_spinoides_NRM_20026503
Acanthis_flammea_NRM_20016449
Acanthis_hornemanni_AJN_000043
Loxia_leucoptera_NRM_20026565
Loxia_curvirostra_NRM_976546
Loxia_pytyopsittacus_NRM_20046001
Linurgus_olivaceus_NRM_20086232
Crithagra_burtoni_NRM_20086267
Crithagra_sulphurata_NRM_20026498
Crithagra_flaviventris_ZMUC_132129
Crithagra_reichardi_ZMUC_122436
Crithagra_gularis_ZMUC_131650
Crithagra_mennelli_NRM_20026500
Crithagra_rufobrunnea_NRM_857618
Crithagra_concolor_CIBIO_MM_ST1_362
Crithagra_whytii_ZMUC_131153
Crithagra_striolata_NRM_570782
Crithagra_capistrata_ZMUC_135500
Crithagra_hyposticta_ZMUC_131127
Crithagra_citrinelloides_NRM_20026501
Crithagra_dorsostriata_ZMUC_118627
Crithagra_mozambica_NRM_20066026
Crithagra_citrinipectus_ZMUC_130559
Crithagra_leucopygia_NRM_20106050
Chrysocorythus_estherae_RMNH_44712
Carduelis_citrinella_NRM_553307
Carduelis_carduelis_NRM_996076
Serinus_serinus_NRM_20046491
Serinus_canaria_NRM_20026502
Serinus_canicollis_NRM_20076189
Serinus_pusillus_NRM_20046715
Serinus_syriacus_NRM_570786
Linaria_cannabina_NRM_966403
Linaria_flavirostris_NRM_20066634
Spinus_thibetanus_BMNH_1948_34_64
Spinus_lawrencei_MVZ_Bird_176979
Spinus_tristis_NRM_20016378
Spinus_psaltria_NRM_20016376
Spinus_spinus_NRM_986184
Spinus_atriceps_Isolate_03N1056
Spinus_pinus_NRM_20016375
Spinus_dominicensis_AMNH_DOT_3049
Spinus_notatus_Isolate_01N7017
Spinus_magellanicus_NRM_986696
Spinus_spinescens_ANSP_18789
Spinus_cucullatus_NRM_20026508
Spinus_barbatus_NRM_546142
Spinus_xanthogastrus_ANSP_18543
Spinus_olivaceus_ANSP_18770
Spinus_uropygialis_MSB_Bird_28312
Spinus_atratus_NRM_546071
Spinus_siemiradzkii_ANSP_18991
Spinus_crassirostris_MSB_Bird_27138
