## Supplementary File 4 for "The formation of the Indo-Pacific montane avifauna"

**Supplementary File 4.** Ancestral state reconstructions of geographic range (pp. 3–25), elevational range (pp. 26–48), and migratory behavior (pp. 49–71).

**Contents:**
Map of biogeographic regions as defined for this study - p. 2
Meliphagidae - pp. 3, 26, 49
Campephagidae - pp. 4, 27, 50
Pachycephalidae - pp. 5, 28, 51
Rhipiduridae A (*Rhipidura*) - pp. 6, 29, 52
Rhipiduridae B (Lamproliidae) - pp. 7, 30, 53
Corvidae - pp. 8, 31, 54
Petroicidae A (*Microeca*) - pp. 9, 32, 55
Petroicidae B (*Petroica*) - pp. 10, 33, 56
Stenostiridae - pp. 11, 34, 57
Pnoepygidae - pp. 12, 35, 58
Cettiidae - pp. 13, 36, 59
Phylloscopidae - pp. 14, 37, 60
Locustellidae - pp. 15, 38, 61
Sturnidae - pp. 16, 39, 62
Turdidae A (*Geokichla*) - pp. 17, 40, 63
Turdidae B (*Zoothera*) - pp. 18, 41, 64
Turdidae C (*Turdus*) - pp. 19, 42, 65
Muscicapidae A (*Eumyias*) - pp. 20, 43, 66
Muscicapidae B (*Brachypteryx*) - pp. 21, 44, 67
Muscicapidae C (*Ficedula*) - pp. 22, 45, 68
Dicaeidae - pp. 23, 46, 69
Motacillidae - pp. 24, 47, 70
Fringillidae - pp. 25, 48, 71

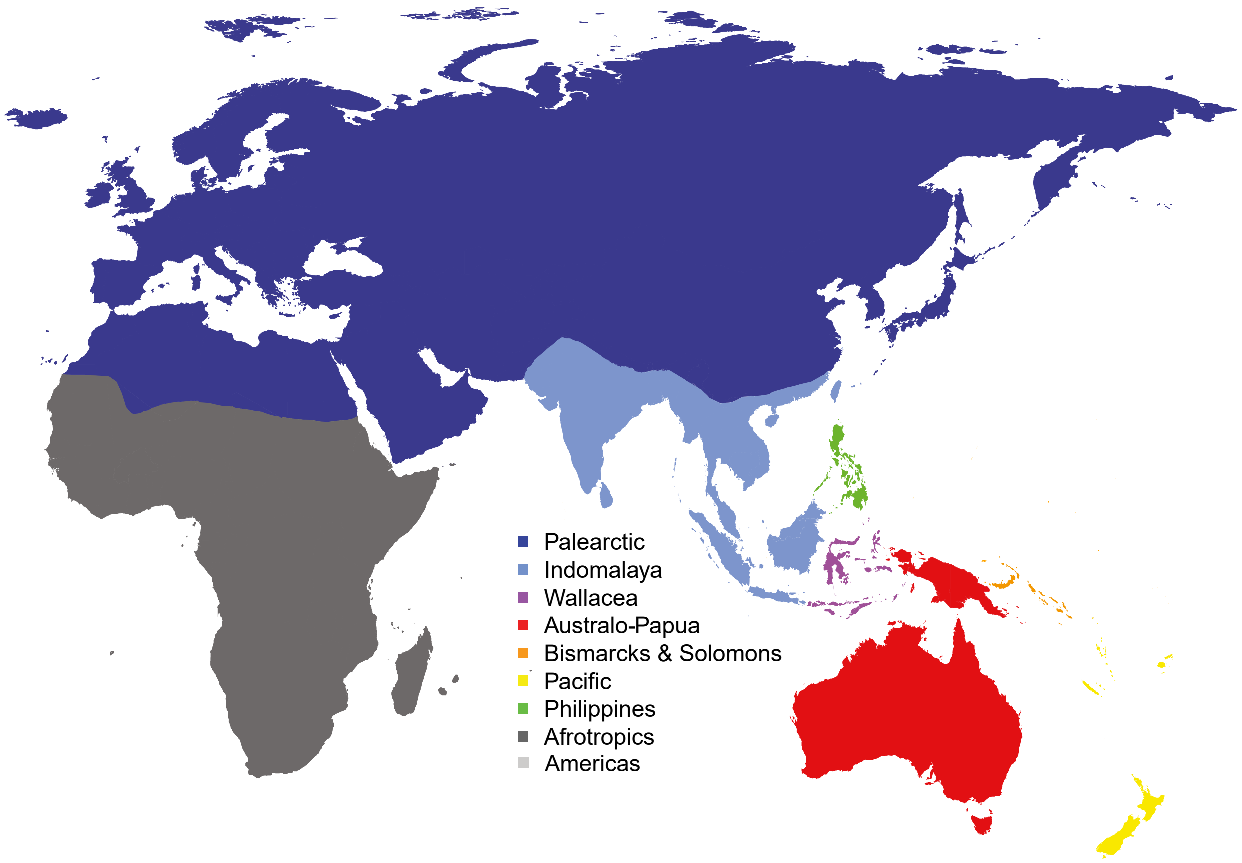


Biogeographic regions as defined for this study. The Americas are not shown.

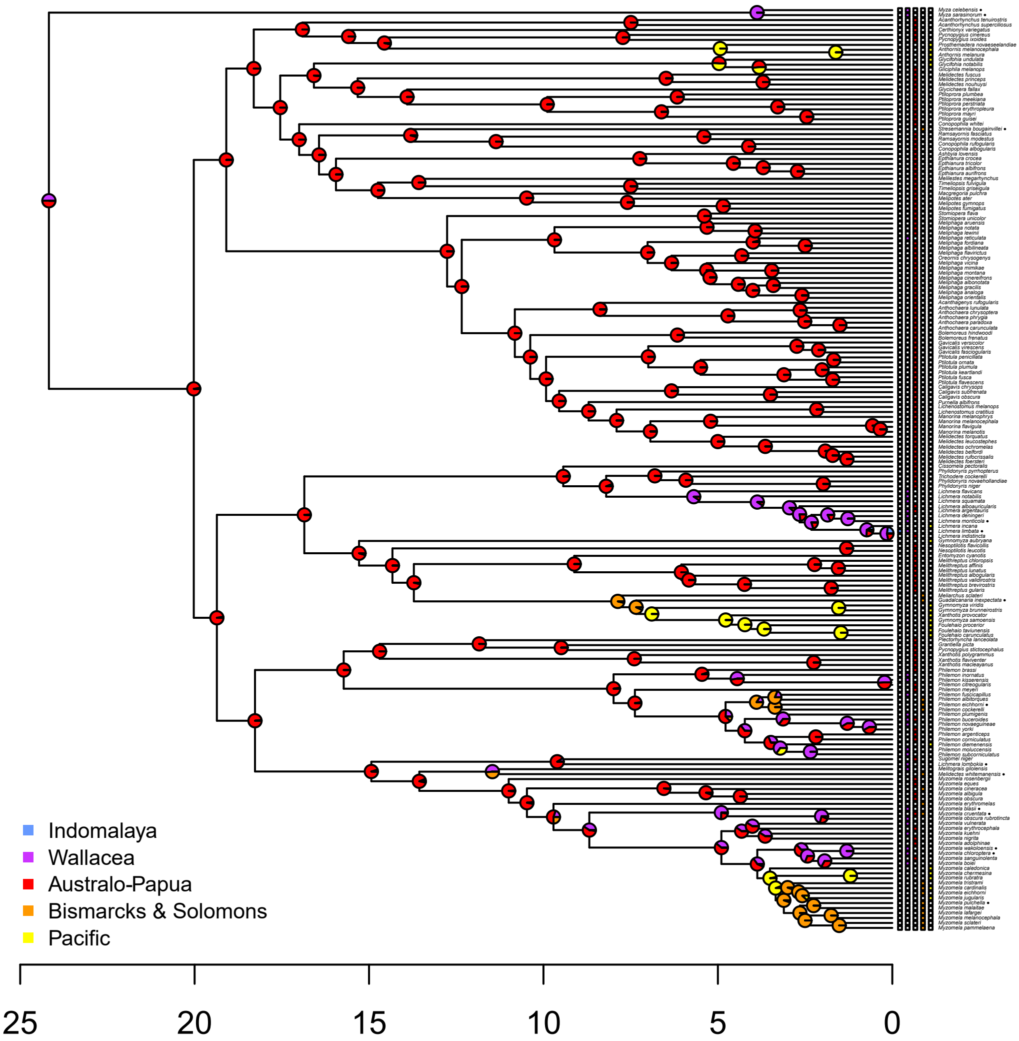
Indomalaya
Wallacea
Australo-Papua
Bismarcks & Solomons
Pacific
25
20
15
10
5
0

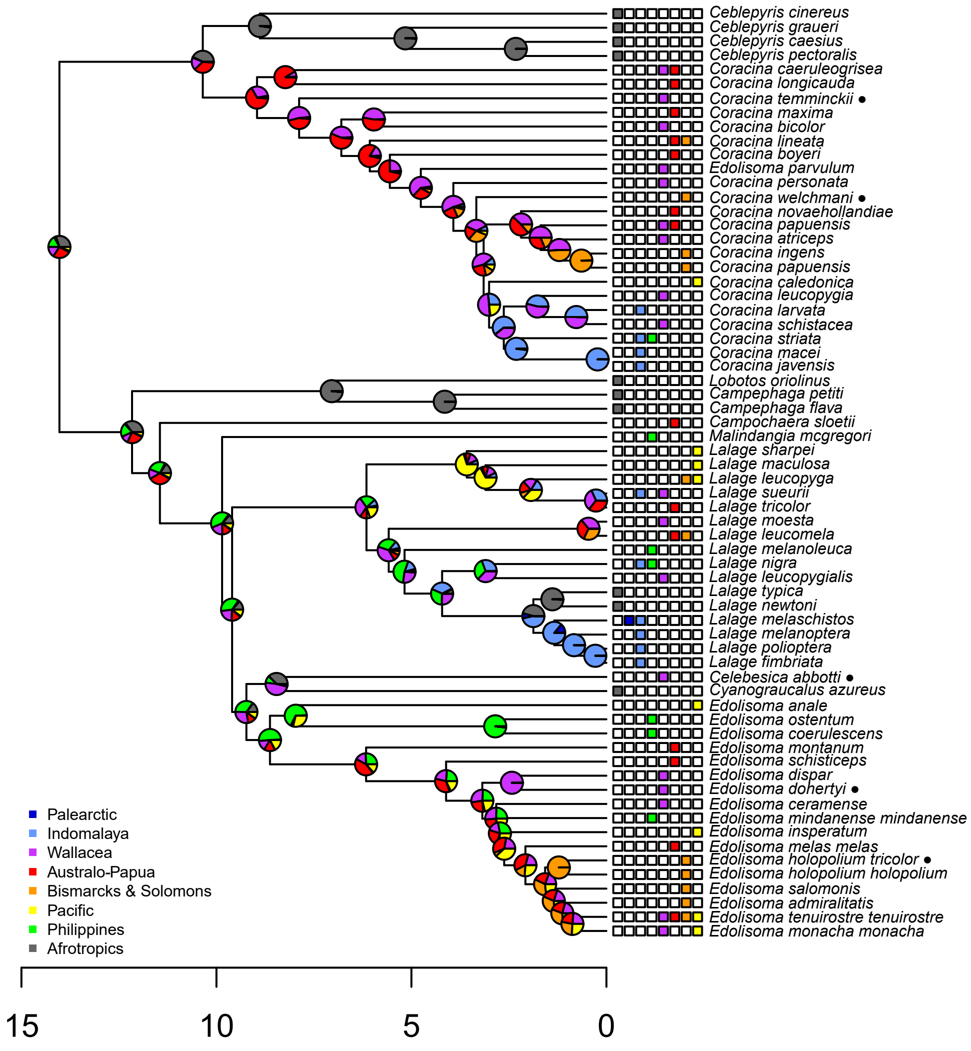

Ceblepyris cinereus
Ceblepyris graueri
Ceblepyris caesius
Ceblepyris pectoralis
Coracina caeruleogrisea
Coracina longicauda
Coracina temminckii ●
Coracina maxima
Coracina bicolor
Coracina lineata
Coracina boyeri
Edolisoma parvulum
Coracina personata
Coracina welchmani ●
Coracina novaehollandiae
Coracina papuensis
Coracina atriceps
Coracina ingens
Coracina papuensis
Coracina caledonica
Coracina leucopygia
Coracina larvata
Coracina schistacea
Coracina striata
Coracina macei
Coracina javensis
Lobotos oriolinus
Campephaga petiti
Campephaga flava
Campochaera sloetii
Malindangia mcgregori
Lalage sharpei
Lalage maculosa
Lalage leucopyga
Lalage sueurii
Lalage tricolor
Lalage moesta
Lalage leucomela
Lalage melanoleuca
Lalage nigra
Lalage leucopygialis
Lalage typica
Lalage newtoni
Lalage melaschistos
Lalage melanoptera
Lalage polioptera
Lalage fimbriata
Celebesica abbotti ●
Cyanograucalus azureus
Edolisoma anale
Edolisoma ostentum
Edolisoma coerulescens
Edolisoma montanum
Edolisoma schisticeps
Edolisoma dispar
Edolisoma dohertyi ●
Edolisoma ceramense
Edolisoma mindanense mindanense
Edolisoma insperatum
Edolisoma melas melas
Edolisoma holopolium tricolor ●
Edolisoma holopolium holopolium
Edolisoma salomonis
Edolisoma admiralitatis
Edolisoma tenuirostre tenuirostre
Edolisoma monacha monacha
Palearctic
Indomalaya
Wallacea
Australo-Papua
Bismarcks & Solomons
Pacific
Philippines
Afrotropics
15
10
5
0

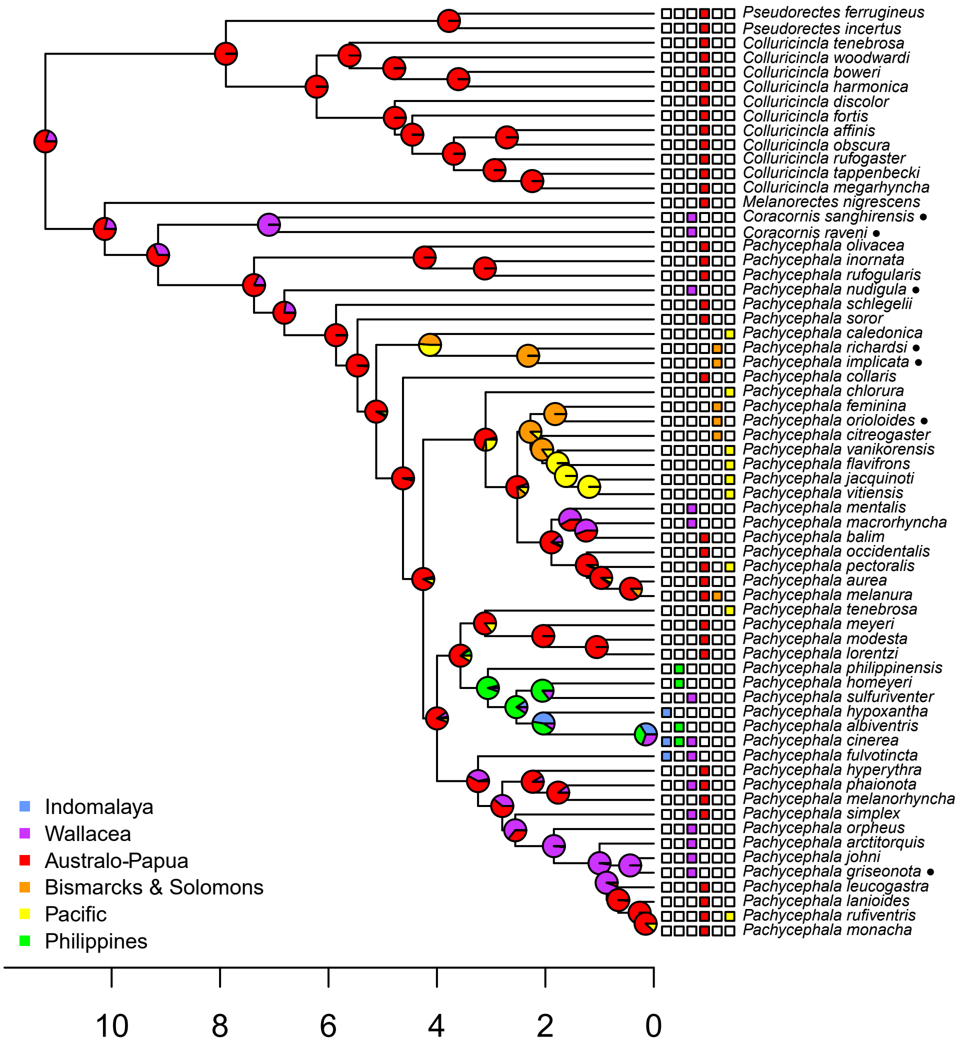
Pseudorectes ferrugineus
Pseudorectes incertus
Colluricincla tenebrosa
Colluricincla woodwardi
Colluricincla boweri
Colluricincla harmonica
Colluricincla discolor
Colluricincla fortis
Colluricincla affinis
Colluricincla obscura
Colluricincla rufogaster
Colluricincla tappenbecki
Colluricincla megarhyncha
Melanorectes nigrescens
Coracornis sanghirensis ●
Coracornis raveni ●
Pachycephala olivacea
Pachycephala inornata
Pachycephala rufogularis
Pachycephala nudigula ●
Pachycephala schlegelii
Pachycephala soror
Pachycephala caledonica
Pachycephala richardsi ●
Pachycephala implicata ●
Pachycephala collaris
Pachycephala chlorura
Pachycephala feminina
Pachycephala orioloides ●
Pachycephala citreogaster
Pachycephala vanikorensis
Pachycephala flavifrons
Pachycephala jacquinoti
Pachycephala vitiensis
Pachycephala mentalis
Pachycephala macrorhyncha
Pachycephala balim
Pachycephala occidentalis
Pachycephala pectoralis
Pachycephala aurea
Pachycephala melanura
Pachycephala tenebrosa
Pachycephala meyeri
Pachycephala modesta
Pachycephala lorentzi
Pachycephala philippinensis
Pachycephala homeyeri
Pachycephala sulfuriventer
Pachycephala hypoxantha
Pachycephala albiventris
Pachycephala cinerea
Pachycephala fulvotincta
Pachycephala hyperythra
Pachycephala phaionota
Pachycephala melanorhyncha
Pachycephala simplex
Pachycephala orpheus
Pachycephala arctitorquis
Pachycephala johni
Pachycephala griseonota ●
Pachycephala leucogastra
Pachycephala lanioides
Pachycephala rufiventris
Pachycephala monacha
Indomalaya
Wallacea
Australo-Papua
Bismarcks & Solomons
Pacific
Philippines
10
8
6
4
2
0

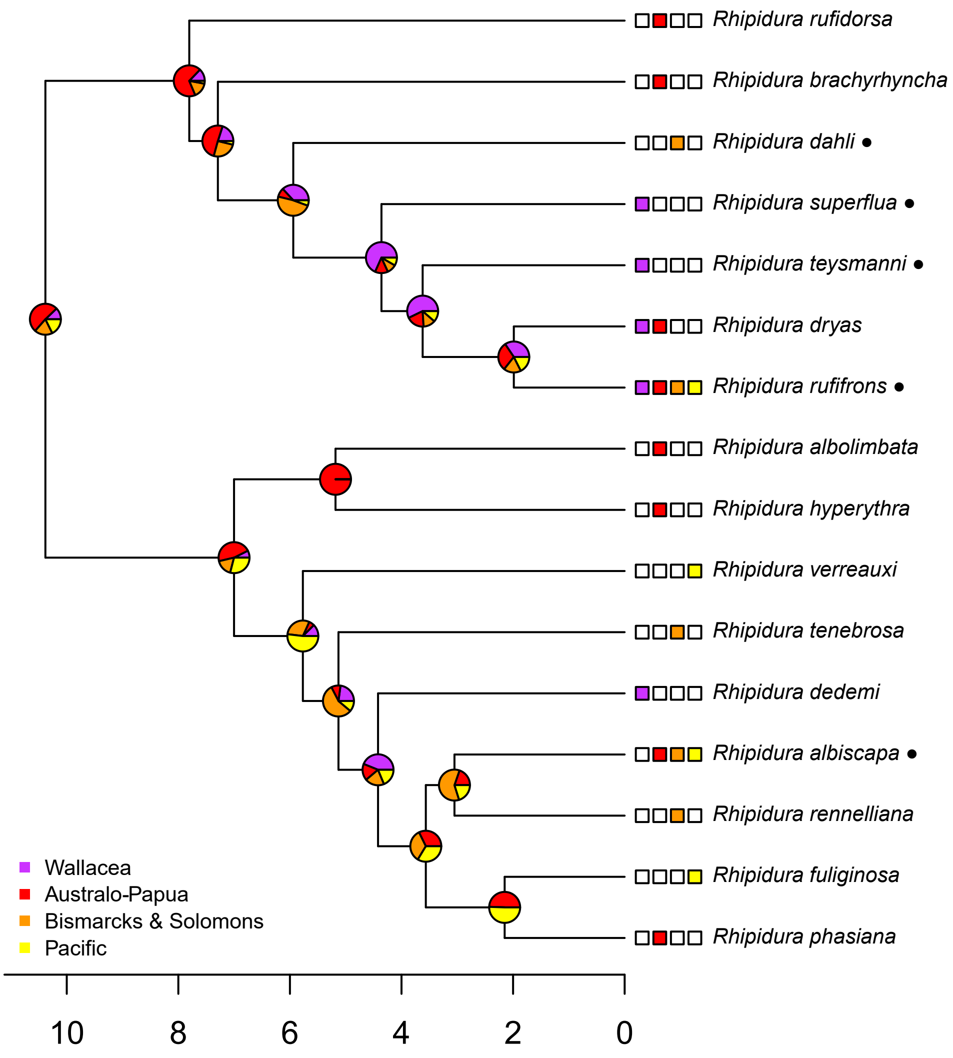

Rhipidura rufidorsa
Rhipidura brachyrhyncha
Rhipidura dahli ●
Rhipidura superflua ●
Rhipidura teysmanni ●
Rhipidura dryas
Rhipidura rufifrons ●
Rhipidura albolimbata
Rhipidura hyperythra
Rhipidura verreauxi
Rhipidura tenebrosa
Rhipidura dedemi
Rhipidura albiscapa ●
Rhipidura rennelliana
Rhipidura fuliginosa
Rhipidura phasiana
Wallacea
Australo-Papua
Bismarcks & Solomons
Pacific
10
8
6
4
2
0

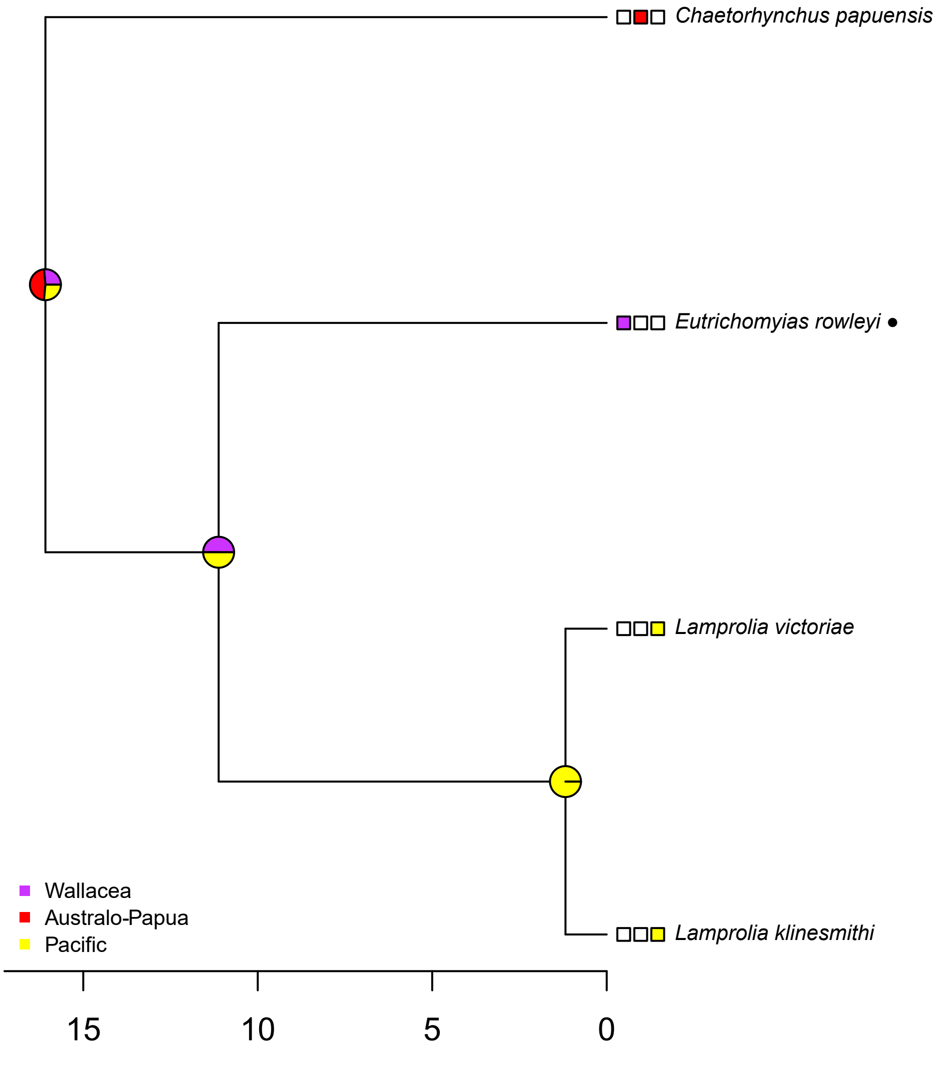

Chaetorhynchus papuensis
Eutrichomyias rowleyi
Lamprolia victoriae
Lamprolia klinesmithi
Wallacea
Australo-Papua
Pacific
15
10
5
0

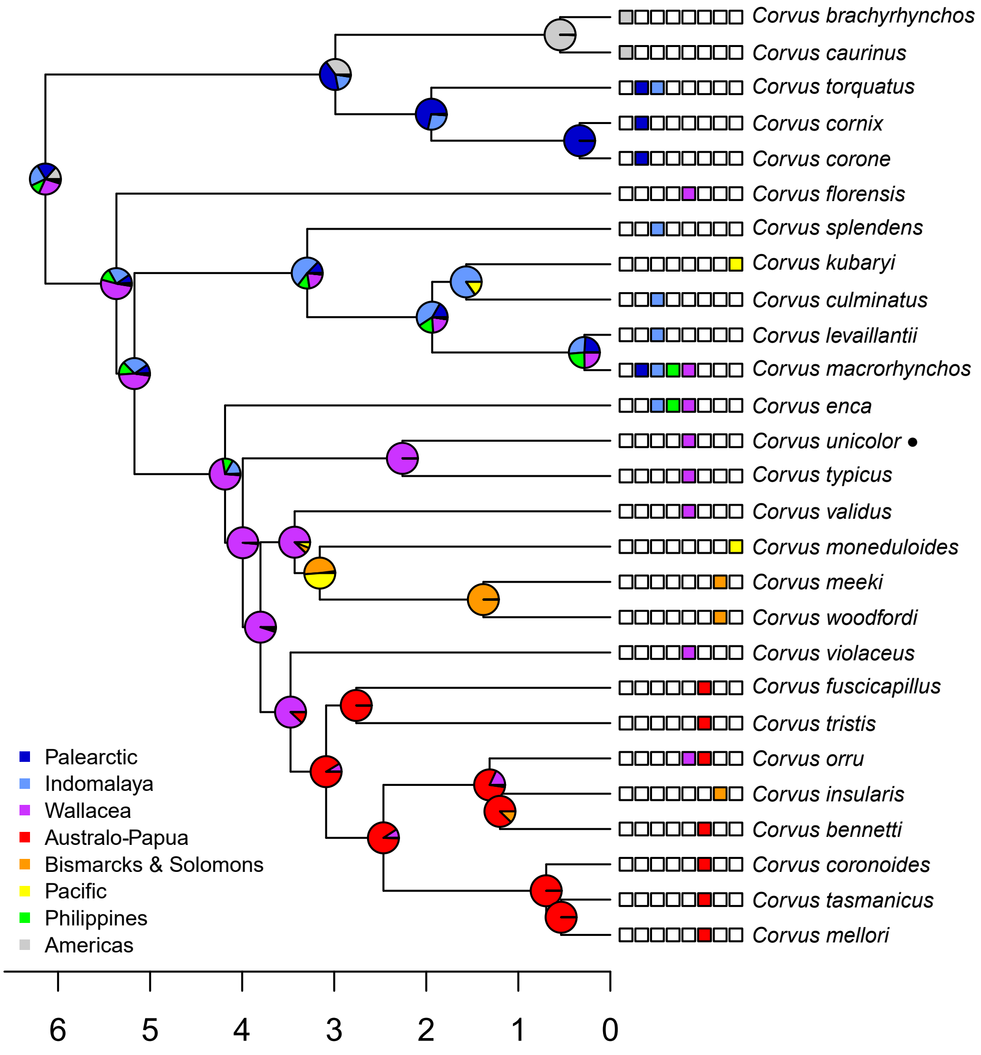

Corvus brachyrhynchos
Corvus caurinus
Corvus torquatus
Corvus cornix
Corvus corone
Corvus florensis
Corvus splendens
Corvus kubaryi
Corvus culminatus
Corvus levaillantii
Corvus macrorhynchos
Corvus enca
Corvus unicolor ●
Corvus typicus
Corvus validus
Corvus moneduloides
Corvus meeki
Corvus woodfordi
Corvus violaceus
Corvus fuscicapillus
Corvus tristis
Corvus orru
Corvus insularis
Corvus bennetti
Corvus coronoides
Corvus tasmanicus
Corvus mellori
Palearctic
Indomalaya
Wallacea
Australo-Papua
Bismarcks & Solomons
Pacific
Philippines
Americas
6
5
4
3
2
1
0

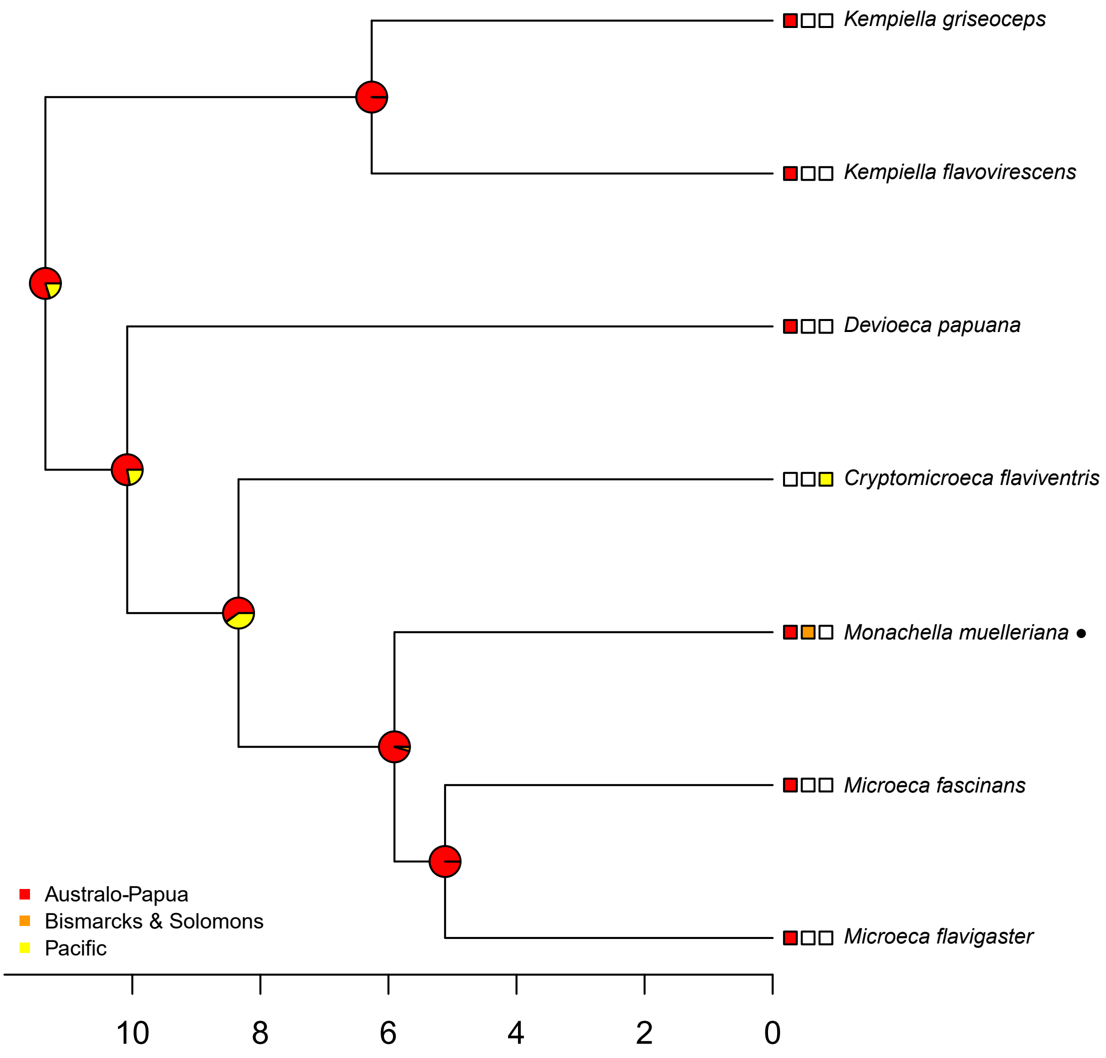

Kempiella griseoceps
Kempiella flavovirescens
Devioeca papuana
Cryptomicroeca flaviventris
Monachella muelleriana ●
Microeca fascinans
Microeca flavigaster
Australo-Papua
Bismarcks & Solomons
Pacific
10
8
6
4
2
0

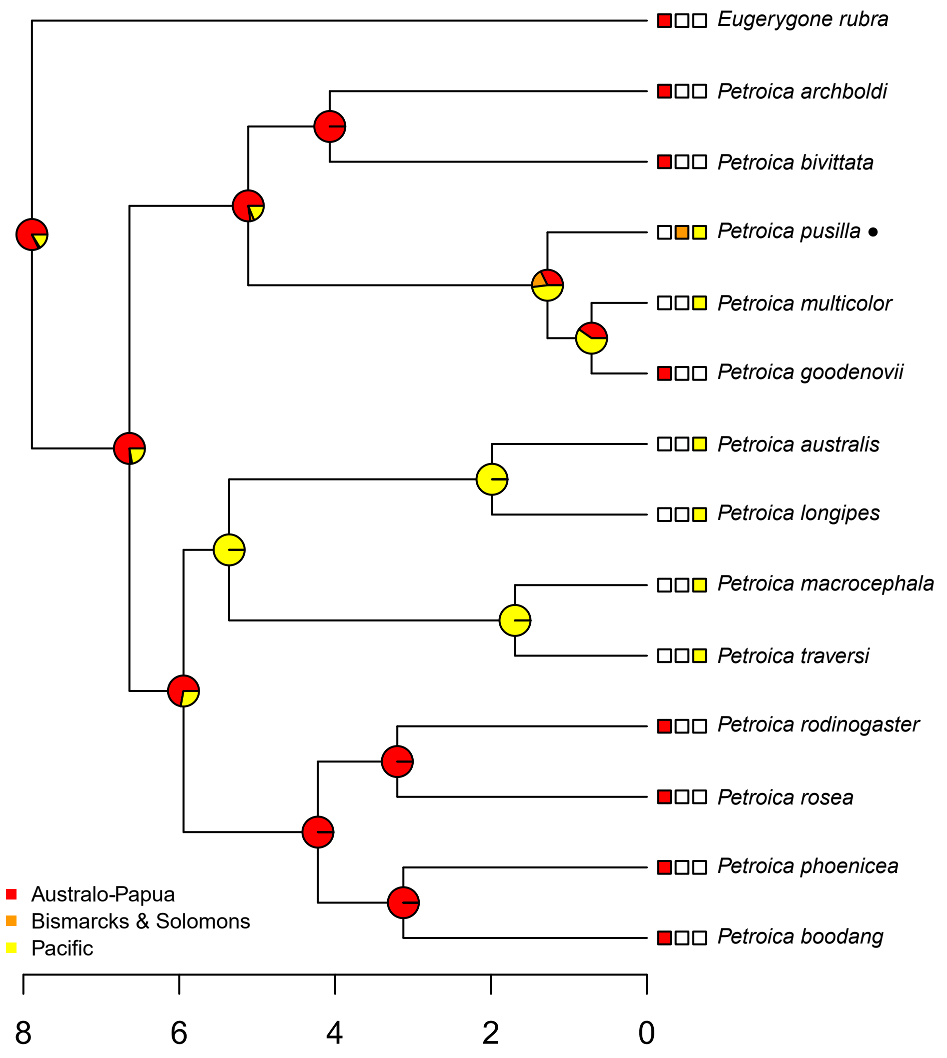

Eugerygone rubra
Petroica archboldi
Petroica bivittata
Petroica pusilla
Petroica multicolor
Petroica goodenovii
Petroica australis
Petroica longipes
Petroica macrocephala
Petroica traversi
Petroica rodinogaster
Petroica rosea
Petroica phoenicea
Petroica boodang
Australo-Papua
Bismarcks & Solomons
Pacific
8
6
4
2
0

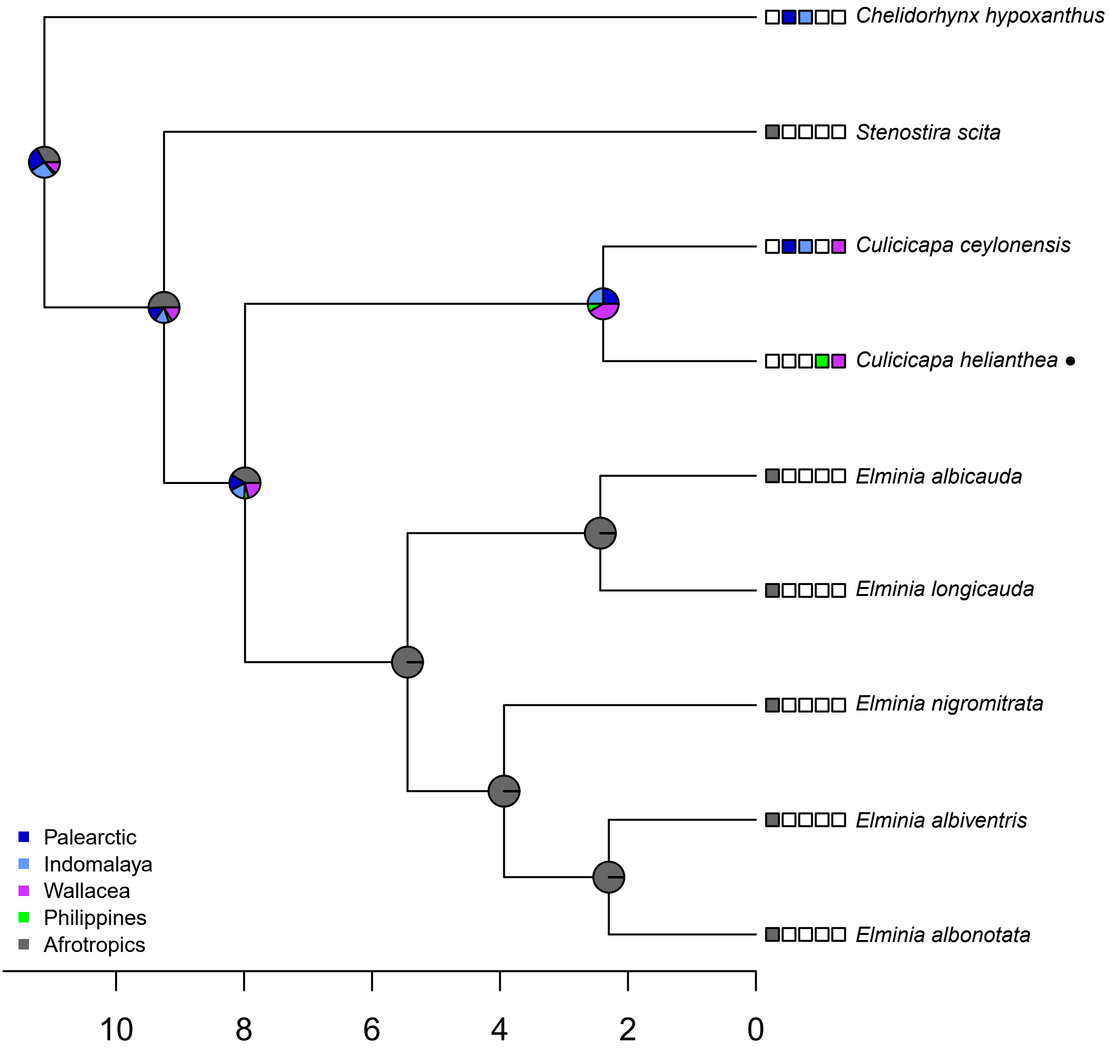

Chelidorhynx hypoxanthus
Stenostira scita
Culicicapa ceylonensis
Culicicapa helianthea ●
Elminia albicauda
Elminia longicauda
Elminia nigromitrata
Elminia albiventris
Elminia albonotata
Palearctic
Indomalaya
Wallacea
Philippines
Afrotropics
10
8
6
4
2
0

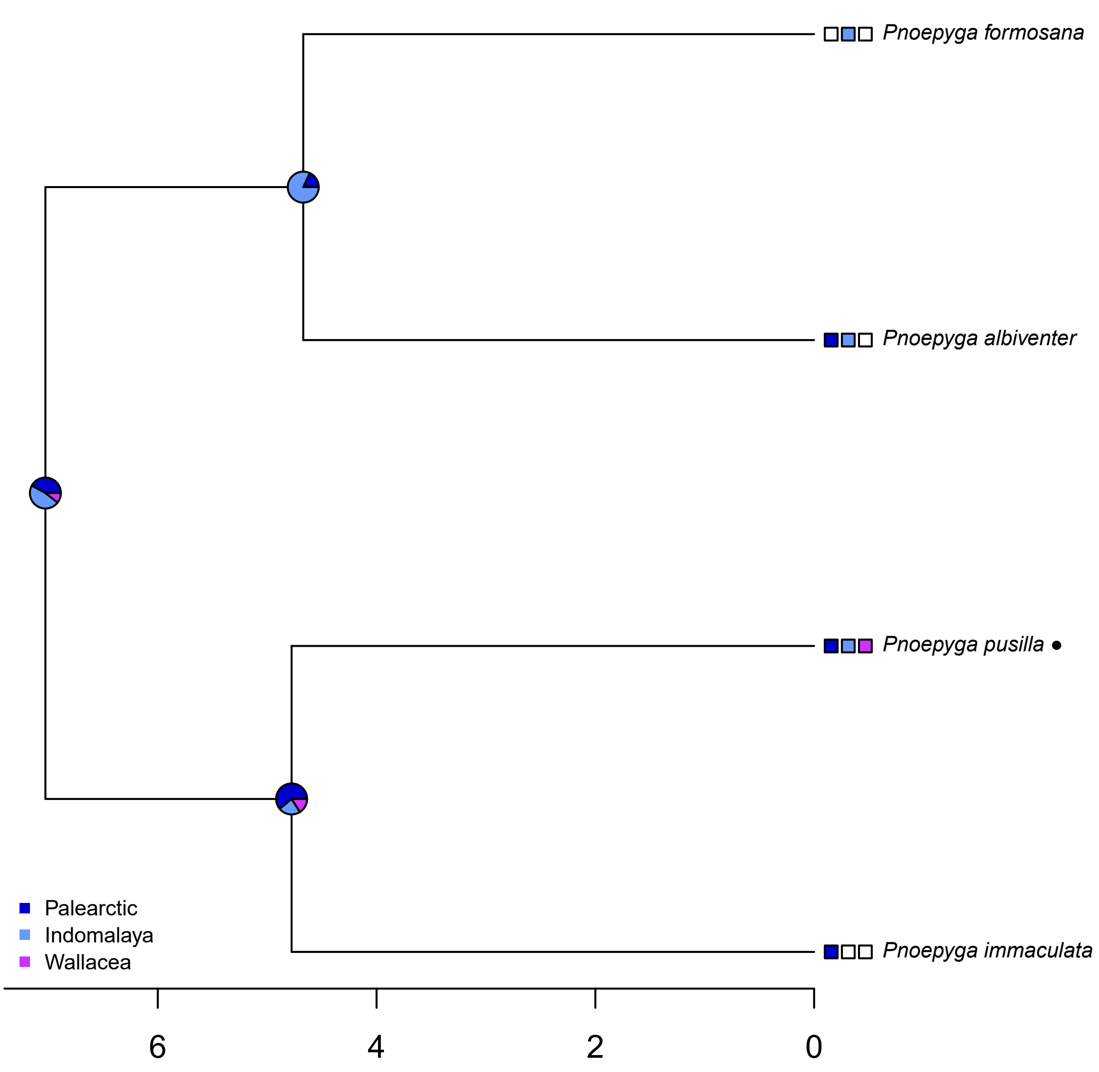

Pnoepyga formosana
Pnoepyga albiventer
Pnoepyga pusilla
Pnoepyga immaculata
Palearctic
Indomalaya
Wallacea
6
4
2
0

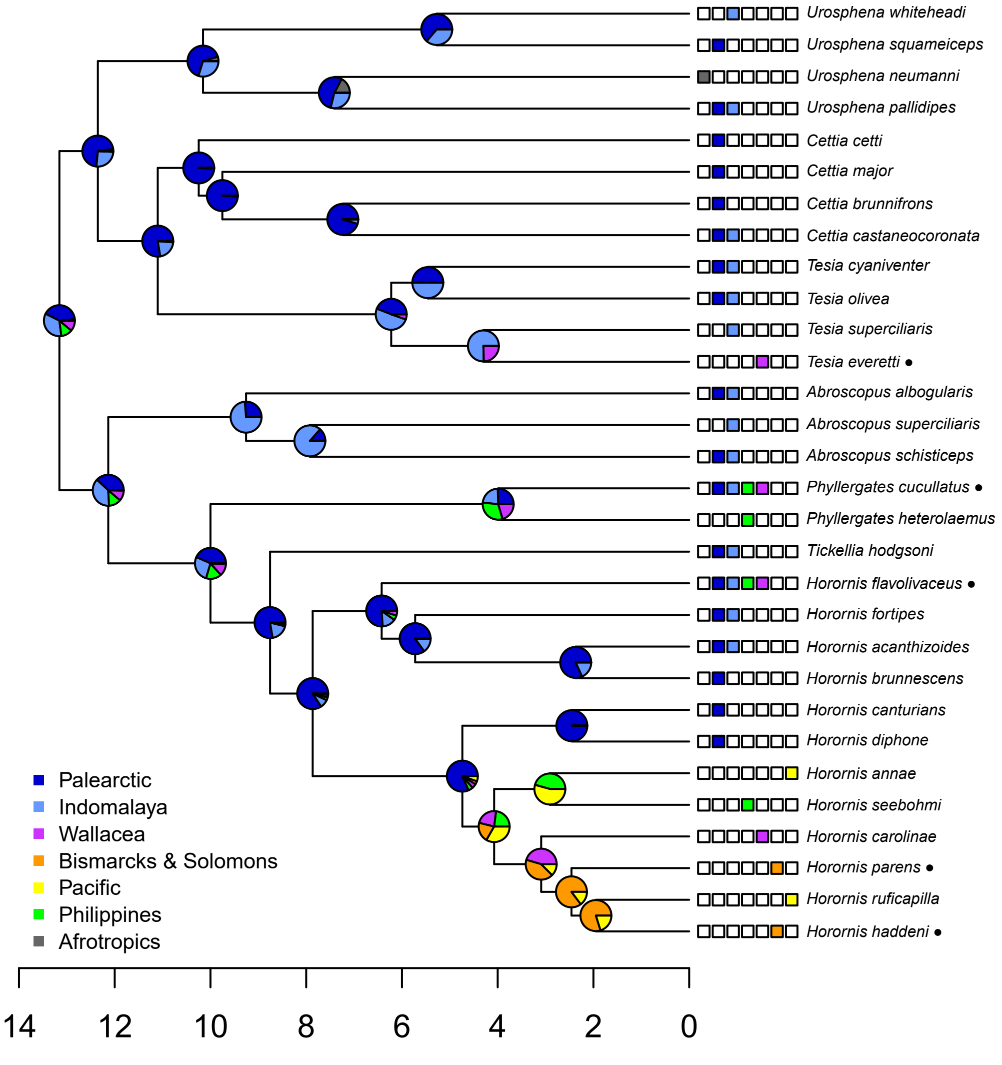

Urosphena whiteheadi
Urosphena squameiceps
Urosphena neumanni
Urosphena pallidipes
Cettia cetti
Cettia major
Cettia brunnifrons
Cettia castaneocoronata
Tesia cyaniventer
Tesia olivea
Tesia superciliaris
Tesia everetti •
Abroscopus albogularis
Abroscopus superciliaris
Abroscopus schisticeps
Phyllergates cucullatus •
Phyllergates heterolaemus
Tickellia hodgsoni
Horornis flavolivaceus •
Horornis fortipes
Horornis acanthizoides
Horornis brunnescens
Horornis canturians
Horornis diphone
Horornis annae
Horornis seebohmi
Horornis carolinae
Horornis parens •
Horornis ruficapilla
Horornis haddeni •
Palearctic
Indomalaya
Wallacea
Bismarcks & Solomons
Pacific
Philippines
Afrotropics
14
12
10
8
6
4
2
0

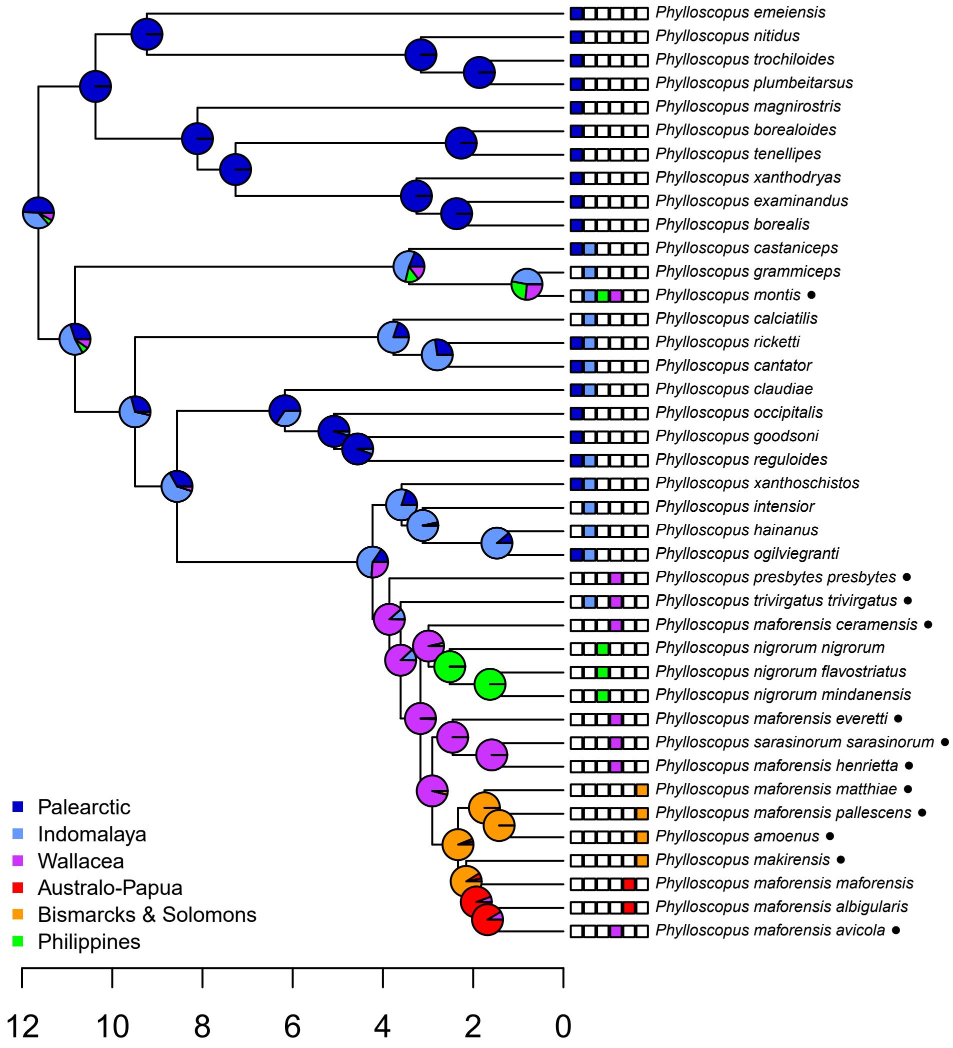

Phylloscopus emeiensis
Phylloscopus nitidus
Phylloscopus trochiloides
Phylloscopus plumbeitarsus
Phylloscopus magnirostris
Phylloscopus borealoides
Phylloscopus tenellipes
Phylloscopus xanthodryas
Phylloscopus examinandus
Phylloscopus borealis
Phylloscopus castaniceps
Phylloscopus grammiceps
Phylloscopus montis ●
Phylloscopus calciatilis
Phylloscopus ricketti
Phylloscopus cantator
Phylloscopus claudiae
Phylloscopus occipitalis
Phylloscopus goodsoni
Phylloscopus reguloides
Phylloscopus xanthoschistos
Phylloscopus intensior
Phylloscopus hainanus
Phylloscopus ogilviegranti
Phylloscopus presbytes presbytes ●
Phylloscopus trivirgatus trivirgatus ●
Phylloscopus maforensis ceramensis ●
Phylloscopus nigrorum nigrorum
Phylloscopus nigrorum flavostriatus
Phylloscopus nigrorum mindanensis
Phylloscopus maforensis everetti ●
Phylloscopus sarasinorum sarasinorum ●
Phylloscopus maforensis henrietta ●
Phylloscopus maforensis matthiae ●
Phylloscopus maforensis pallescens ●
Phylloscopus amoenus ●
Phylloscopus makirensis ●
Phylloscopus maforensis maforensis
Phylloscopus maforensis albigularis
Phylloscopus maforensis avicola ●
Palearctic
Indomalaya
Wallacea
Australo-Papua
Bismarcks & Solomons
Philippines
12
10
8
6
4
2
0

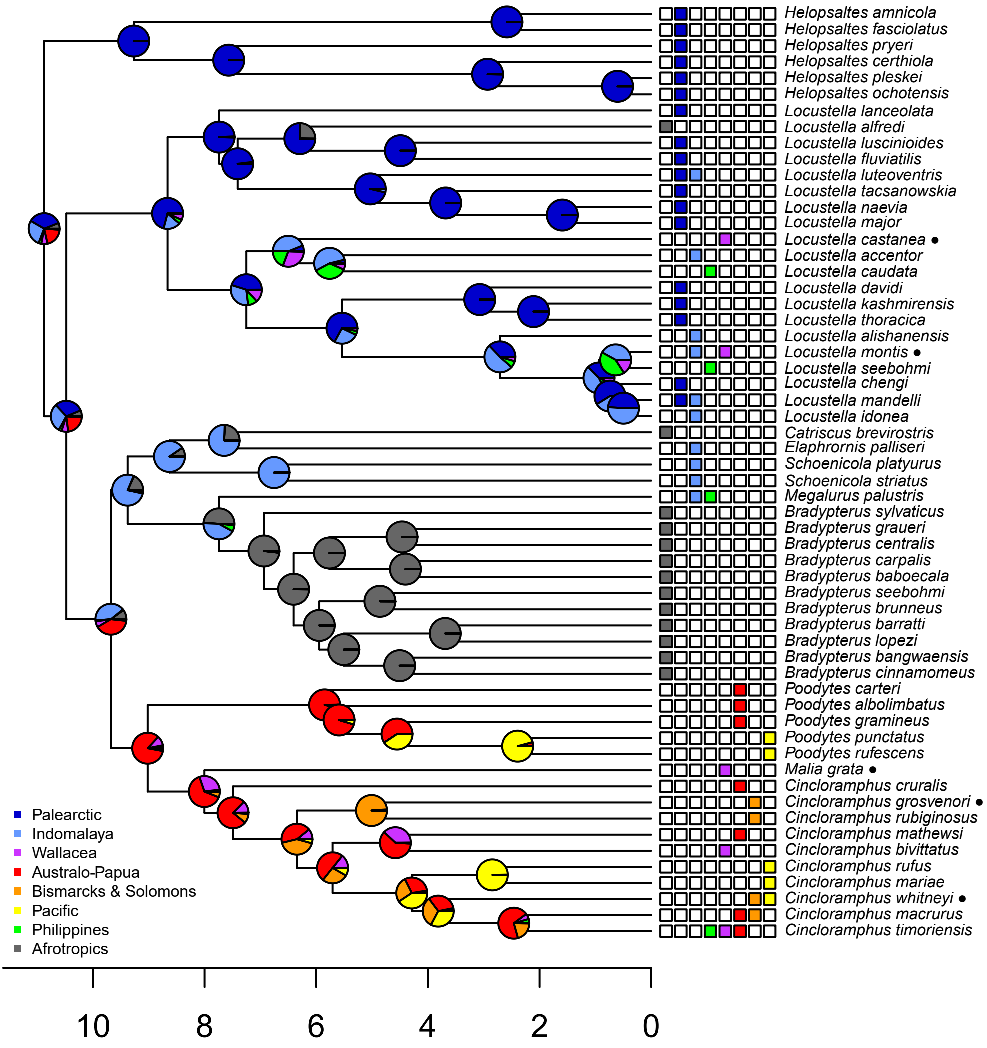

Helopsaltes amnicola
Helopsaltes fasciolatus
Helopsaltes pryeri
Helopsaltes certhiola
Helopsaltes pleskei
Helopsaltes ochotensis
Locustella lanceolata
Locustella alfredi
Locustella luscinioides
Locustella fluviatilis
Locustella luteoventris
Locustella tacsanowskia
Locustella naevia
Locustella major
Locustella castanea ●
Locustella accentor
Locustella caudata
Locustella davidi
Locustella kashmirensis
Locustella thoracica
Locustella alishanensis
Locustella montis ●
Locustella seebohmi
Locustella chengi
Locustella mandelli
Locustella idonea
Catriscus brevirostris
Elaphrornis palliseri
Schoenicola platyurus
Schoenicola striatus
Megalurus palustris
Bradypterus sylvaticus
Bradypterus graueri
Bradypterus centralis
Bradypterus carpalis
Bradypterus baboecala
Bradypterus seebohmi
Bradypterus brunneus
Bradypterus barratti
Bradypterus lopezi
Bradypterus bangwaensis
Bradypterus cinnamomeus
Poodytes carteri
Poodytes albolimbatus
Poodytes gramineus
Poodytes punctatus
Poodytes rufescens
Malia grata ●
Cincloramphus cruralis
Cincloramphus grosvenori ●
Cincloramphus rubiginosus
Cincloramphus mathewsi
Cincloramphus bivittatus
Cincloramphus rufus
Cincloramphus mariae
Cincloramphus whitneyi ●
Cincloramphus macrurus
Cincloramphus timoriensis
Palearctic
Indomalaya
Wallacea
Australo-Papua
Bismarcks & Solomons
Pacific
Philippines
Afrotropics
10
8
6
4
2
0

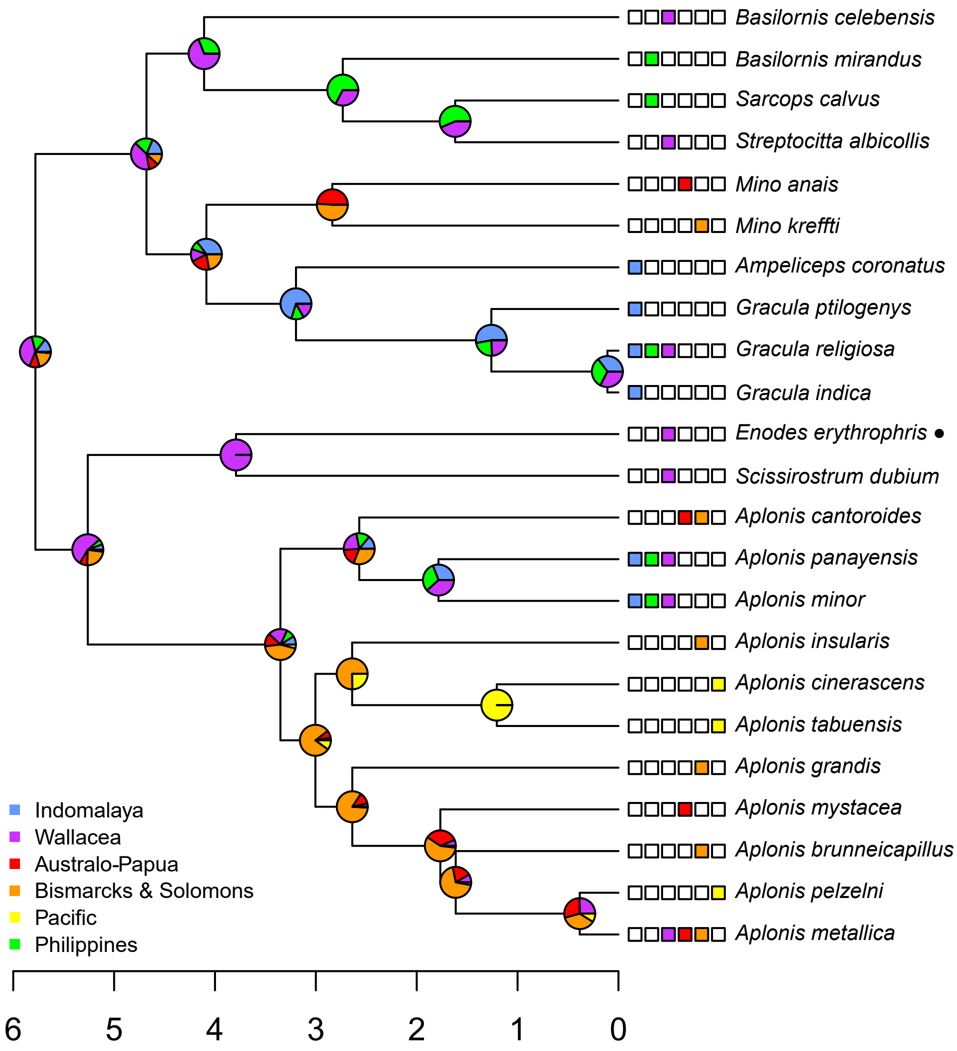

Basilornis celebensis
Basilornis mirandus
Sarcops calvus
Streptocitta albicollis
Mino anais
Mino kreffti
Ampeliceps coronatus
Gracula ptilogenys
Gracula religiosa
Gracula indica
Enodes erythrophris ●
Scissirostrum dubium
Aplonis cantoroides
Aplonis panayensis
Aplonis minor
Aplonis insularis
Aplonis cinerascens
Aplonis tabuensis
Aplonis grandis
Aplonis mystacea
Aplonis brunneicapillus
Aplonis pelzelni
Aplonis metallica
Indomalaya
Wallacea
Australo-Papua
Bismarcks & Solomons
Pacific
Philippines
6
5
4
3
2
1
0

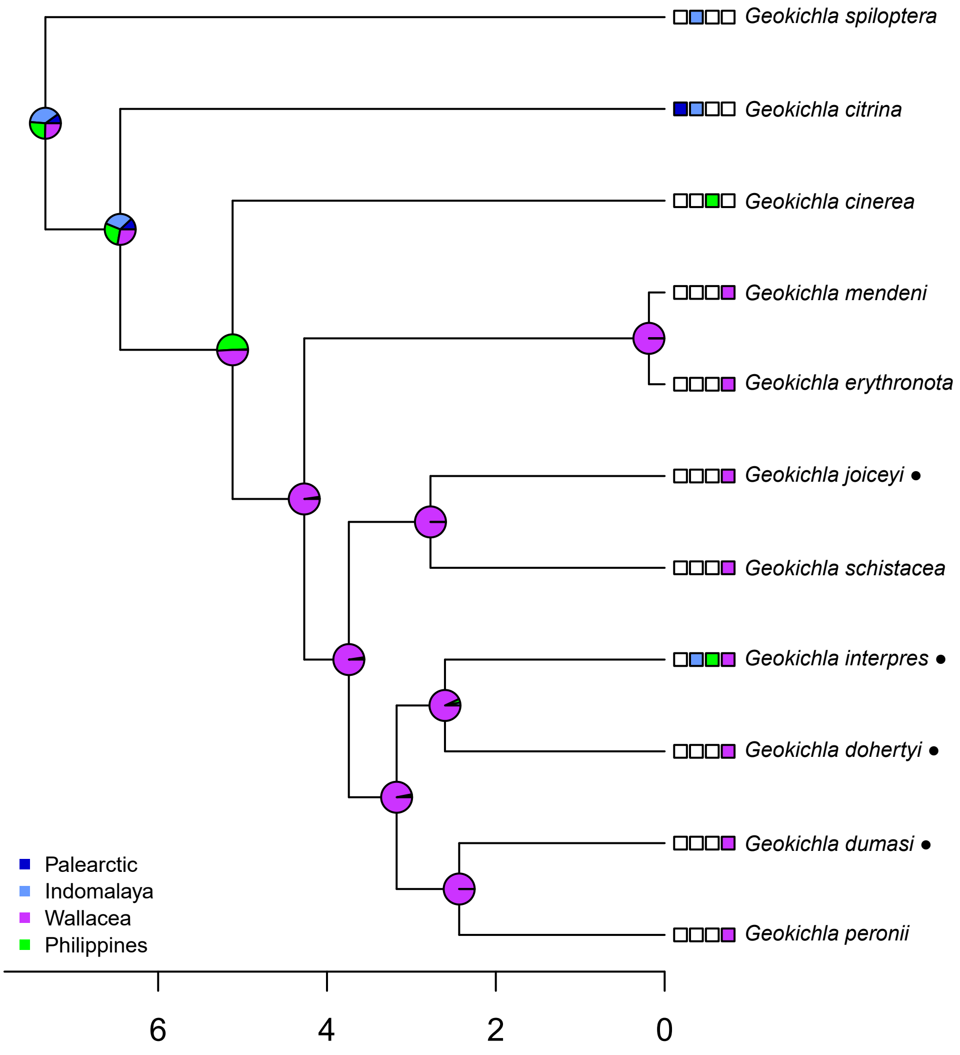

Geokichla spiloptera
Geokichla citrina
Geokichla cinerea
Geokichla mendeni
Geokichla erythronota
Geokichla joiceyi ●
Geokichla schistacea
Geokichla interpres ●
Geokichla dohertyi ●
Geokichla dumasi ●
Geokichla peronii
Palearctic
Indomalaya
Wallacea
Philippines
6
4
2
0

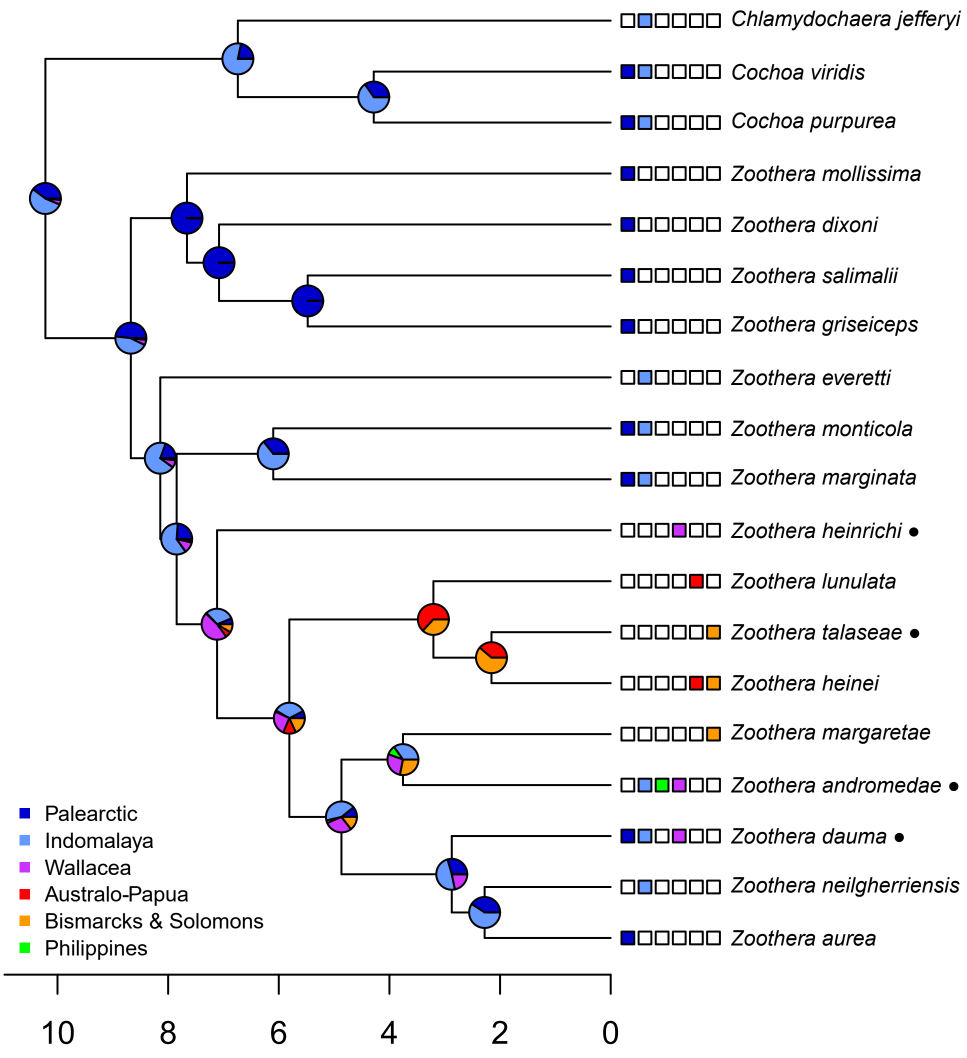

Chlamydochaera jefferyi
Cochoa viridis
Cochoa purpurea
Zoothera mollissima
Zoothera dixoni
Zoothera salimalii
Zoothera griseiceps
Zoothera everetti
Zoothera monticola
Zoothera marginata
Zoothera heinrichi ●
Zoothera lunulata
Zoothera talaseae ●
Zoothera heinei
Zoothera margaretae
Zoothera andromedae ●
Zoothera dauma ●
Zoothera neilgherriensis
Zoothera aurea
Palearctic
Indomalaya
Wallacea
Australo-Papua
Bismarcks & Solomons
Philippines
10
8
6
4
2
0

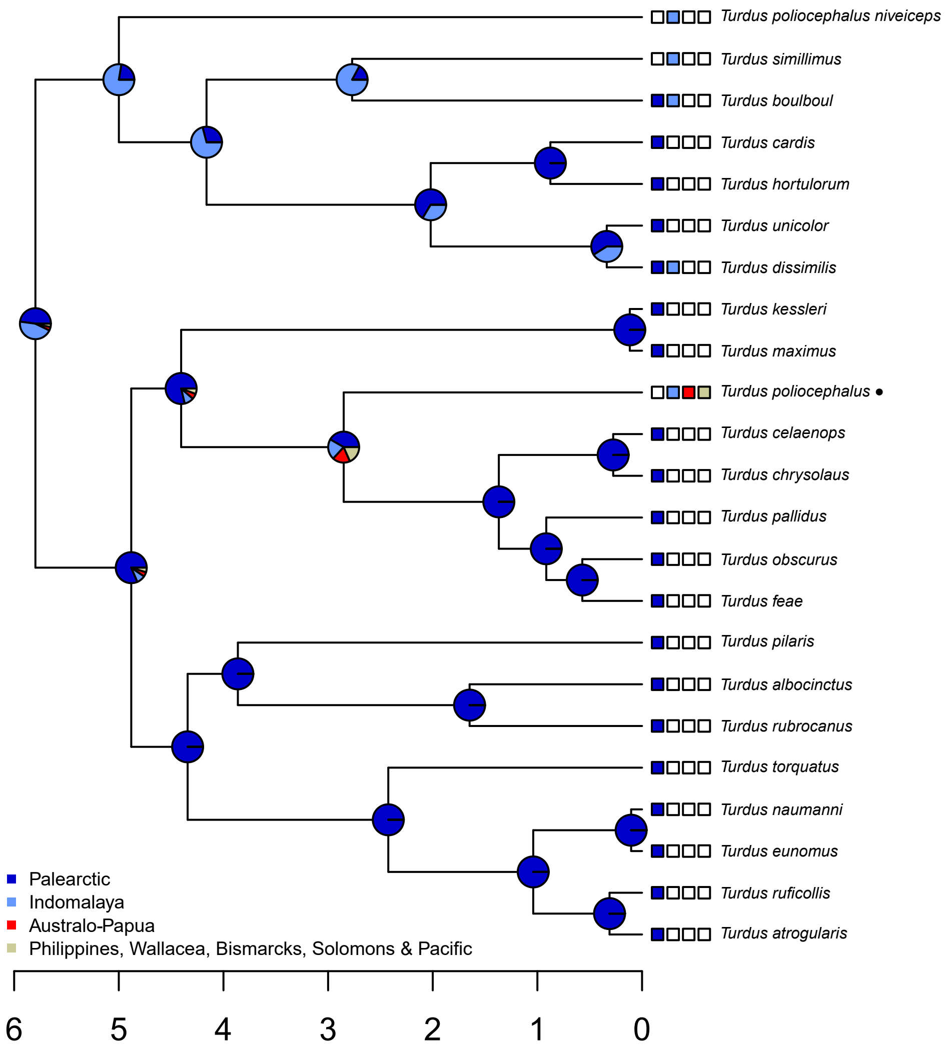

Turdus poliocephalus niveiceps
Turdus simillimus
Turdus boulboul
Turdus cardis
Turdus hortulorum
Turdus unicolor
Turdus dissimilis
Turdus kessleri
Turdus maximus
Turdus poliocephalus ●
Turdus celaenops
Turdus chrysolaus
Turdus pallidus
Turdus obscurus
Turdus feae
Turdus pilaris
Turdus albocinctus
Turdus rubrocanus
Turdus torquatus
Turdus naumanni
Turdus eunomus
Turdus ruficollis
Turdus atrogularis
Palearctic
Indomalaya
Australo-Papua
Philippines, Wallacea, Bismarcks, Solomons & Pacific
6
5
4
3
2
1
0

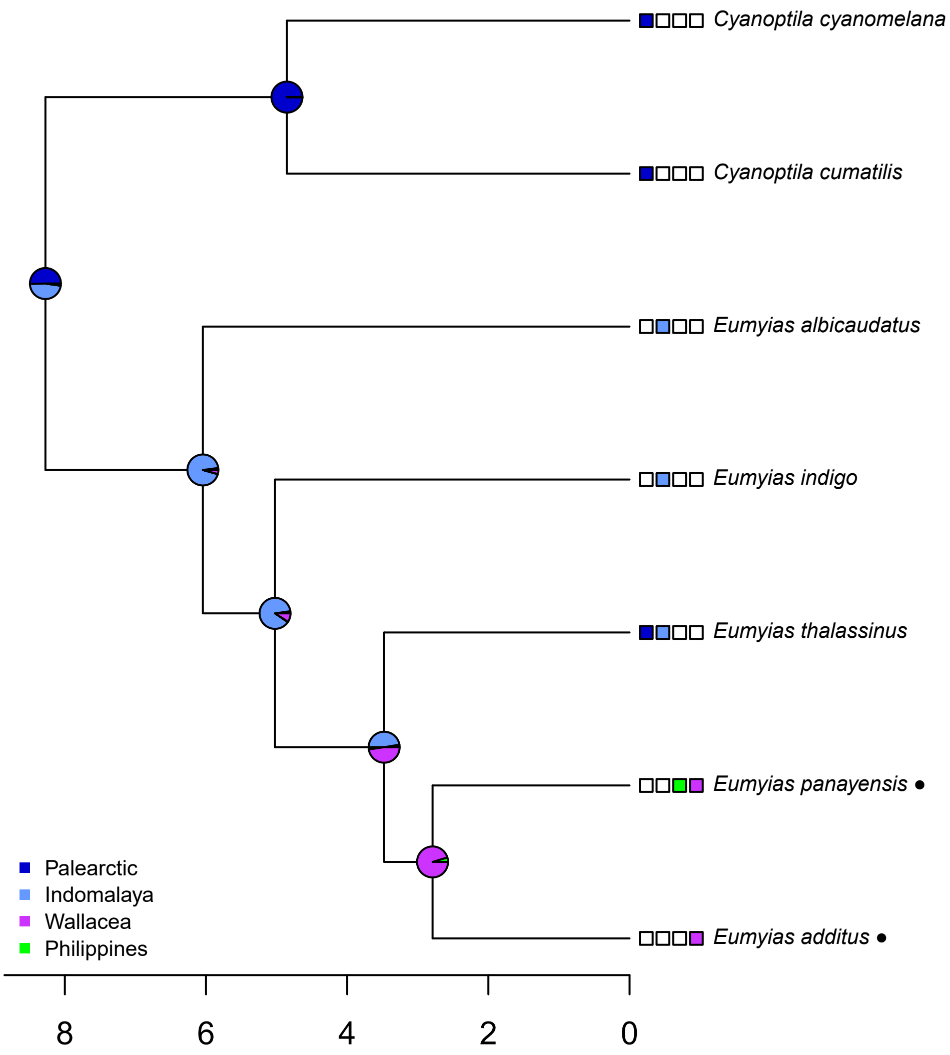

Cyanoptila cyanomelana
Cyanoptila cumatilis
Eumyias albicaudatus
Eumyias indigo
Eumyias thalassinus
Eumyias panayensis
Eumyias additus
Palearctic
Indomalaya
Wallacea
Philippines
8
6
4
2
0

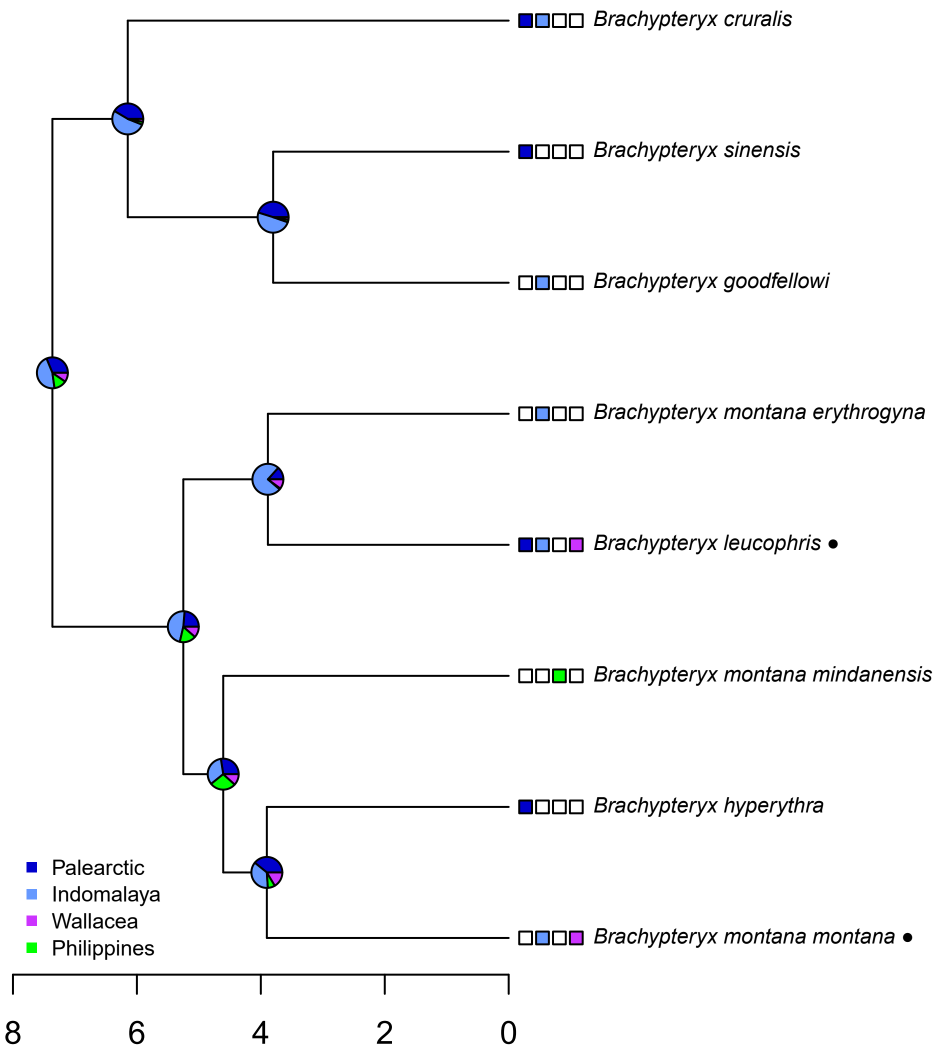

Brachypteryx cruralis
Brachypteryx sinensis
Brachypteryx goodfellowi
Brachypteryx montana erythrogyna
Brachypteryx leucophris ●
Brachypteryx montana mindanensis
Brachypteryx hyperythra
Brachypteryx montana montana ●
Palearctic
Indomalaya
Wallacea
Philippines
8
6
4
2
0

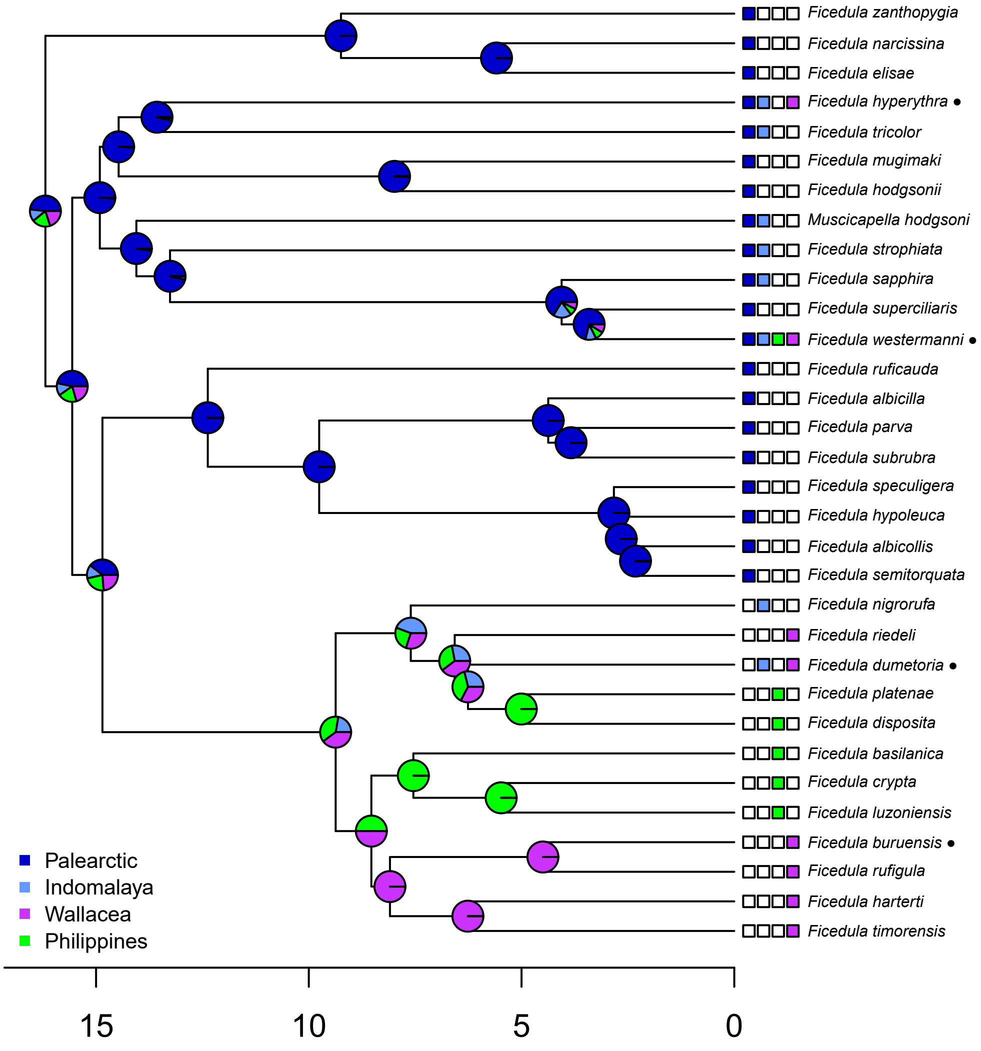

Ficedula zanthopygia
Ficedula narcissina
Ficedula elisae
Ficedula hyperythra •
Ficedula tricolor
Ficedula mugimaki
Ficedula hodgsonii
Muscicapella hodgsoni
Ficedula strophiata
Ficedula sapphira
Ficedula superciliaris
Ficedula westermanni •
Ficedula ruficauda
Ficedula albicilla
Ficedula parva
Ficedula subrubra
Ficedula speculigera
Ficedula hypoleuca
Ficedula albicollis
Ficedula semitorquata
Ficedula nigrorufa
Ficedula riedeli
Ficedula dumetoria •
Ficedula platenae
Ficedula disposita
Ficedula basilanica
Ficedula crypta
Ficedula luzoniensis
Ficedula buruensis •
Ficedula rufigula
Ficedula harterti
Ficedula timorensis
Palearctic
Indomalaya
Wallacea
Philippines
15
10
5
0

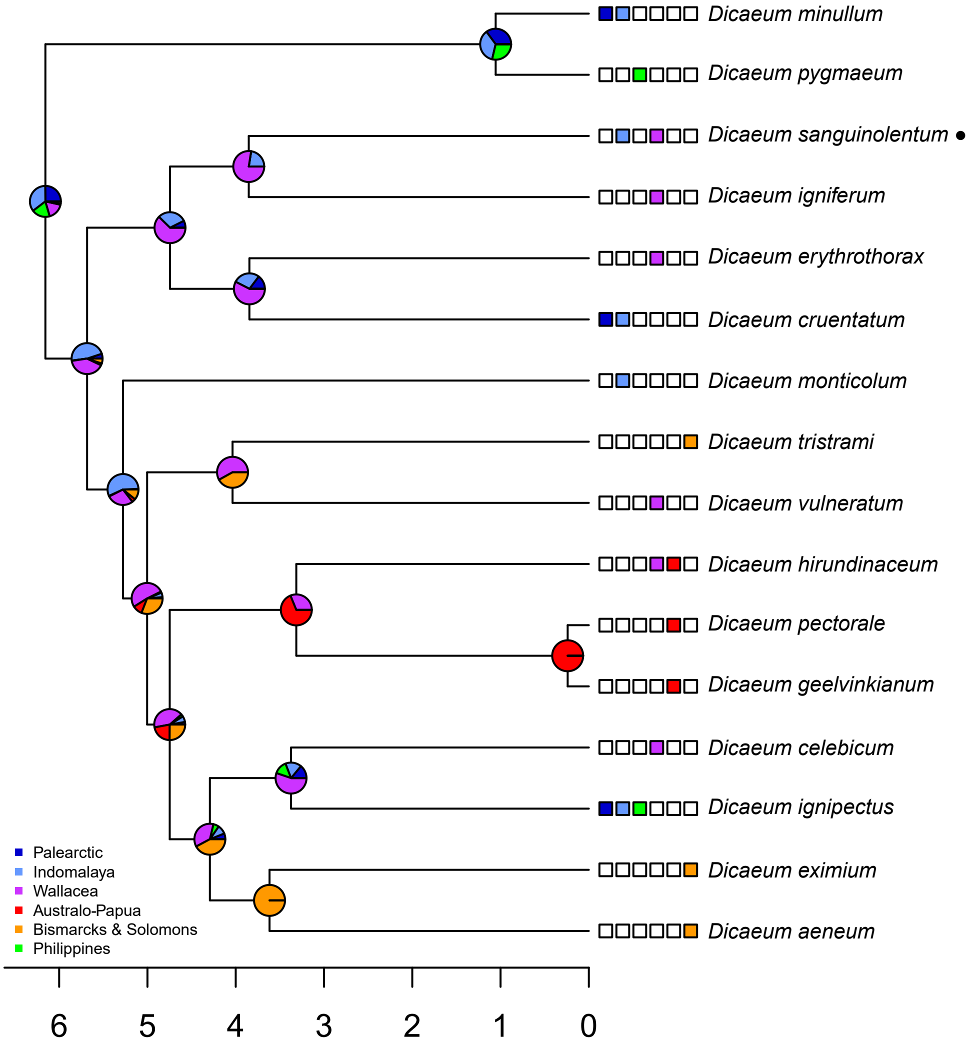

Dicaeum minullum
Dicaeum pygmaeum
Dicaeum sanguinolentum
Dicaeum igniferum
Dicaeum erythrothorax
Dicaeum cruentatum
Dicaeum monticolum
Dicaeum tristrami
Dicaeum vulneratum
Dicaeum hirundinaceum
Dicaeum pectorale
Dicaeum geelvinkianum
Dicaeum celebicum
Dicaeum ignipectus
Dicaeum eximium
Dicaeum aeneum
Palearctic
Indomalaya
Wallacea
Australo-Papua
Bismarcks & Solomons
Philippines
6
5
4
3
2
1
0

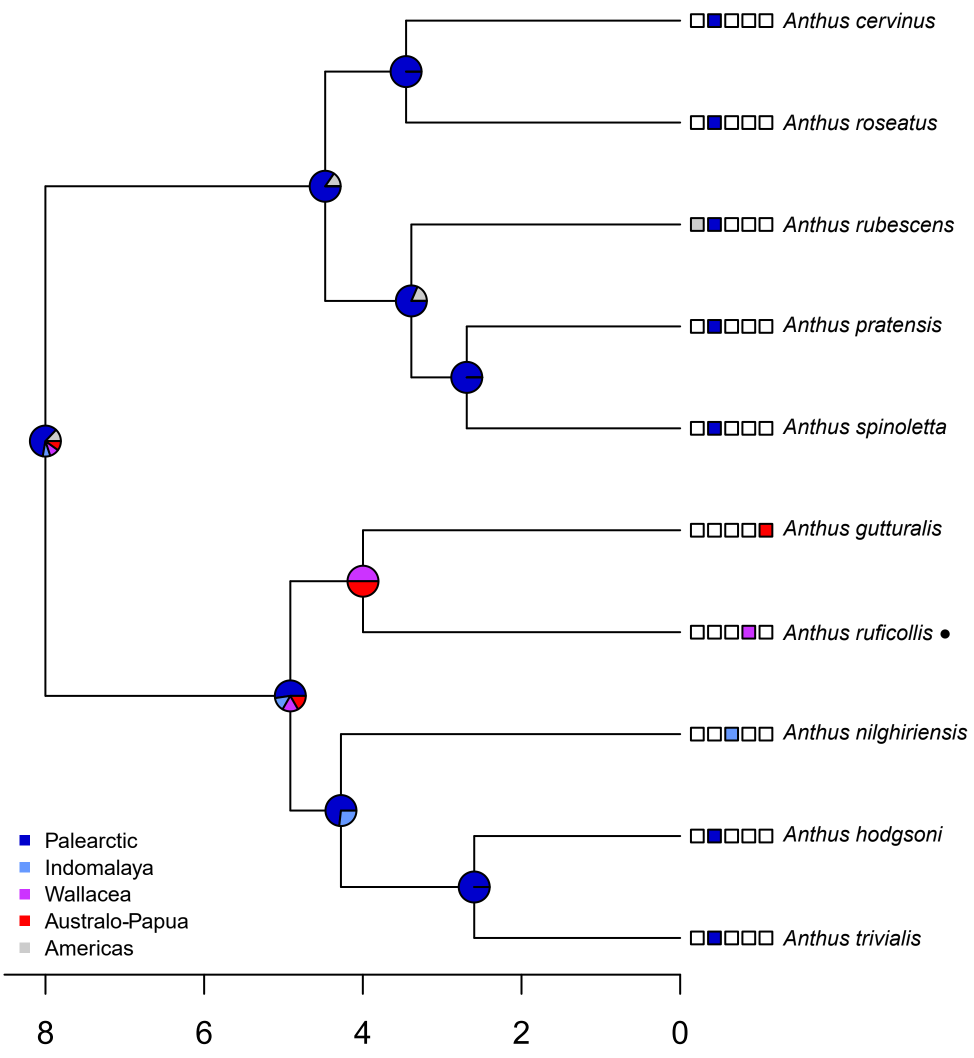

Anthus cervinus
Anthus roseatus
Anthus rubescens
Anthus pratensis
Anthus spinoletta
Anthus gutturalis
Anthus ruficollis •
Anthus nilghiriensis
Anthus hodgsoni
Anthus trivialis
Palearctic
Indomalaya
Wallacea
Australo-Papua
Americas
8
6
4
2
0

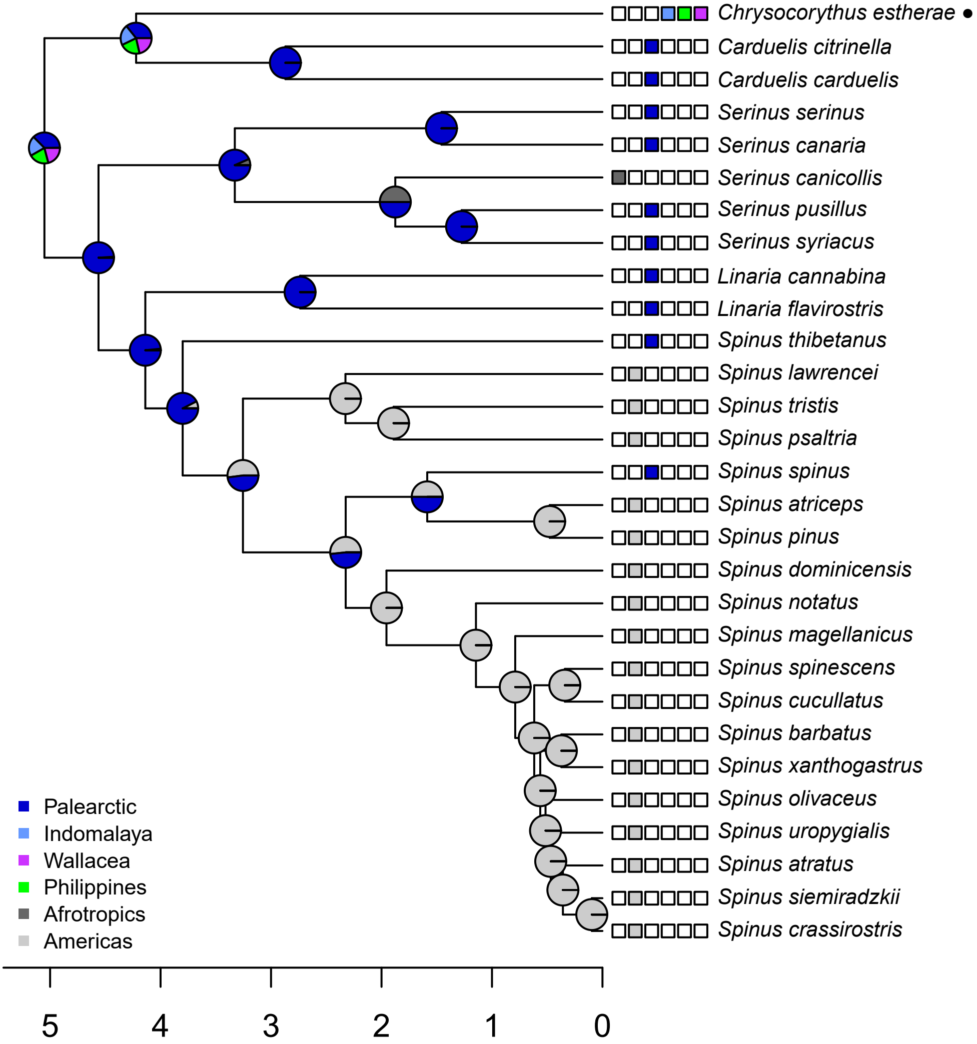
Chrysocorythus estherae ●
Carduelis citrinella
Carduelis carduelis
Serinus serinus
Serinus canaria
Serinus canicollis
Serinus pusillus
Serinus syriacus
Linaria cannabina
Linaria flavirostris
Spinus thibetanus
Spinus lawrencei
Spinus tristis
Spinus psaltria
Spinus spinus
Spinus atriceps
Spinus pinus
Spinus dominicensis
Spinus notatus
Spinus magellanicus
Spinus spinescens
Spinus cucullatus
Spinus barbatus
Spinus xanthogastrus
Spinus olivaceus
Spinus uropygialis
Spinus atratus
Spinus siemiradzkii
Spinus crassirostris
Palearctic
Indomalaya
Wallacea
Philippines
Afrotropics
Americas
5
4
3
2
1
0

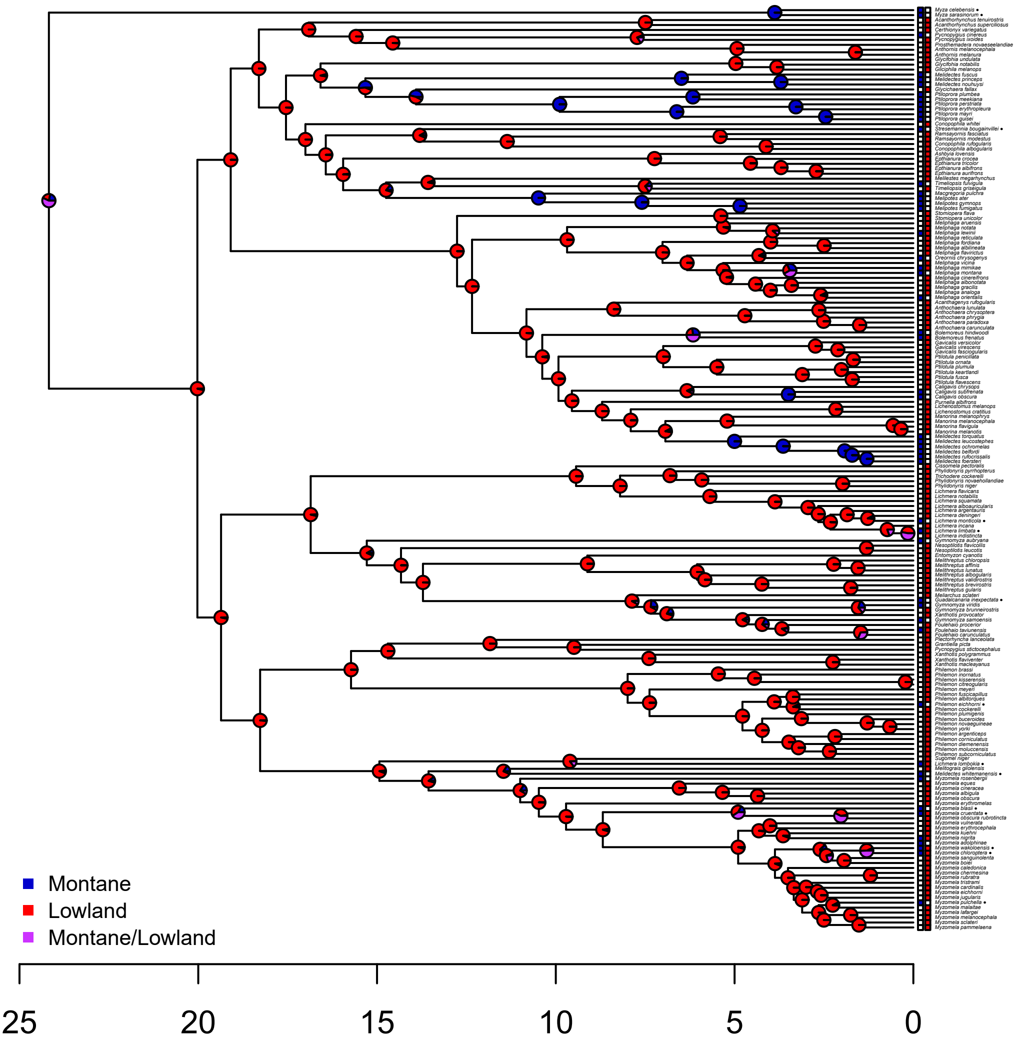

Montane
Lowland
Montane/Lowland
25
20
15
10
5
0

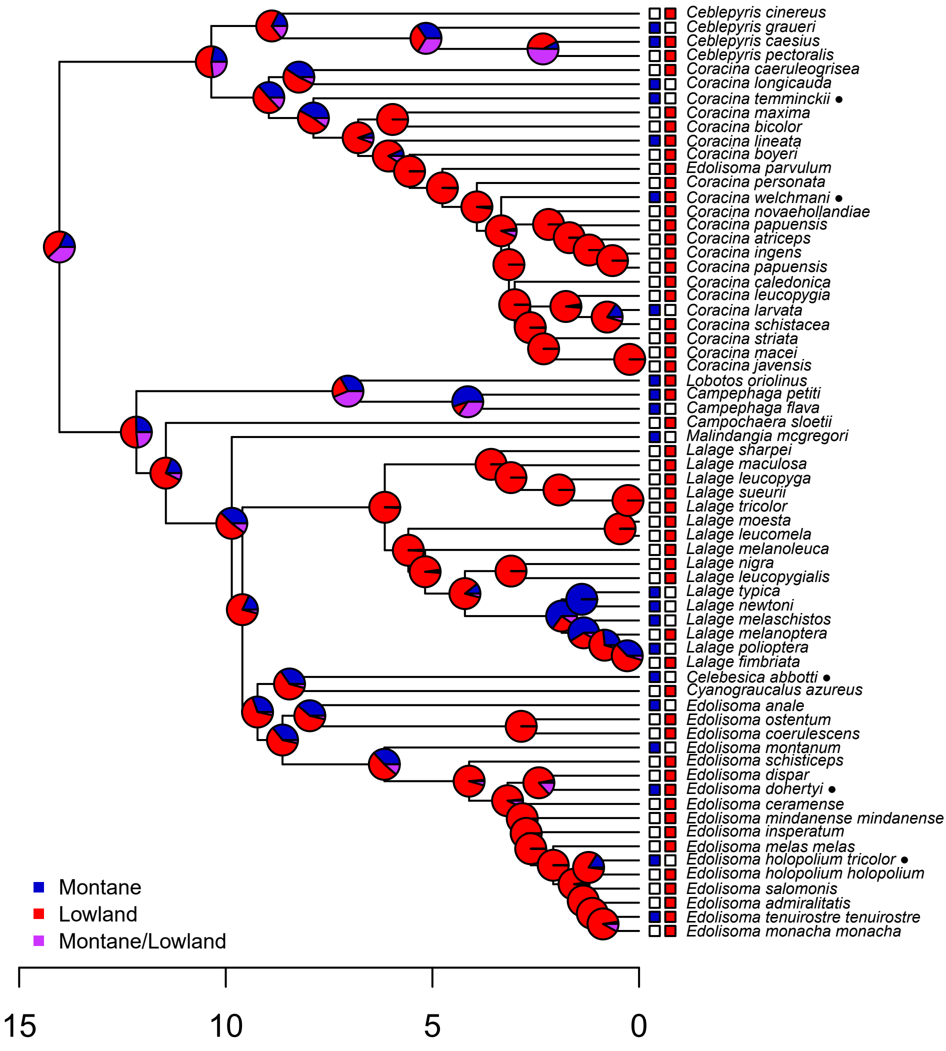

Ceblepyris cinereus
Ceblepyris graueri
Ceblepyris caesius
Ceblepyris pectoralis
Coracina caeruleogrisea
Coracina longicauda
Coracina temminckii ●
Coracina maxima
Coracina bicolor
Coracina lineata
Coracina boyeri
Edolisoma parvulum
Coracina personata
Coracina welchmani ●
Coracina novaehollandiae
Coracina papuensis
Coracina atriceps
Coracina ingens
Coracina papuensis
Coracina caledonica
Coracina leucopygia
Coracina larvata
Coracina schistacea
Coracina striata
Coracina macei
Coracina javensis
Lobotos oriolinus
Campephaga petiti
Campephaga flava
Campochaera sloetii
Malindangia mcgregori
Lalage sharpei
Lalage maculosa
Lalage leucopyga
Lalage sueurii
Lalage tricolor
Lalage moesta
Lalage leucomela
Lalage melanoleuca
Lalage nigra
Lalage leucopygialis
Lalage typica
Lalage newtoni
Lalage melaschistos
Lalage melanoptera
Lalage polioptera
Lalage fimbriata
Celebesica abbotti ●
Cyanograucalus azureus
Edolisoma anale
Edolisoma ostentum
Edolisoma coerulescens
Edolisoma montanum
Edolisoma schisticeps
Edolisoma dispar
Edolisoma dohertyi ●
Edolisoma ceramense
Edolisoma mindanense mindanense
Edolisoma insperatum
Edolisoma melas melas
Edolisoma holopolium tricolor ●
Edolisoma holopolium holopolium
Edolisoma salomonis
Edolisoma admiralitatis
Edolisoma tenuirostre tenuirostre
Edolisoma monacha monacha
Montane
Lowland
Montane/Lowland
15
10
5
0

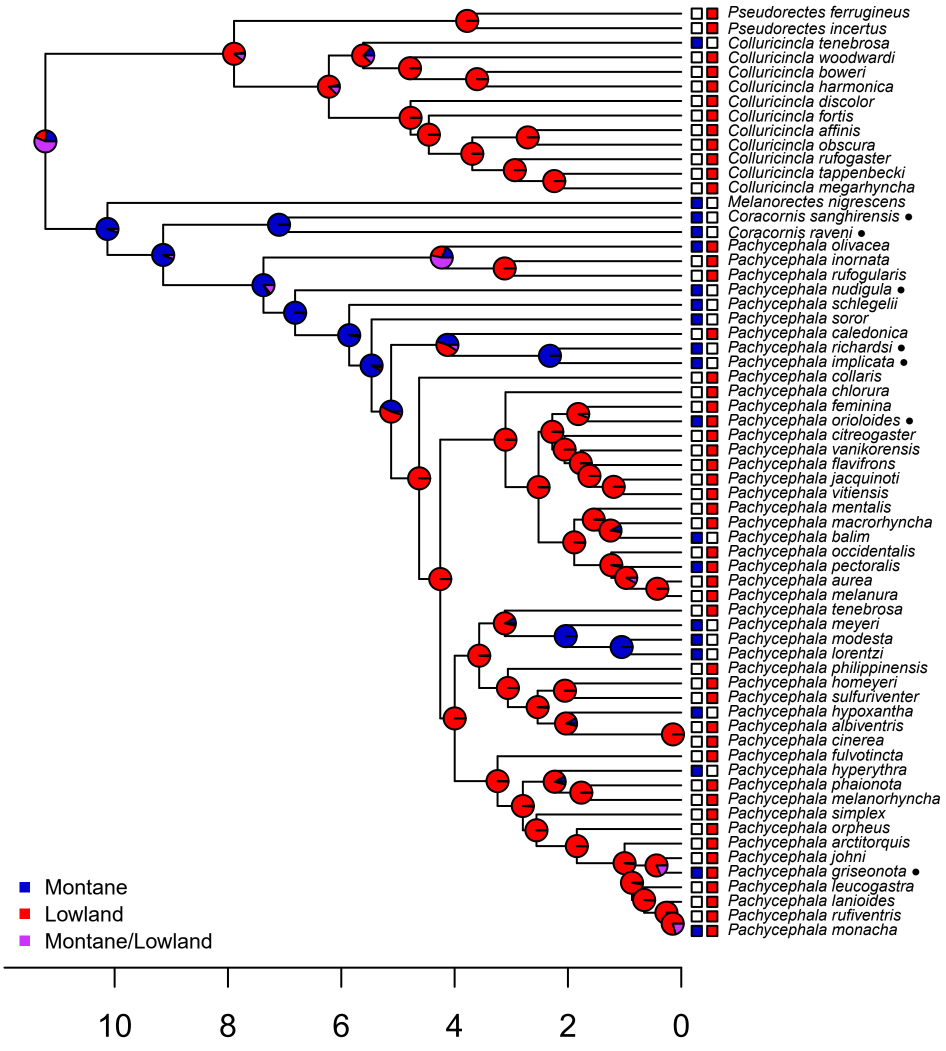

Pseudorectes ferrugineus
Pseudorectes incertus
Colluricincla tenebrosa
Colluricincla woodwardi
Colluricincla boweri
Colluricincla harmonica
Colluricincla discolor
Colluricincla fortis
Colluricincla affinis
Colluricincla obscura
Colluricincla rufogaster
Colluricincla tappenbecki
Colluricincla megarhyncha
Melanorectes nigrescens
Coracornis sanghirensis ●
Coracornis raveni ●
Pachycephala olivacea
Pachycephala inornata
Pachycephala rufogularis
Pachycephala nudigula ●
Pachycephala schlegelii
Pachycephala soror
Pachycephala caledonica
Pachycephala richardsi ●
Pachycephala implicata ●
Pachycephala collaris
Pachycephala chlorura
Pachycephala feminina
Pachycephala orioloides ●
Pachycephala citreogaster
Pachycephala vanikorensis
Pachycephala flavifrons
Pachycephala jacquinoti
Pachycephala vitiensis
Pachycephala mentalis
Pachycephala macrorhyncha
Pachycephala balim
Pachycephala occidentalis
Pachycephala pectoralis
Pachycephala aurea
Pachycephala melanura
Pachycephala tenebrosa
Pachycephala meyeri
Pachycephala modesta
Pachycephala lorentzi
Pachycephala philippinensis
Pachycephala homeyeri
Pachycephala sulfuriventer
Pachycephala hypoxantha
Pachycephala albiventris
Pachycephala cinerea
Pachycephala fulvotincta
Pachycephala hyperythra
Pachycephala phaionota
Pachycephala melanorhyncha
Pachycephala simplex
Pachycephala orpheus
Pachycephala arctitorquis
Pachycephala johni
Pachycephala griseonota ●
Pachycephala leucogastra
Pachycephala lanioides
Pachycephala rufiventris
Pachycephala monacha
Montane
Lowland
Montane/Lowland
10
8
6
4
2
0

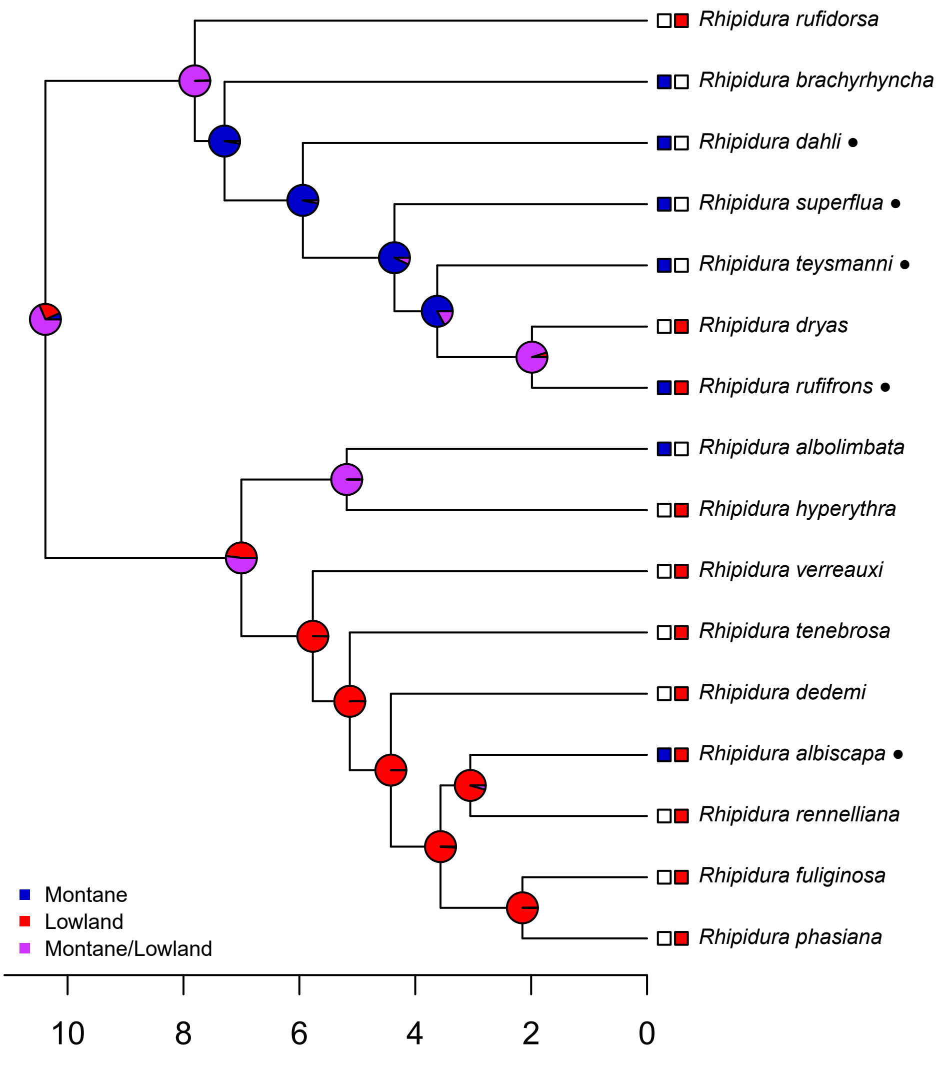

Rhipidura rufidorsa
Rhipidura brachyrhyncha
Rhipidura dahli ●
Rhipidura superflua ●
Rhipidura teysmanni ●
Rhipidura dryas
Rhipidura rufifrons ●
Rhipidura albolimbata
Rhipidura hyperythra
Rhipidura verreauxi
Rhipidura tenebrosa
Rhipidura dedemi
Rhipidura albiscapa ●
Rhipidura rennelliana
Rhipidura fuliginosa
Rhipidura phasiana
Montane
Lowland
Montane/Lowland
10
8
6
4
2
0

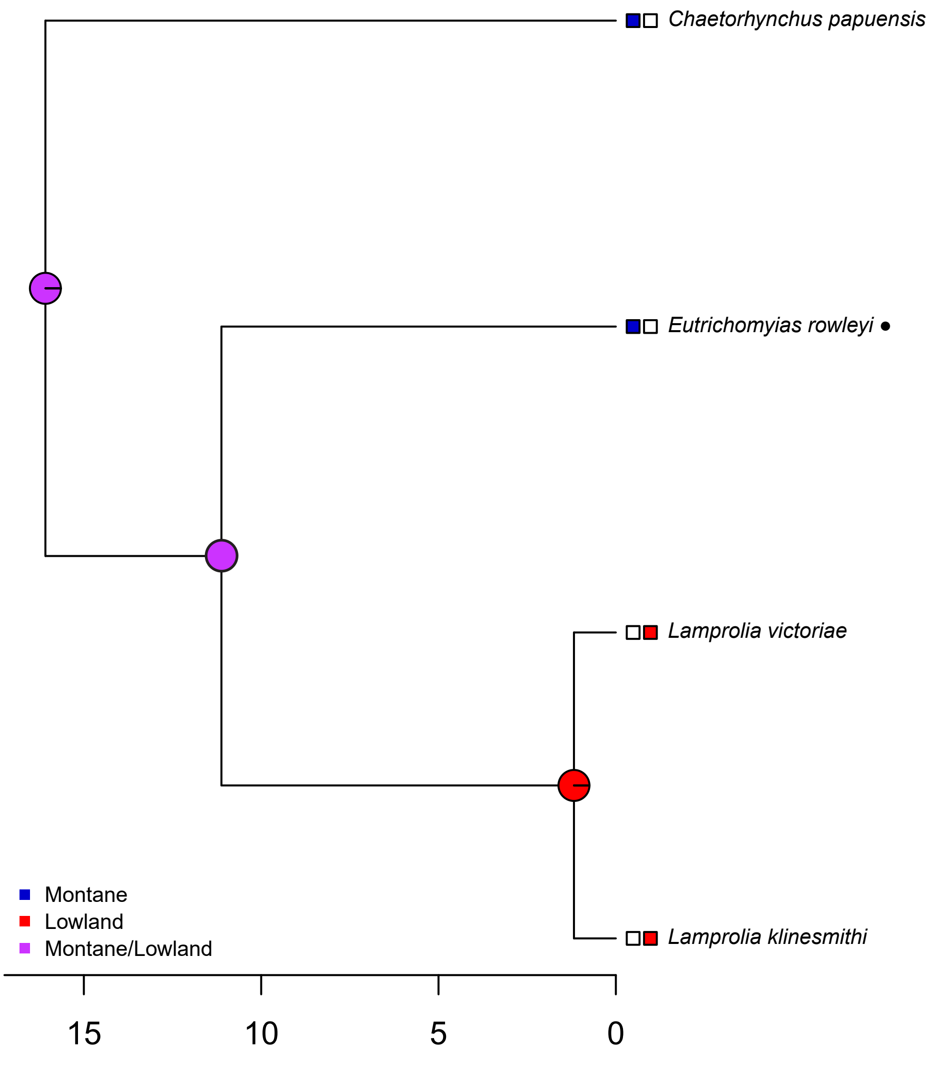

Chaetorhynchus papuensis
Eutrichomyias rowleyi ●
Lamprolia victoriae
Lamprolia klinesmithi
Montane
Lowland
Montane/Lowland
15
10
5
0

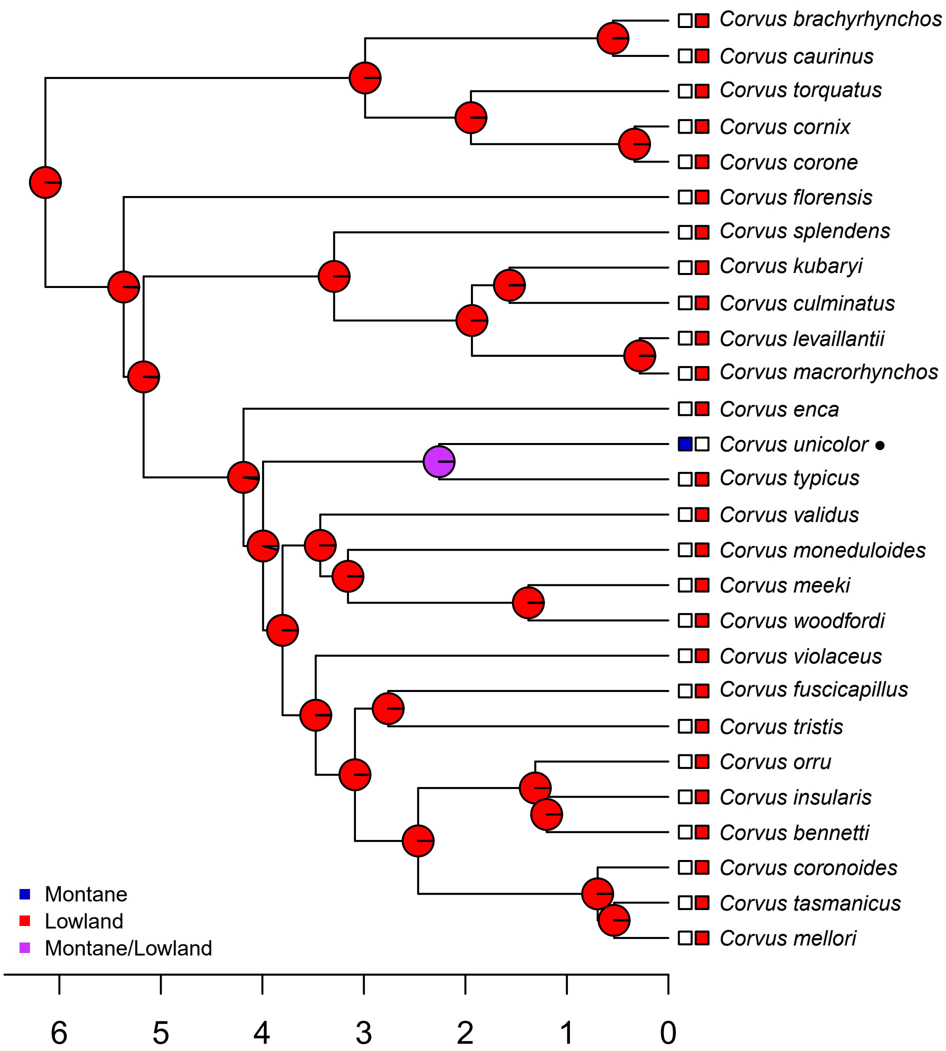
Corvus brachyrhynchos
Corvus caurinus
Corvus torquatus
Corvus cornix
Corvus corone
Corvus florensis
Corvus splendens
Corvus kubaryi
Corvus culminatus
Corvus levaillantii
Corvus macrorhynchos
Corvus enca
Corvus unicolor ●
Corvus typicus
Corvus validus
Corvus moneduloides
Corvus meeki
Corvus woodfordi
Corvus violaceus
Corvus fuscicapillus
Corvus tristis
Corvus orru
Corvus insularis
Corvus bennetti
Corvus coronoides
Corvus tasmanicus
Corvus mellori
Montane
Lowland
Montane/Lowland
6
5
4
3
2
1
0

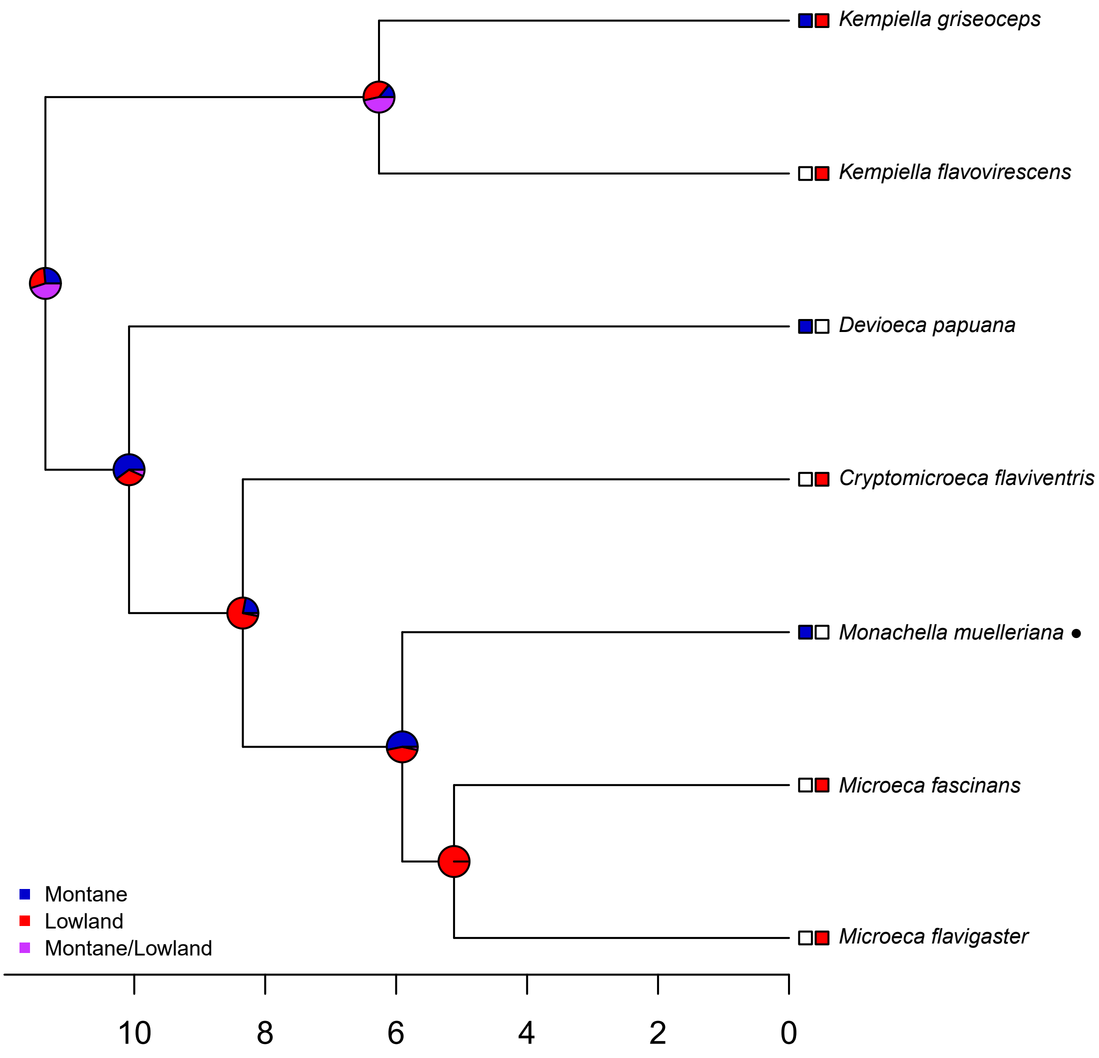

Kempiella griseoceps
Kempiella flavovirescens
Devioeca papuana
Cryptomicroeca flaviventris
Monachella muelleriana ●
Microeca fascinans
Microeca flavigaster
Montane
Lowland
Montane/Lowland
10
8
6
4
2
0

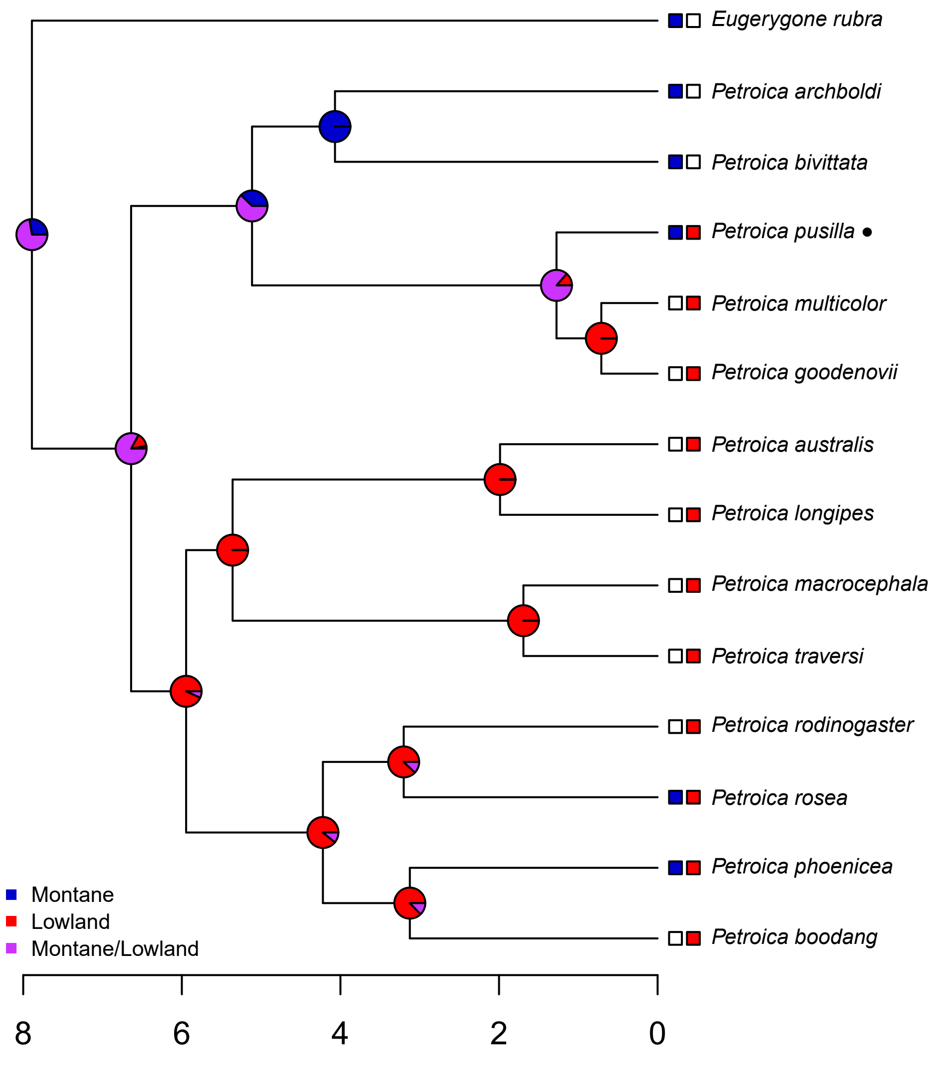

Eugerygone rubra
Petroica archboldi
Petroica bivittata
Petroica pusilla ●
Petroica multicolor
Petroica goodenovii
Petroica australis
Petroica longipes
Petroica macrocephala
Petroica traversi
Petroica rodinogaster
Petroica rosea
Petroica phoenicea
Petroica boodang
Montane
Lowland
Montane/Lowland
8
6
4
2
0

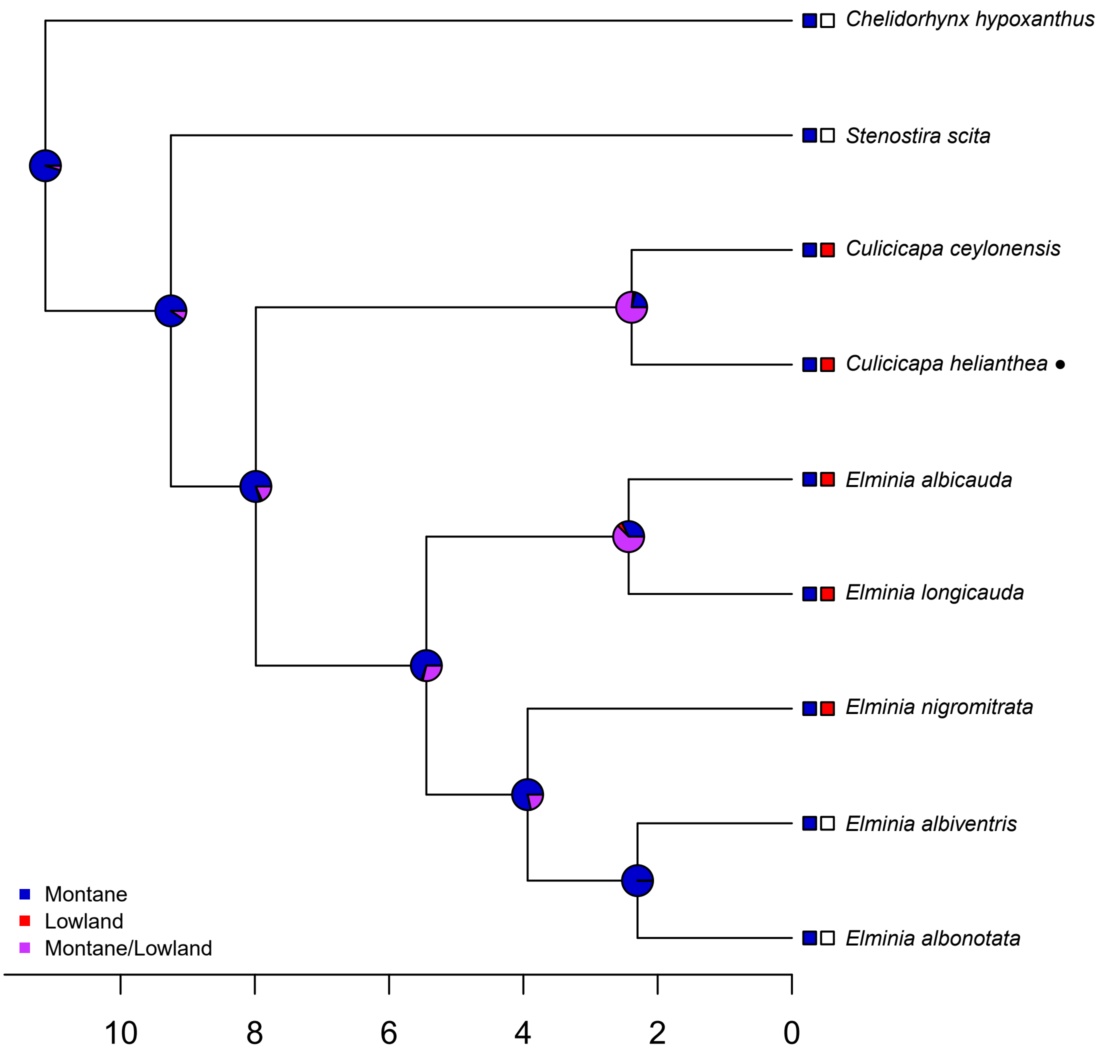
Chelidorhynx hypoxanthus
Stenostira scita
Culicicapa ceylonensis
Culicicapa helianthea ●
Elminia albicauda
Elminia longicauda
Elminia nigromitrata
Elminia albiventris
Elminia albonotata
Montane
Lowland
Montane/Lowland
10
8
6
4
2
0

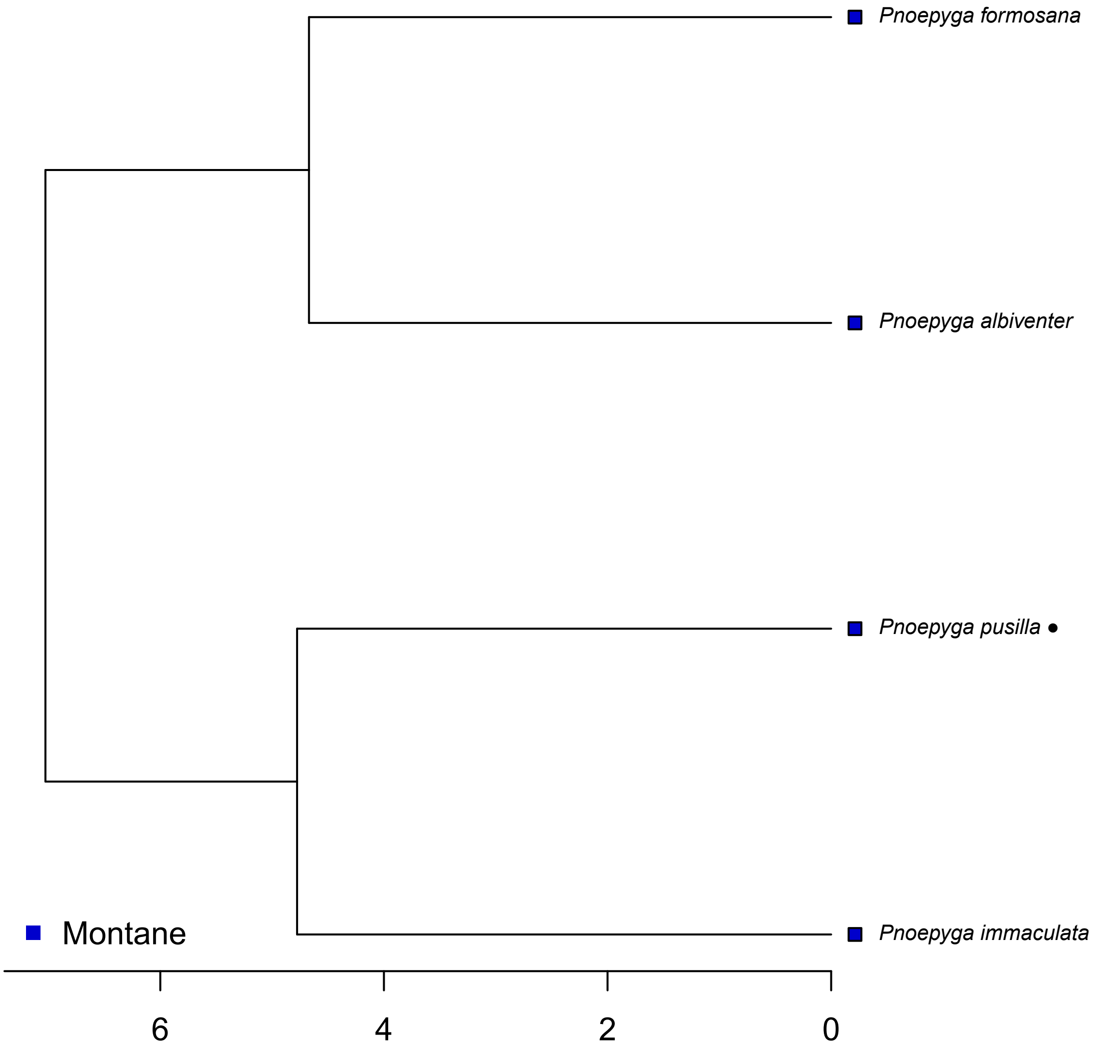

Pnoepyga formosana
Pnoepyga albiventer
Pnoepyga pusilla ●
Pnoepyga immaculata
Montane
6
4
2
0

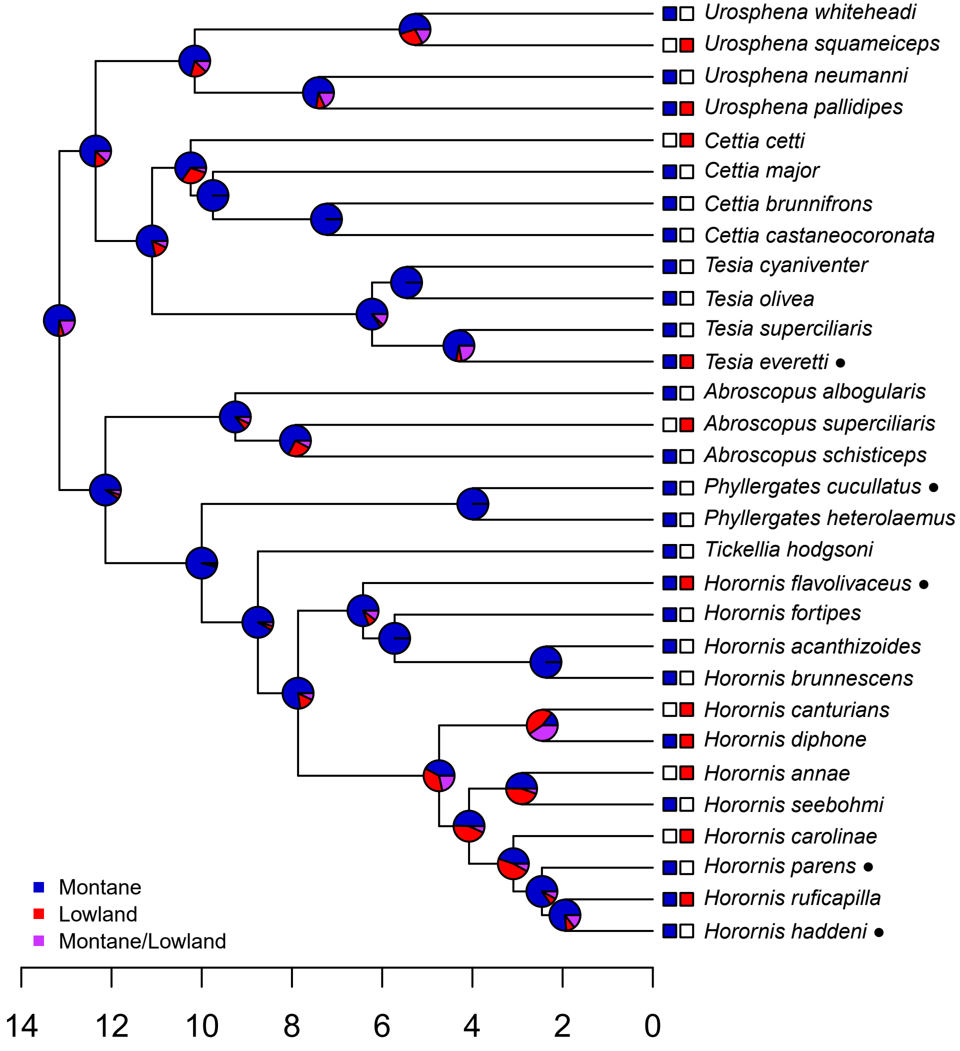

Urosphena whiteheadi
Urosphena squameiceps
Urosphena neumanni
Urosphena pallidipes
Cettia cetti
Cettia major
Cettia brunnifrons
Cettia castaneocoronata
Tesia cyaniventer
Tesia olivea
Tesia superciliaris
Tesia everetti ●
Abroscopus albogularis
Abroscopus superciliaris
Abroscopus schisticeps
Phyllergates cucullatus ●
Phyllergates heterolaemus
Tickellia hodgsoni
Horornis flavolivaceus ●
Horornis fortipes
Horornis acanthizoides
Horornis brunnescens
Horornis canturians
Horornis diphone
Horornis annae
Horornis seebohmi
Horornis carolinae
Horornis parens ●
Horornis ruficapilla
Horornis haddeni ●
Montane
Lowland
Montane/Lowland
14
12
10
8
6
4
2
0

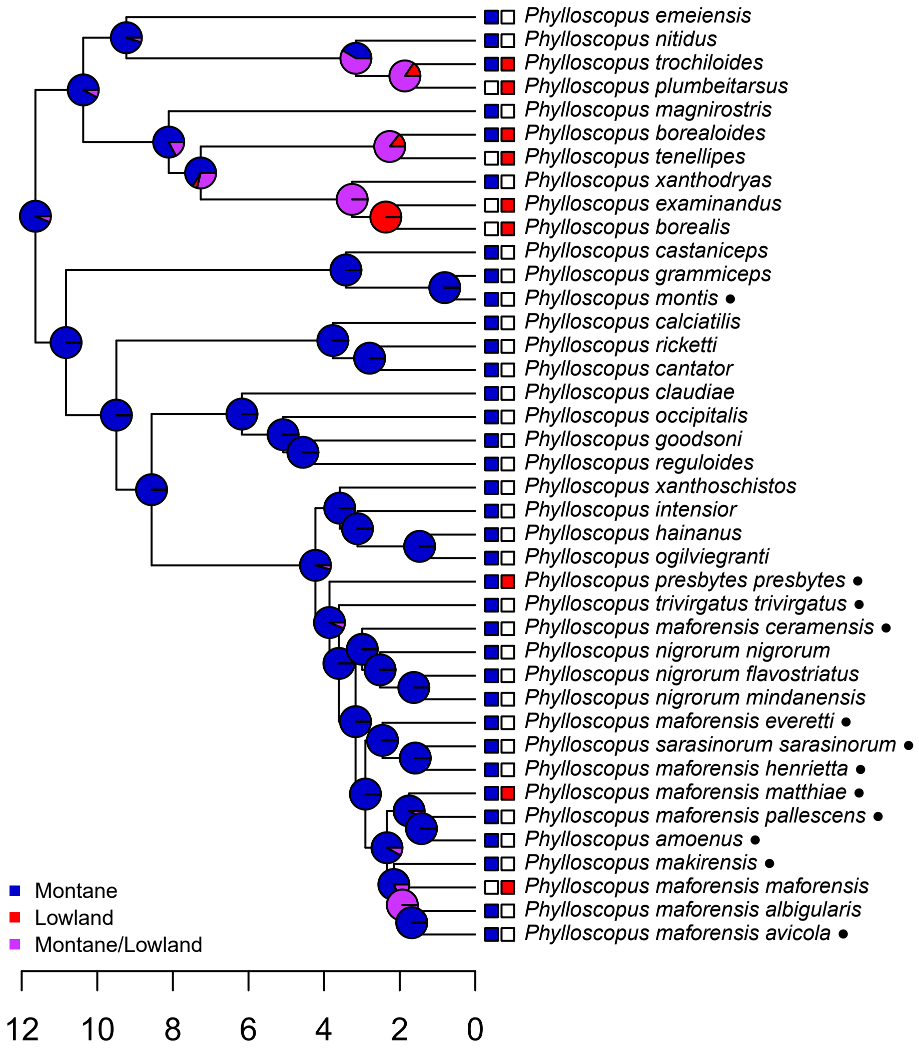

Phylloscopus emeiensis
Phylloscopus nitidus
Phylloscopus trochiloides
Phylloscopus plumbeitarsus
Phylloscopus magnirostris
Phylloscopus borealoides
Phylloscopus tenellipes
Phylloscopus xanthodryas
Phylloscopus examinandus
Phylloscopus borealis
Phylloscopus castaniceps
Phylloscopus grammiceps
Phylloscopus montis ●
Phylloscopus calciatilis
Phylloscopus ricketti
Phylloscopus cantator
Phylloscopus claudiae
Phylloscopus occipitalis
Phylloscopus goodsoni
Phylloscopus reguloides
Phylloscopus xanthoschistos
Phylloscopus intensior
Phylloscopus hainanus
Phylloscopus ogilviegranti
Phylloscopus presbytes presbytes ●
Phylloscopus trivirgatus trivirgatus ●
Phylloscopus maforensis ceramensis ●
Phylloscopus nigrorum nigrorum
Phylloscopus nigrorum flavostriatus
Phylloscopus nigrorum mindanensis
Phylloscopus maforensis everetti ●
Phylloscopus sarasinorum sarasinorum ●
Phylloscopus maforensis henrietta ●
Phylloscopus maforensis matthiae ●
Phylloscopus maforensis pallescens ●
Phylloscopus amoenus ●
Phylloscopus makirensis ●
Phylloscopus maforensis maforensis
Phylloscopus maforensis albigularis
Phylloscopus maforensis avicola ●
Montane
Lowland
Montane/Lowland
12
10
8
6
4
2
0

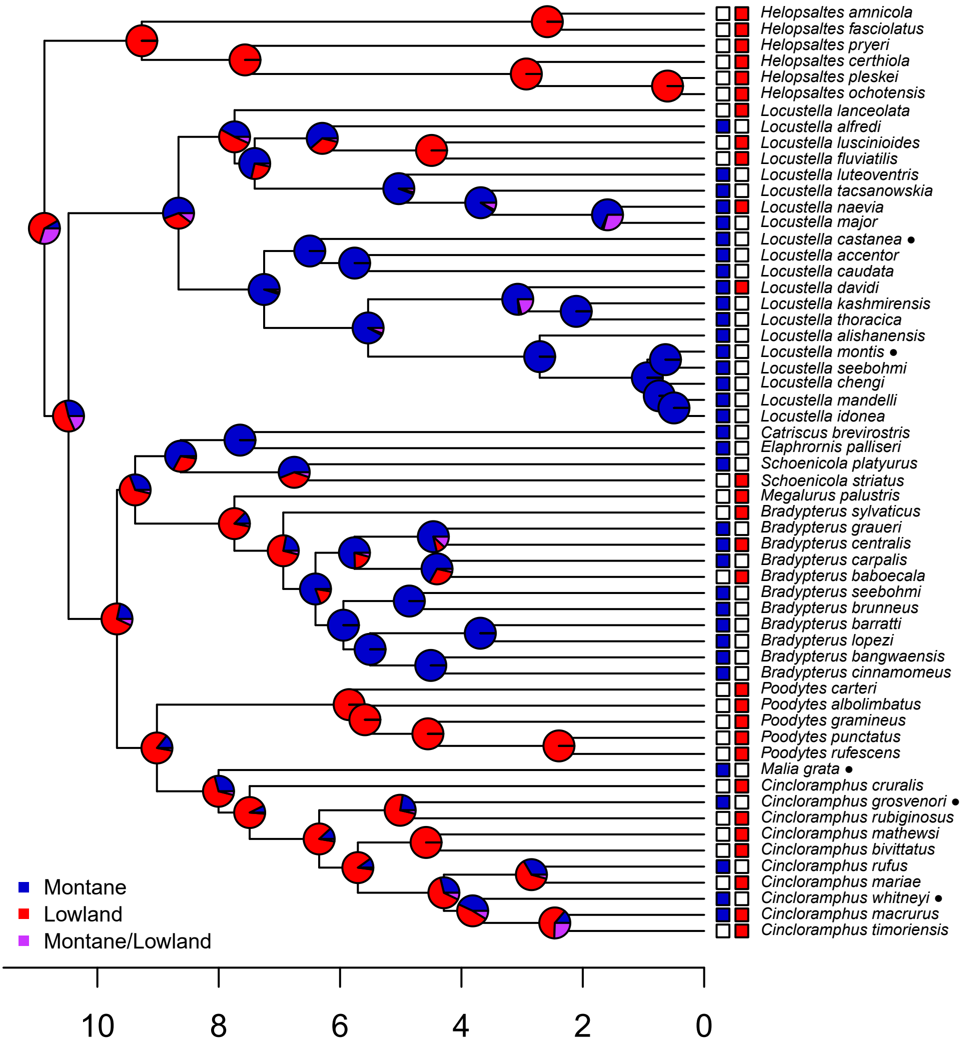
Helopsaltes amnicola
Helopsaltes fasciolatus
Helopsaltes pryeri
Helopsaltes certhiola
Helopsaltes pleskei
Helopsaltes ochotensis
Locustella lanceolata
Locustella alfredi
Locustella luscinioides
Locustella fluviatilis
Locustella luteoventris
Locustella tacsanowskia
Locustella naevia
Locustella major
Locustella castanea ●
Locustella accentor
Locustella caudata
Locustella davidi
Locustella kashmirensis
Locustella thoracica
Locustella alishanensis
Locustella montis ●
Locustella seebohmi
Locustella chengi
Locustella mandelli
Locustella idonea
Catriscus brevirostris
Elaphrornis palliseri
Schoenicola platyurus
Schoenicola striatus
Megalurus palustris
Bradypterus sylvaticus
Bradypterus graueri
Bradypterus centralis
Bradypterus carpalis
Bradypterus baboecala
Bradypterus seebohmi
Bradypterus brunneus
Bradypterus barratti
Bradypterus lopezi
Bradypterus bangwaensis
Bradypterus cinnamomeus
Poodytes carteri
Poodytes albolimbatus
Poodytes gramineus
Poodytes punctatus
Poodytes rufescens
Malia grata ●
Cincloramphus cruralis
Cincloramphus grosvenori ●
Cincloramphus rubiginosus
Cincloramphus mathewsi
Cincloramphus bivittatus
Cincloramphus rufus
Cincloramphus mariae
Cincloramphus whitneyi ●
Cincloramphus macrurus
Cincloramphus timoriensis
Montane
Lowland
Montane/Lowland
10
8
6
4
2
0

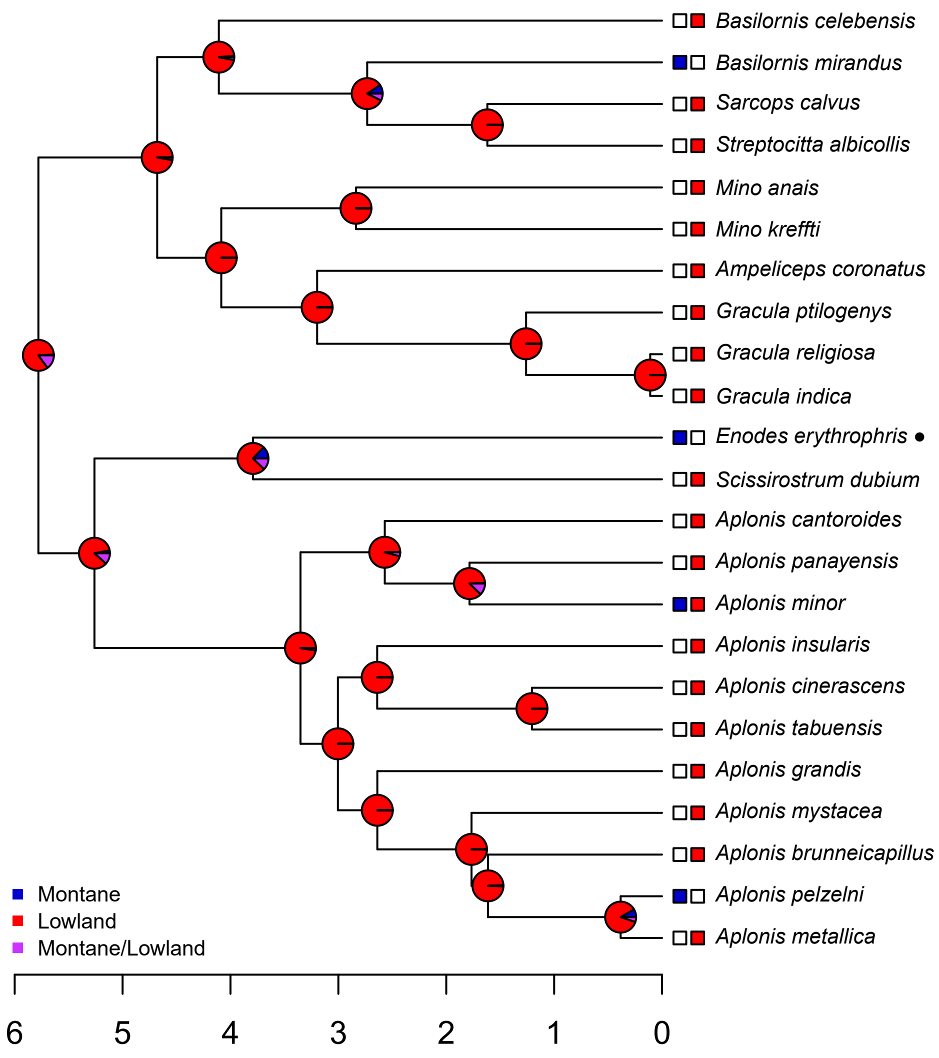
Basilornis celebensis
Basilornis mirandus
Sarcops calvus
Streptocitta albicollis
Mino anais
Mino kreffti
Ampeliceps coronatus
Gracula ptilogenys
Gracula religiosa
Gracula indica
Enodes erythrophris ●
Scissirostrum dubium
Aplonis cantoroides
Aplonis panayensis
Aplonis minor
Aplonis insularis
Aplonis cinerascens
Aplonis tabuensis
Aplonis grandis
Aplonis mystacea
Aplonis brunneicapillus
Aplonis pelzelni
Aplonis metallica
Montane
Lowland
Montane/Lowland
6
5
4
3
2
1
0

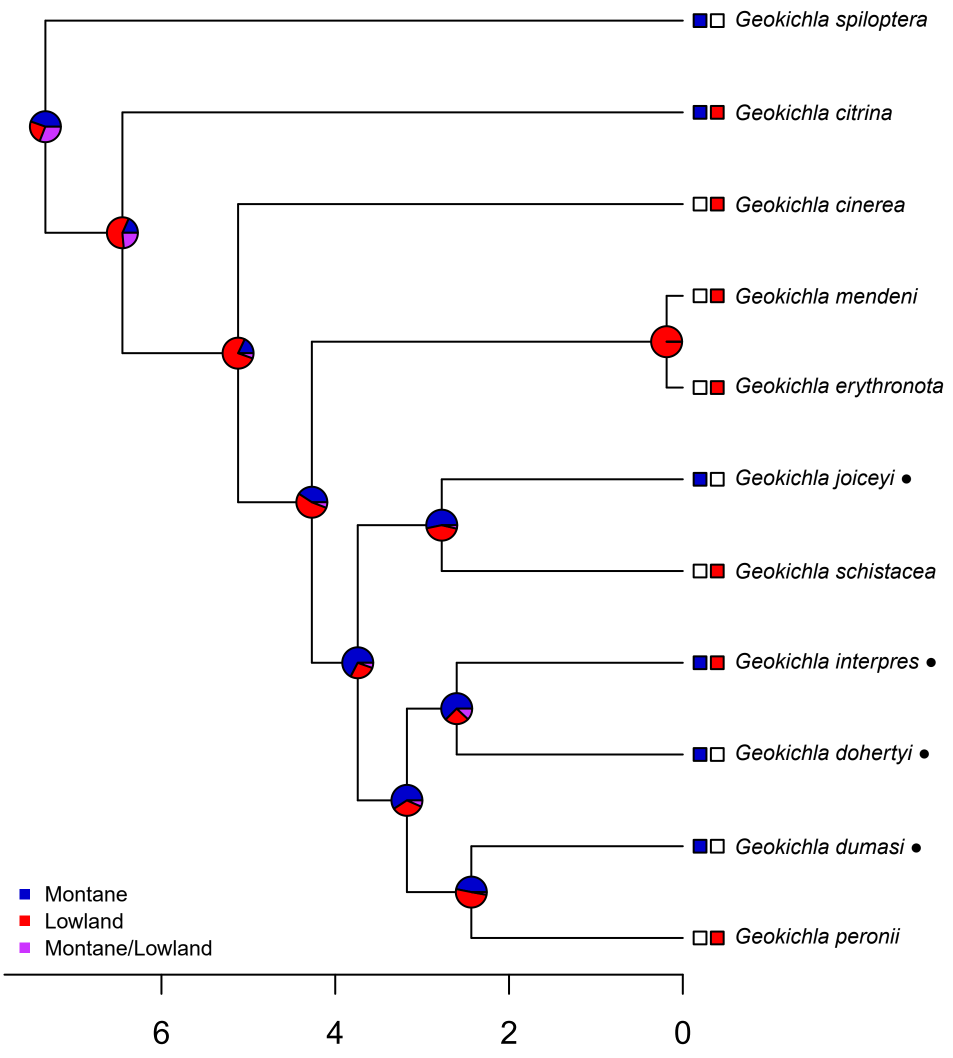

Geokichla spiloptera
Geokichla citrina
Geokichla cinerea
Geokichla mendeni
Geokichla erythronota
Geokichla joiceyi ●
Geokichla schistacea
Geokichla interpres ●
Geokichla dohertyi ●
Geokichla dumasi ●
Geokichla peronii
Montane
Lowland
Montane/Lowland
6
4
2
0

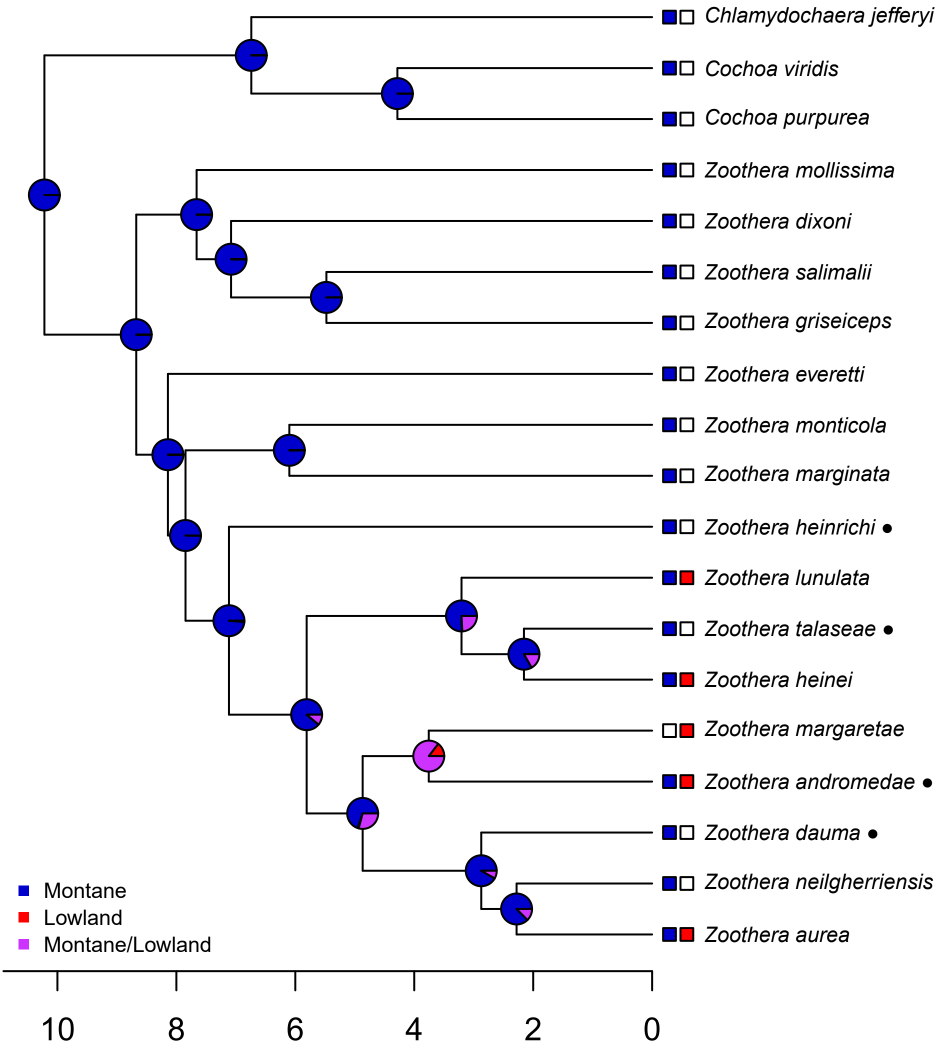

Chlamydochaera jefferyi
Cochoa viridis
Cochoa purpurea
Zoothera mollissima
Zoothera dixoni
Zoothera salimalii
Zoothera griseiceps
Zoothera everetti
Zoothera monticola
Zoothera marginata
Zoothera heinrichi ●
Zoothera lunulata
Zoothera talaseae ●
Zoothera heinei
Zoothera margaretae
Zoothera andromedae ●
Zoothera dauma ●
Zoothera neilgherriensis
Zoothera aurea
Montane
Lowland
Montane/Lowland
10
8
6
4
2
0

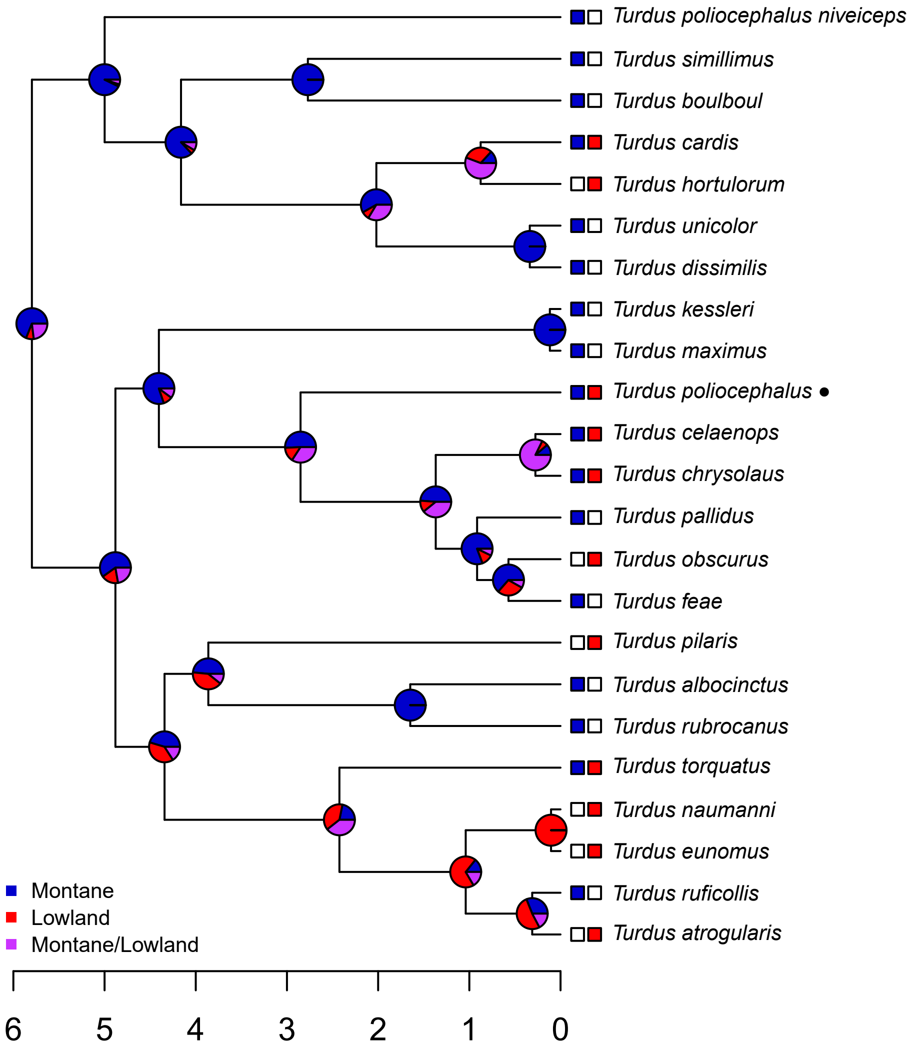

Turdus poliocephalus niveiceps
Turdus simillimus
Turdus boulboul
Turdus cardis
Turdus hortulorum
Turdus unicolor
Turdus dissimilis
Turdus kessleri
Turdus maximus
Turdus poliocephalus ●
Turdus celaenops
Turdus chrysolaus
Turdus pallidus
Turdus obscurus
Turdus feae
Turdus pilaris
Turdus albocinctus
Turdus rubrocanus
Turdus torquatus
Turdus naumanni
Turdus eunomus
Turdus ruficollis
Turdus atrogularis
Montane
Lowland
Montane/Lowland
6
5
4
3
2
1
0

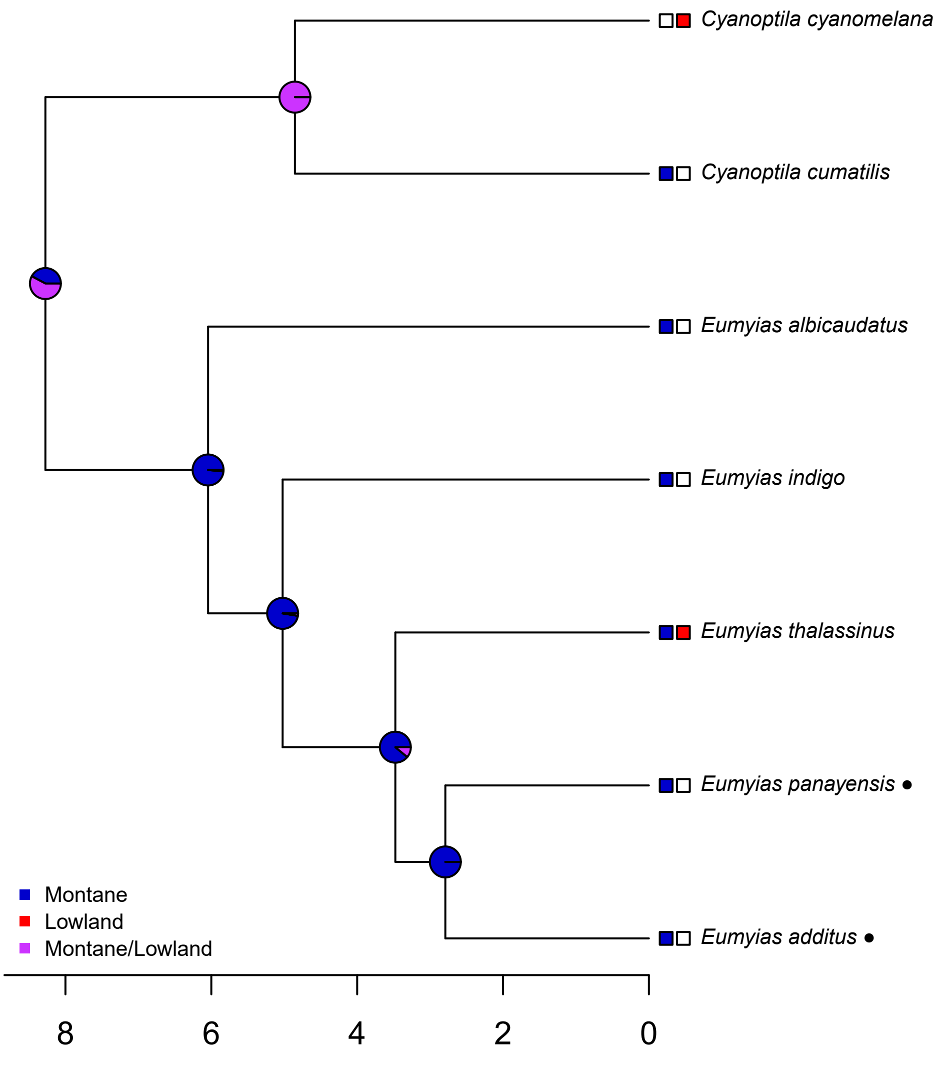
Cyanoptila cyanomelana
Cyanoptila cumatilis
Eumyias albicaudatus
Eumyias indigo
Eumyias thalassinus
Eumyias panayensis ●
Eumyias additus ●
Montane
Lowland
Montane/Lowland
8
6
4
2
0

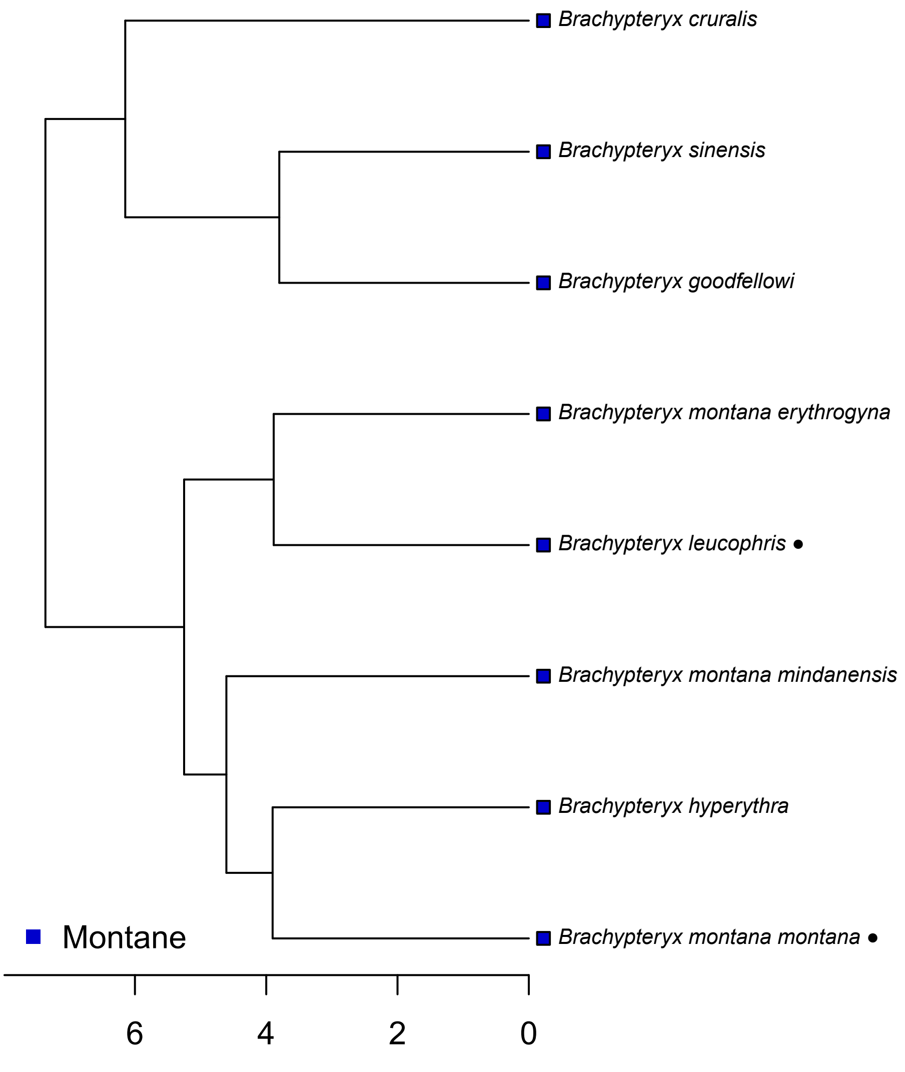

Brachypteryx cruralis
Brachypteryx sinensis
Brachypteryx goodfellowi
Brachypteryx montana erythrogyna
Brachypteryx leucophris ●
Brachypteryx montana mindanensis
Brachypteryx hyperythra
Brachypteryx montana montana ●
Montane
6
4
2
0

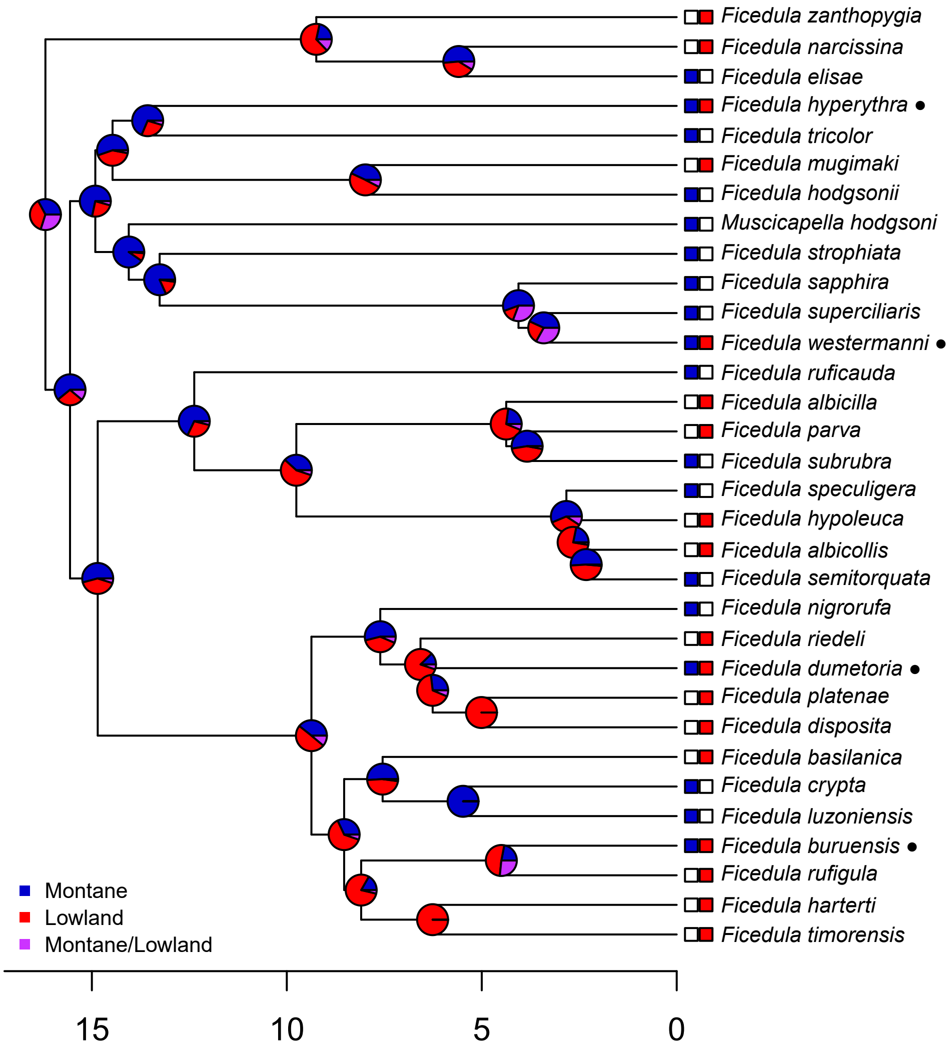

Ficedula zanthopygia
Ficedula narcissina
Ficedula elisae
Ficedula hyperythra ●
Ficedula tricolor
Ficedula mugimaki
Ficedula hodgsonii
Muscicapella hodgsoni
Ficedula strophiata
Ficedula sapphira
Ficedula superciliaris
Ficedula westermanni ●
Ficedula ruficauda
Ficedula albicilla
Ficedula parva
Ficedula subrubra
Ficedula speculigera
Ficedula hypoleuca
Ficedula albicollis
Ficedula semitorquata
Ficedula nigrorufa
Ficedula riedeli
Ficedula dumetoria ●
Ficedula platenae
Ficedula disposita
Ficedula basilanica
Ficedula crypta
Ficedula luzoniensis
Ficedula buruensis ●
Ficedula rufigula
Ficedula harterti
Ficedula timorensis
Montane
Lowland
Montane/Lowland
15
10
5
0

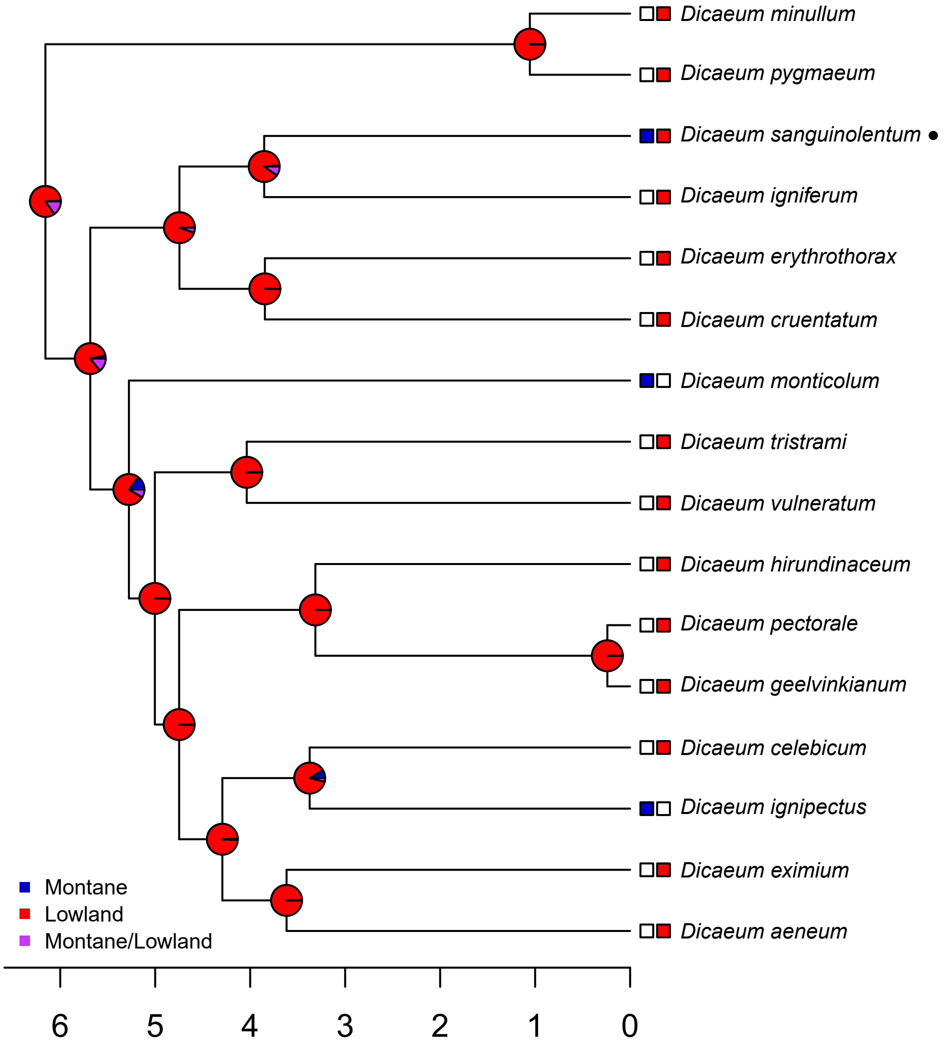

Dicaeum minullum
Dicaeum pygmaeum
Dicaeum sanguinolentum ●
Dicaeum igniferum
Dicaeum erythrothorax
Dicaeum cruentatum
Dicaeum monticolum
Dicaeum tristrami
Dicaeum vulneratum
Dicaeum hirundinaceum
Dicaeum pectorale
Dicaeum geelvinkianum
Dicaeum celebicum
Dicaeum ignipectus
Dicaeum eximium
Dicaeum aeneum
Montane
Lowland
Montane/Lowland
6
5
4
3
2
1
0

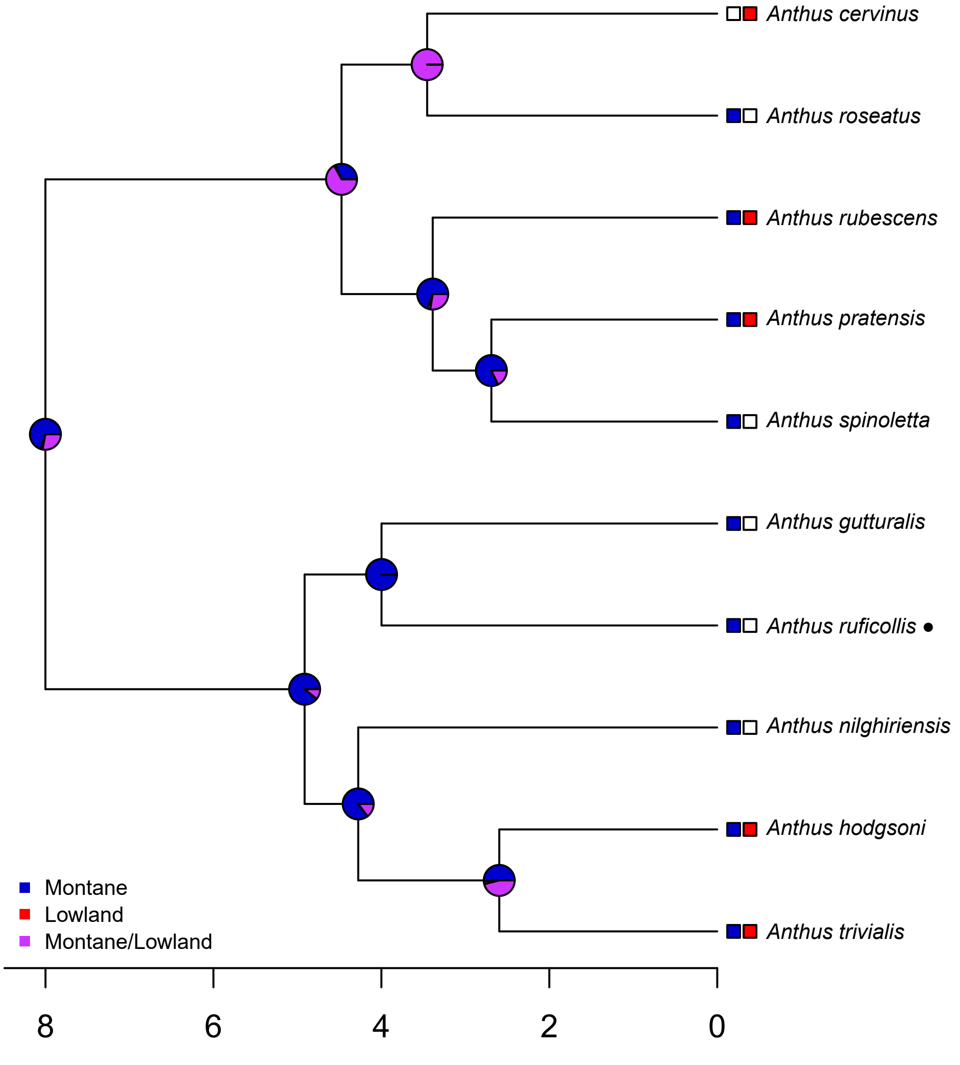
Anthus cervinus
Anthus roseatus
Anthus rubescens
Anthus pratensis
Anthus spinoletta
Anthus gutturalis
Anthus ruficollis •
Anthus nilghiriensis
Anthus hodgsoni
Anthus trivialis
Montane
Lowland
Montane/Lowland
8
6
4
2
0

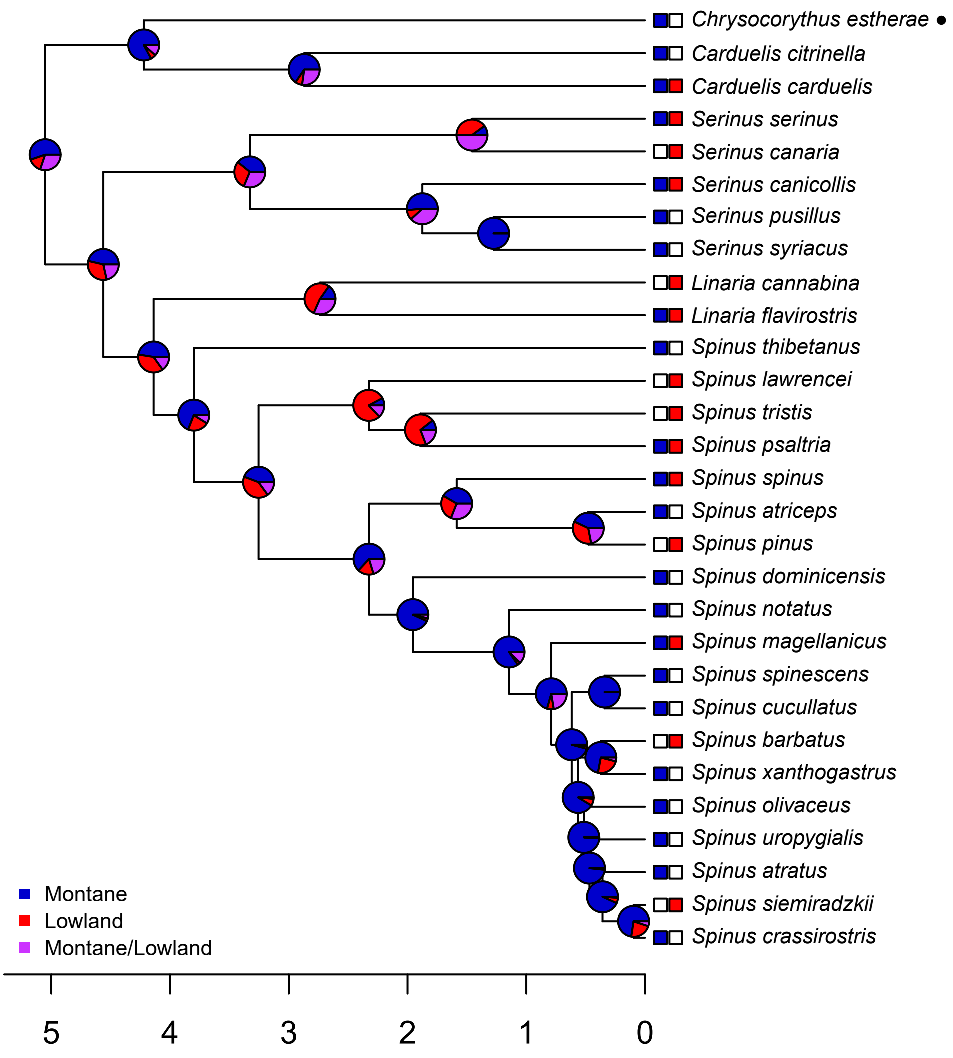

Chrysocorythus estherae ●
Carduelis citrinella
Carduelis carduelis
Serinus serinus
Serinus canaria
Serinus canicollis
Serinus pusillus
Serinus syriacus
Linaria cannabina
Linaria flavirostris
Spinus thibetanus
Spinus lawrencei
Spinus tristis
Spinus psaltria
Spinus spinus
Spinus atriceps
Spinus pinus
Spinus dominicensis
Spinus notatus
Spinus magellanicus
Spinus spinescens
Spinus cucullatus
Spinus barbatus
Spinus xanthogastrus
Spinus olivaceus
Spinus uropygialis
Spinus atratus
Spinus siemiradzkii
Spinus crassirostris
Montane
Lowland
Montane/Lowland
5
4
3
2
1
0

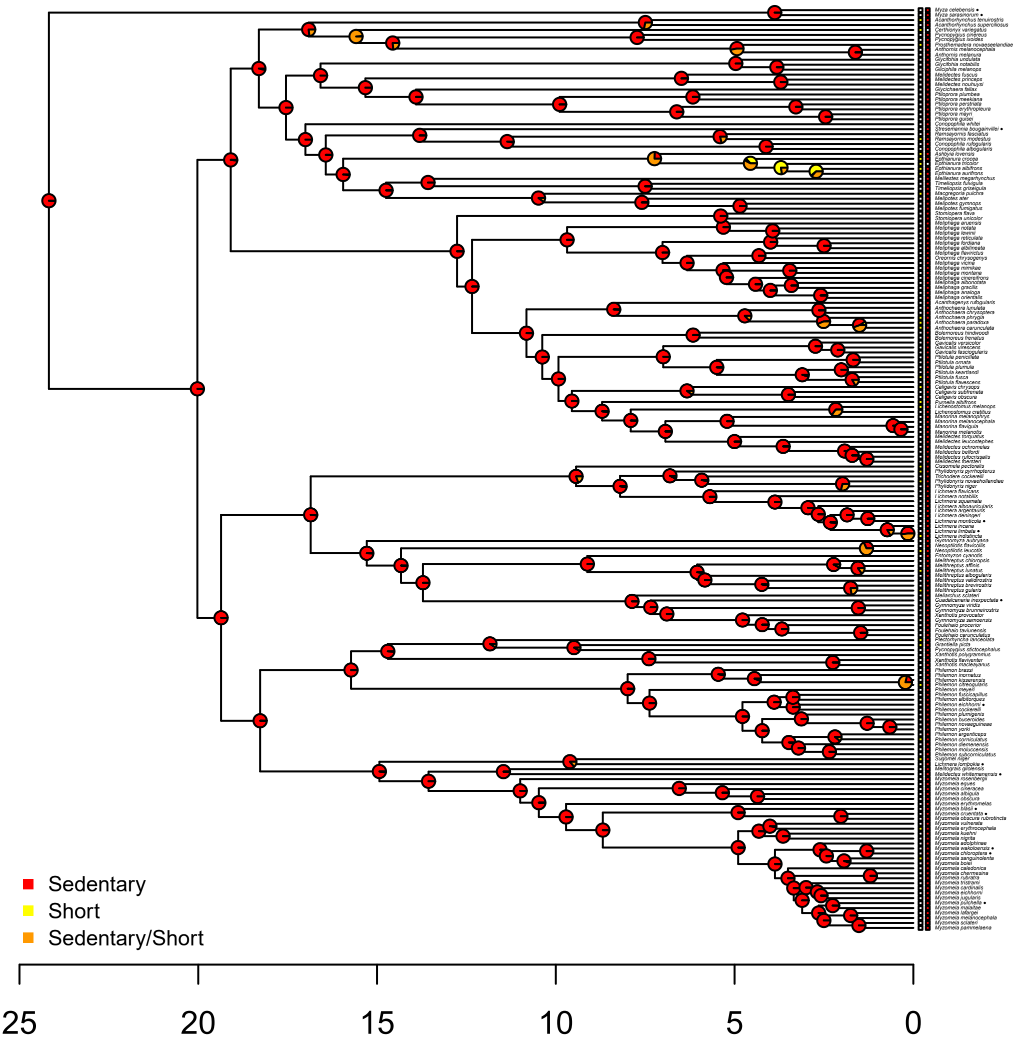
Sedentary
Short
Sedentary/Short
25
20
15
10
5
0

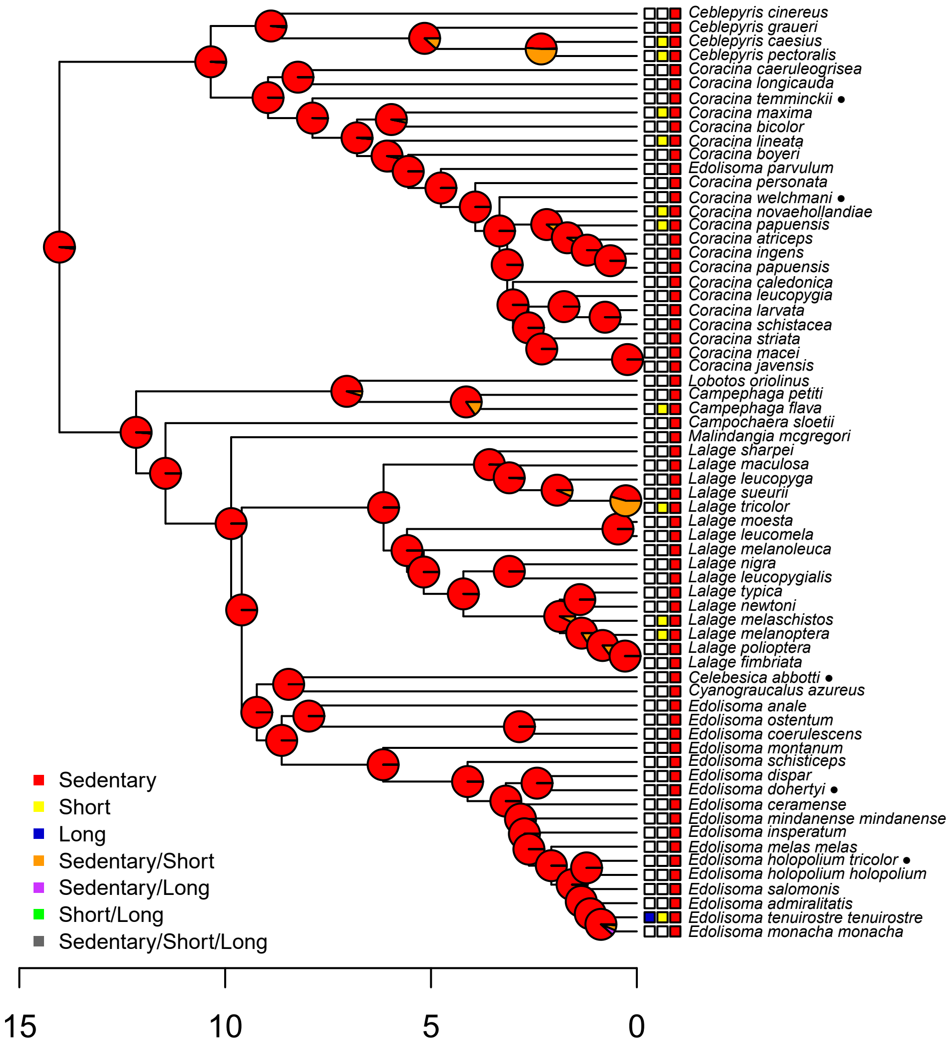

Ceblepyris cinereus
Ceblepyris graueri
Ceblepyris caesius
Ceblepyris pectoralis
Coracina caeruleogrisea
Coracina longicauda
Coracina temminckii ●
Coracina maxima
Coracina bicolor
Coracina lineata
Coracina boyeri
Edolisoma parvulum
Coracina personata
Coracina welchmani ●
Coracina novaehollandiae
Coracina papuensis
Coracina atriceps
Coracina ingens
Coracina papuensis
Coracina caledonica
Coracina leucopygia
Coracina larvata
Coracina schistacea
Coracina striata
Coracina macei
Coracina javensis
Lobotos oriolinus
Campephaga petiti
Campephaga flava
Campochaera sloetii
Malindangia mcgregori
Lalage sharpei
Lalage maculosa
Lalage leucopyga
Lalage sueurii
Lalage tricolor
Lalage moesta
Lalage leucomela
Lalage melanoleuca
Lalage nigra
Lalage leucopygialis
Lalage typica
Lalage newtoni
Lalage melaschistos
Lalage melanoptera
Lalage polioptera
Lalage fimbriata
Celebesica abbotti ●
Cyanograucalus azureus
Edolisoma anale
Edolisoma ostentum
Edolisoma coerulescens
Edolisoma montanum
Edolisoma schisticeps
Edolisoma dispar
Edolisoma dohertyi ●
Edolisoma ceramense
Edolisoma mindanense mindanense
Edolisoma insperatum
Edolisoma melas melas
Edolisoma holopolium tricolor ●
Edolisoma holopolium holopolium
Edolisoma salomonis
Edolisoma admiralitatis
Edolisoma tenuirostre tenuirostre
Edolisoma monacha monacha
Sedentary
Short
Long
Sedentary/Short
Sedentary/Long
Short/Long
Sedentary/Short/Long
15
10
5
0

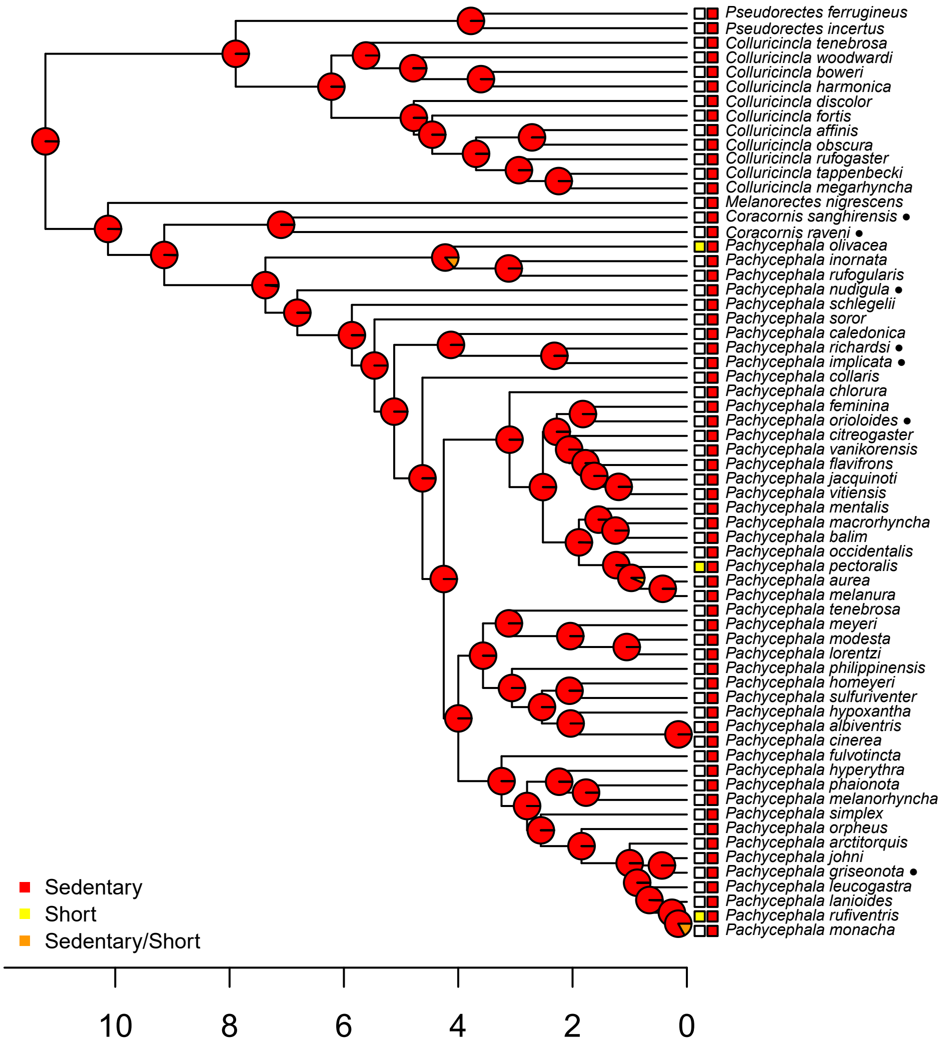

Pseudorectes ferrugineus
Pseudorectes incertus
Colluricincla tenebrosa
Colluricincla woodwardi
Colluricincla boweri
Colluricincla harmonica
Colluricincla discolor
Colluricincla fortis
Colluricincla affinis
Colluricincla obscura
Colluricincla rufogaster
Colluricincla tappenbecki
Colluricincla megarhyncha
Melanorectes nigrescens
Coracornis sanghirensis ●
Coracornis raveni ●
Pachycephala olivacea
Pachycephala inornata
Pachycephala rufogularis
Pachycephala nudigula ●
Pachycephala schlegelii
Pachycephala soror
Pachycephala caledonica
Pachycephala richardsi ●
Pachycephala implicata ●
Pachycephala collaris
Pachycephala chlorura
Pachycephala feminina
Pachycephala orioloides ●
Pachycephala citreogaster
Pachycephala vanikorensis
Pachycephala flavifrons
Pachycephala jacquinoti
Pachycephala vitiensis
Pachycephala mentalis
Pachycephala macrorhyncha
Pachycephala balim
Pachycephala occidentalis
Pachycephala pectoralis
Pachycephala aurea
Pachycephala melanura
Pachycephala tenebrosa
Pachycephala meyeri
Pachycephala modesta
Pachycephala lorentzi
Pachycephala philippinensis
Pachycephala homeyeri
Pachycephala sulfuriventer
Pachycephala hypoxantha
Pachycephala albiventris
Pachycephala cinerea
Pachycephala fulvotincta
Pachycephala hyperythra
Pachycephala phaionota
Pachycephala melanorhyncha
Pachycephala simplex
Pachycephala orpheus
Pachycephala arctitorquis
Pachycephala johni
Pachycephala griseonota ●
Pachycephala leucogastra
Pachycephala lanioides
Pachycephala rufiventris
Pachycephala monacha
Sedentary
Short
Sedentary/Short
10
8
6
4
2
0

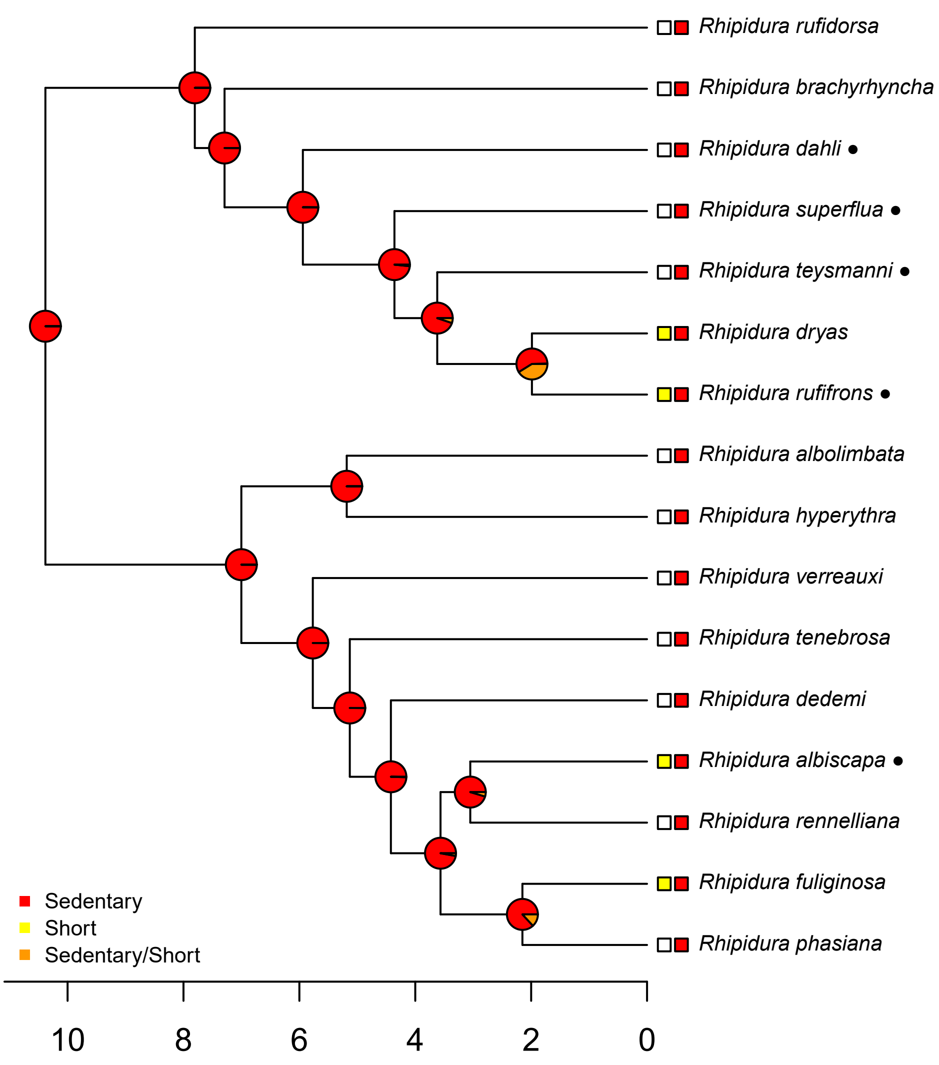

Rhipidura rufidorsa
Rhipidura brachyrhyncha
Rhipidura dahli ●
Rhipidura superflua ●
Rhipidura teysmanni ●
Rhipidura dryas
Rhipidura rufifrons ●
Rhipidura albolimbata
Rhipidura hyperythra
Rhipidura verreauxi
Rhipidura tenebrosa
Rhipidura dedemi
Rhipidura albiscapa ●
Rhipidura rennelliana
Rhipidura fuliginosa
Rhipidura phasiana
Sedentary
Short
Sedentary/Short
10
8
6
4
2
0

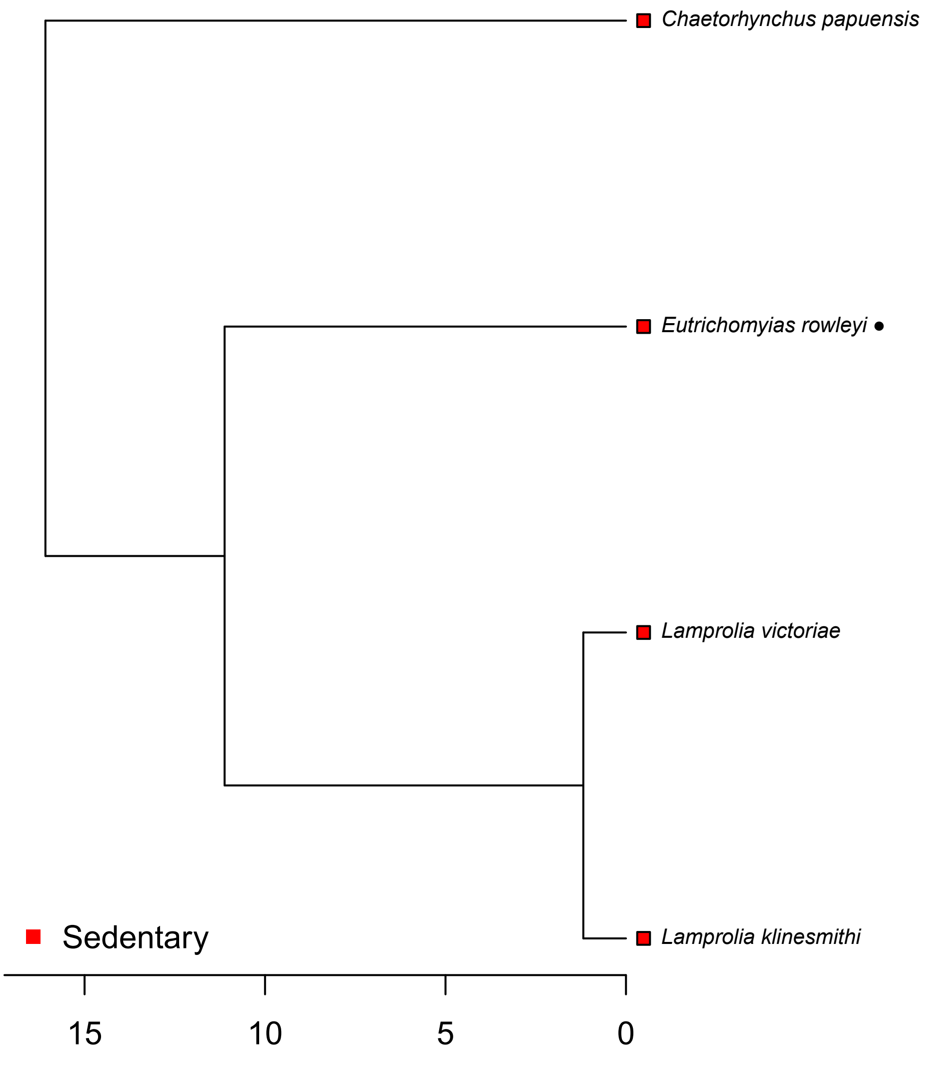

Chaetorhynchus papuensis
Eutrichomyias rowleyi ●
Lamprolia victoriae
Lamprolia klinesmithi
Sedentary
15
10
5
0

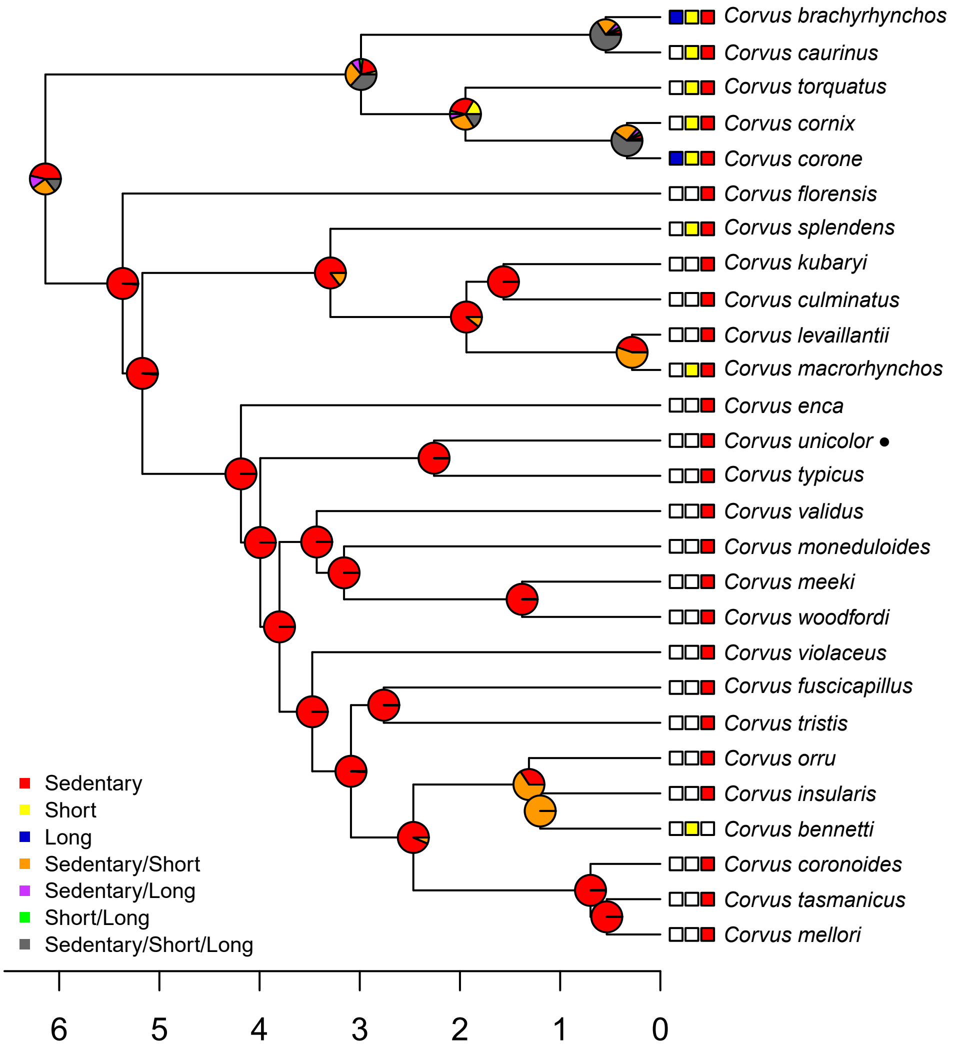

Corvus brachyrhynchos
Corvus caurinus
Corvus torquatus
Corvus cornix
Corvus corone
Corvus florensis
Corvus splendens
Corvus kubaryi
Corvus culminatus
Corvus levaillantii
Corvus macrorhynchos
Corvus enca
Corvus unicolor ●
Corvus typicus
Corvus validus
Corvus moneduloides
Corvus meeki
Corvus woodfordi
Corvus violaceus
Corvus fuscicapillus
Corvus tristis
Corvus orru
Corvus insularis
Corvus bennetti
Corvus coronoides
Corvus tasmanicus
Corvus mellori
Sedentary
Short
Long
Sedentary/Short
Sedentary/Long
Short/Long
Sedentary/Short/Long
6
5
4
3
2
1
0

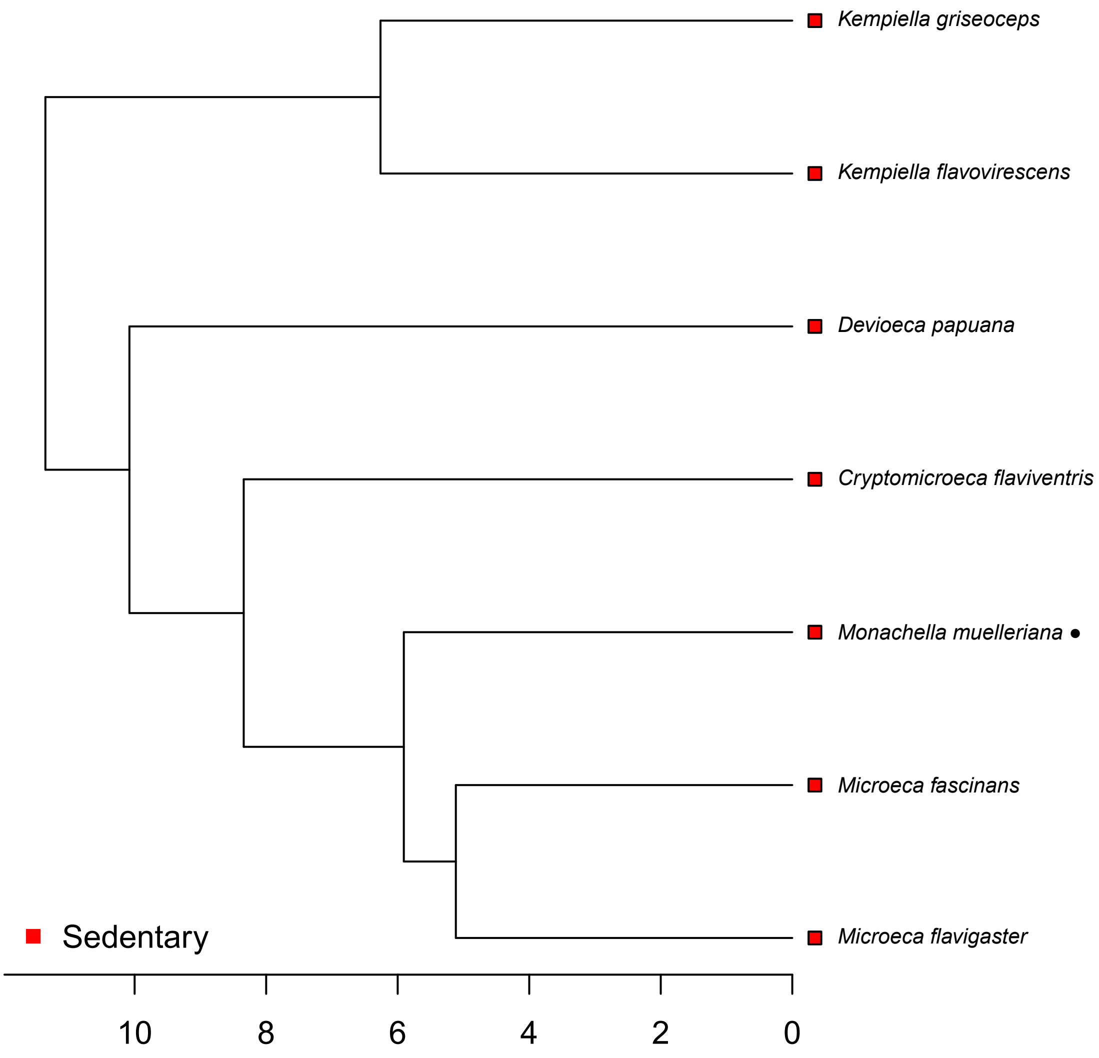
Kempiella griseoceps
Kempiella flavovirescens
Devioeca papuana
Cryptomicroeca flaviventris
Monachella muelleriana ●
Microeca fascinans
Microeca flavigaster
Sedentary
10
8
6
4
2
0

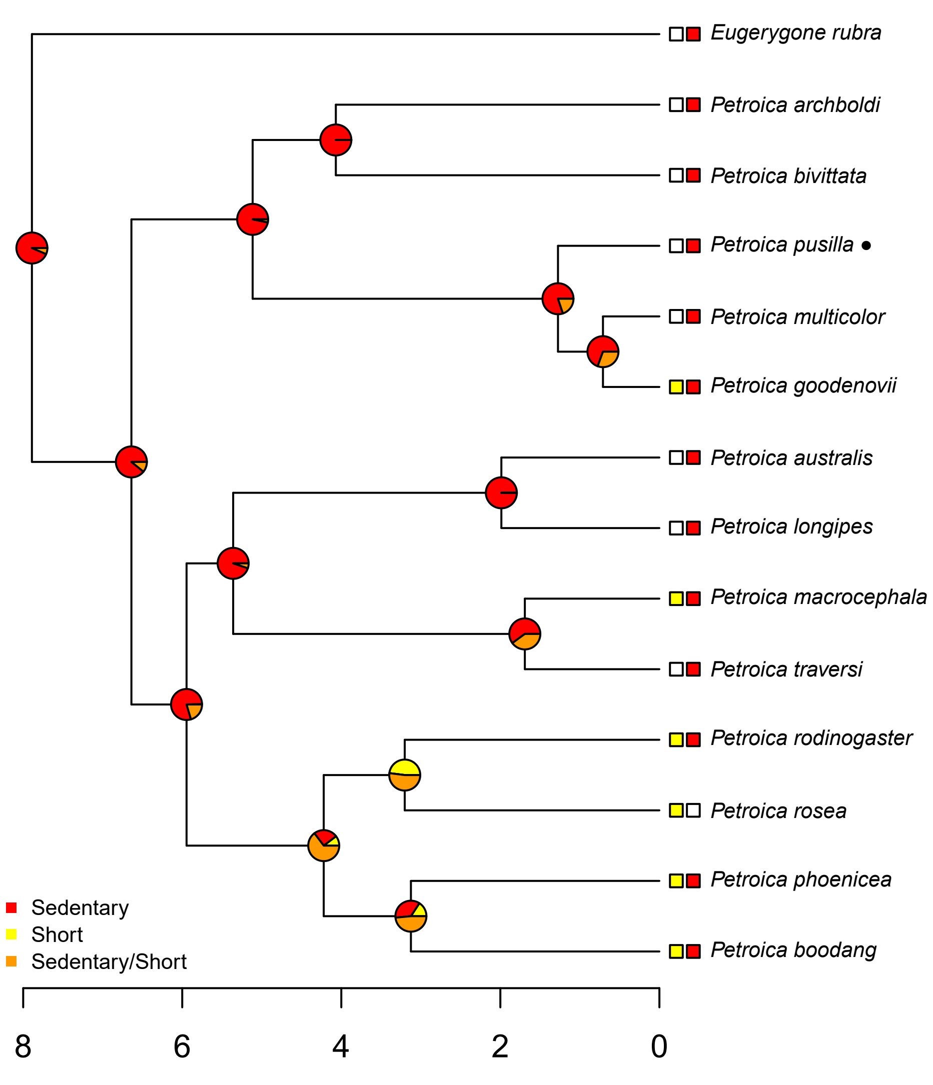

Eugerygone rubra
Petroica archboldi
Petroica bivittata
Petroica pusilla ●
Petroica multicolor
Petroica goodenovii
Petroica australis
Petroica longipes
Petroica macrocephala
Petroica traversi
Petroica rodinogaster
Petroica rosea
Petroica phoenicea
Petroica boodang
Sedentary
Short
Sedentary/Short
8
6
4
2
0

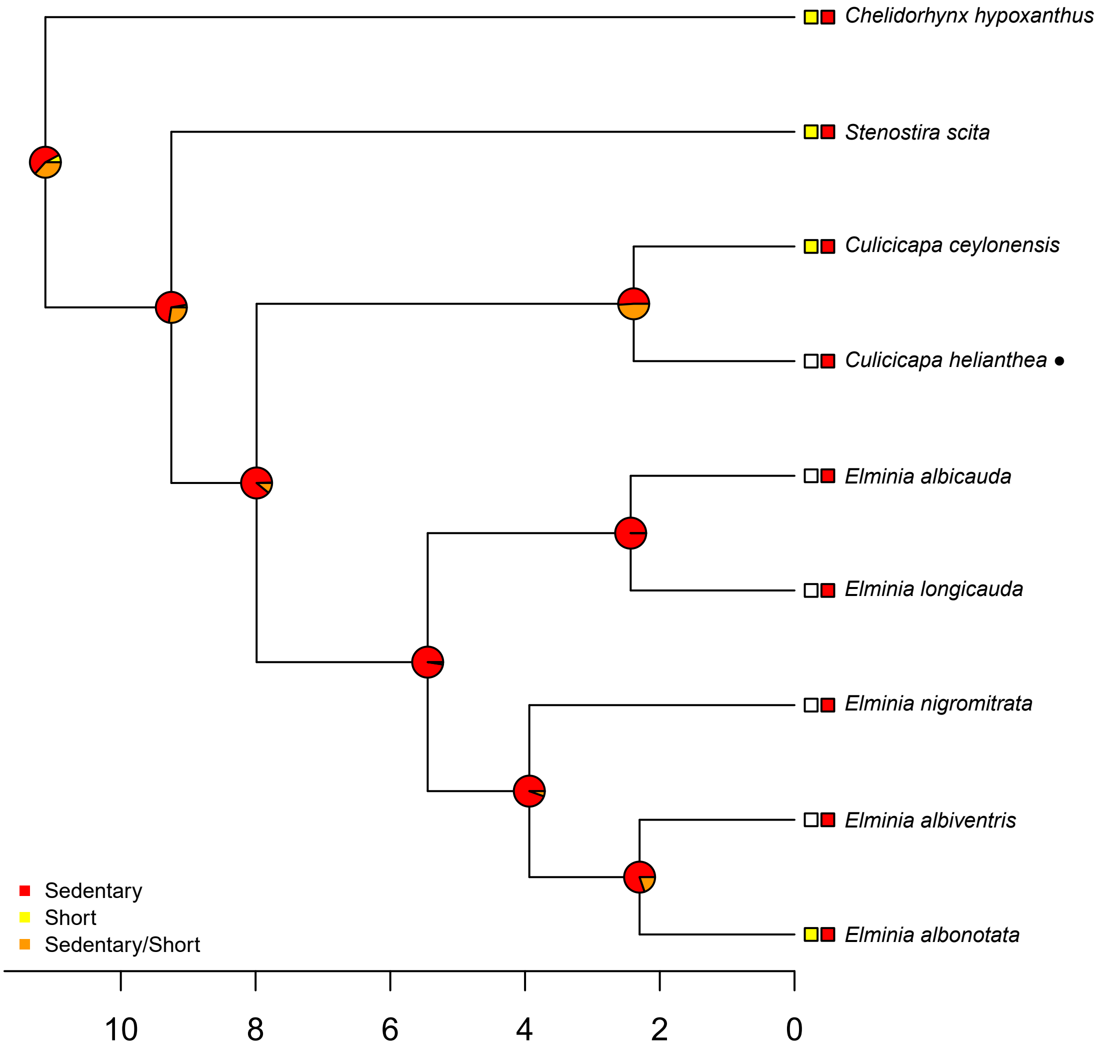

Chelidorhynx hypoxanthus
Stenostira scita
Culicicapa ceylonensis
Culicicapa helianthea ●
Elminia albicauda
Elminia longicauda
Elminia nigromitrata
Elminia albiventris
Elminia albonotata
Sedentary
Short
Sedentary/Short
10
8
6
4
2
0

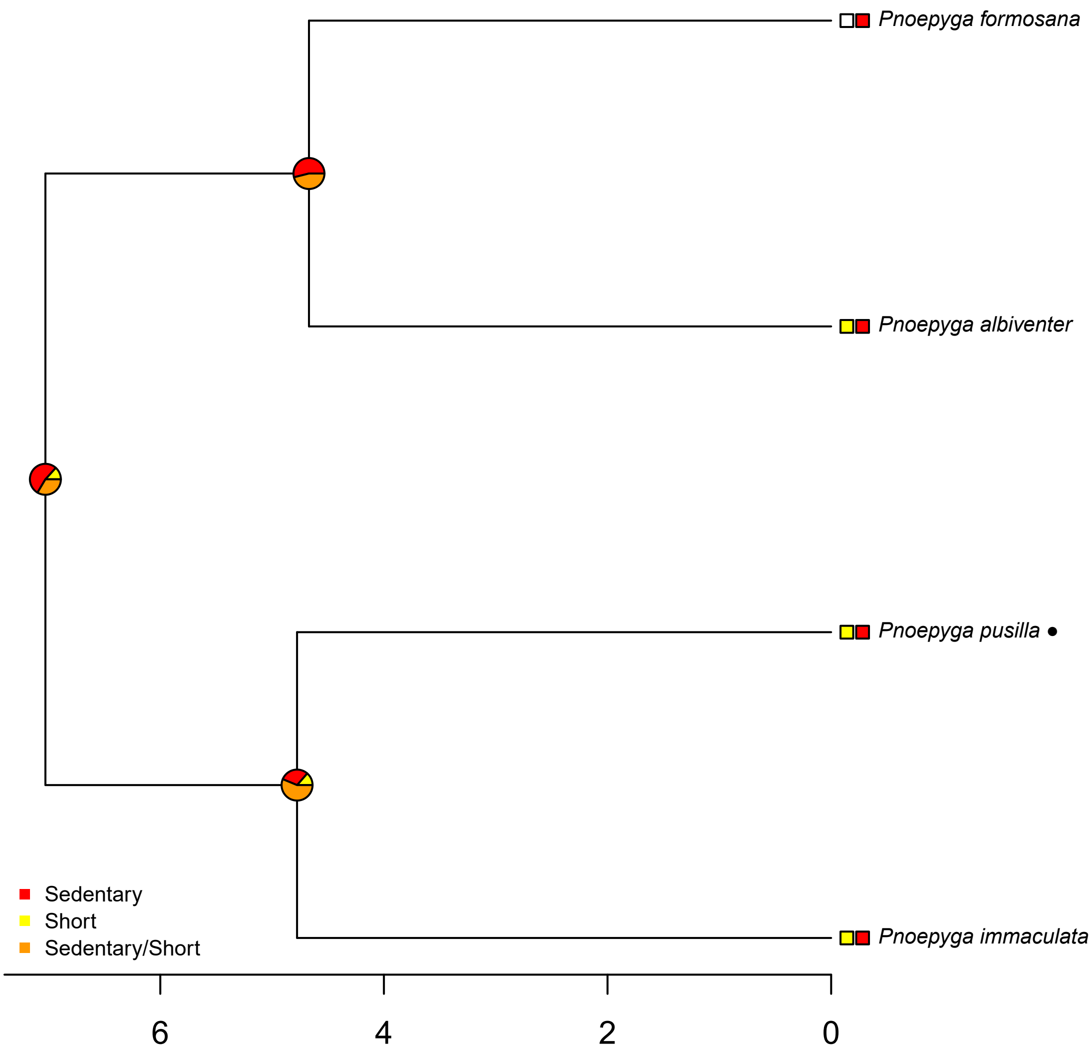

Pnoepyga formosana
Pnoepyga albiventer
Pnoepyga pusilla
Pnoepyga immaculata
Sedentary
Short
Sedentary/Short
6
4
2
0

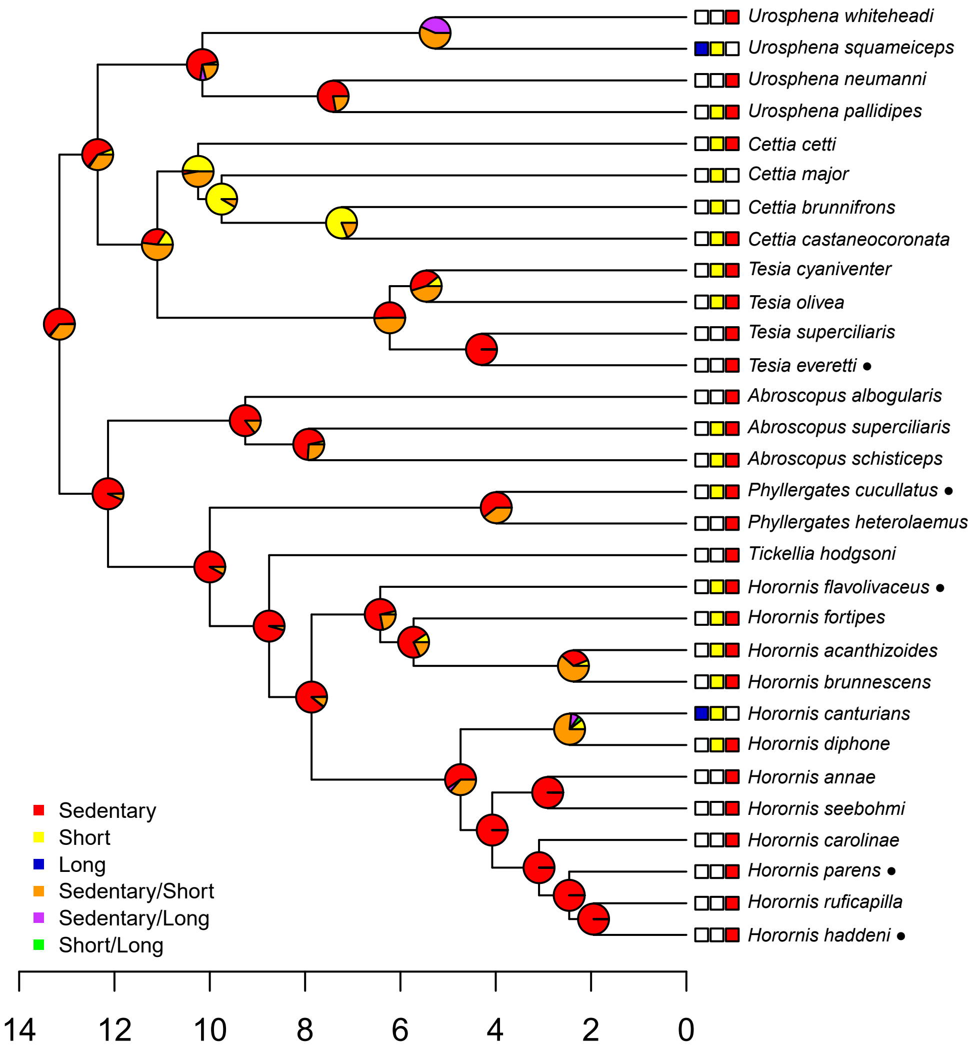

Urosphena whiteheadi
Urosphena squameiceps
Urosphena neumanni
Urosphena pallidipes
Cettia cetti
Cettia major
Cettia brunnifrons
Cettia castaneocoronata
Tesia cyaniventer
Tesia olivea
Tesia superciliaris
Tesia everetti ●
Abroscopus albogularis
Abroscopus superciliaris
Abroscopus schisticeps
Phyllergates cucullatus ●
Phyllergates heterolaemus
Tickellia hodgsoni
Horornis flavolivaceus ●
Horornis fortipes
Horornis acanthizoides
Horornis brunnescens
Horornis canturians
Horornis diphone
Horornis annae
Horornis seebohmi
Horornis carolinae
Horornis parens ●
Horornis ruficapilla
Horornis haddeni ●
Sedentary
Short
Long
Sedentary/Short
Sedentary/Long
Short/Long
14
12
10
8
6
4
2
0

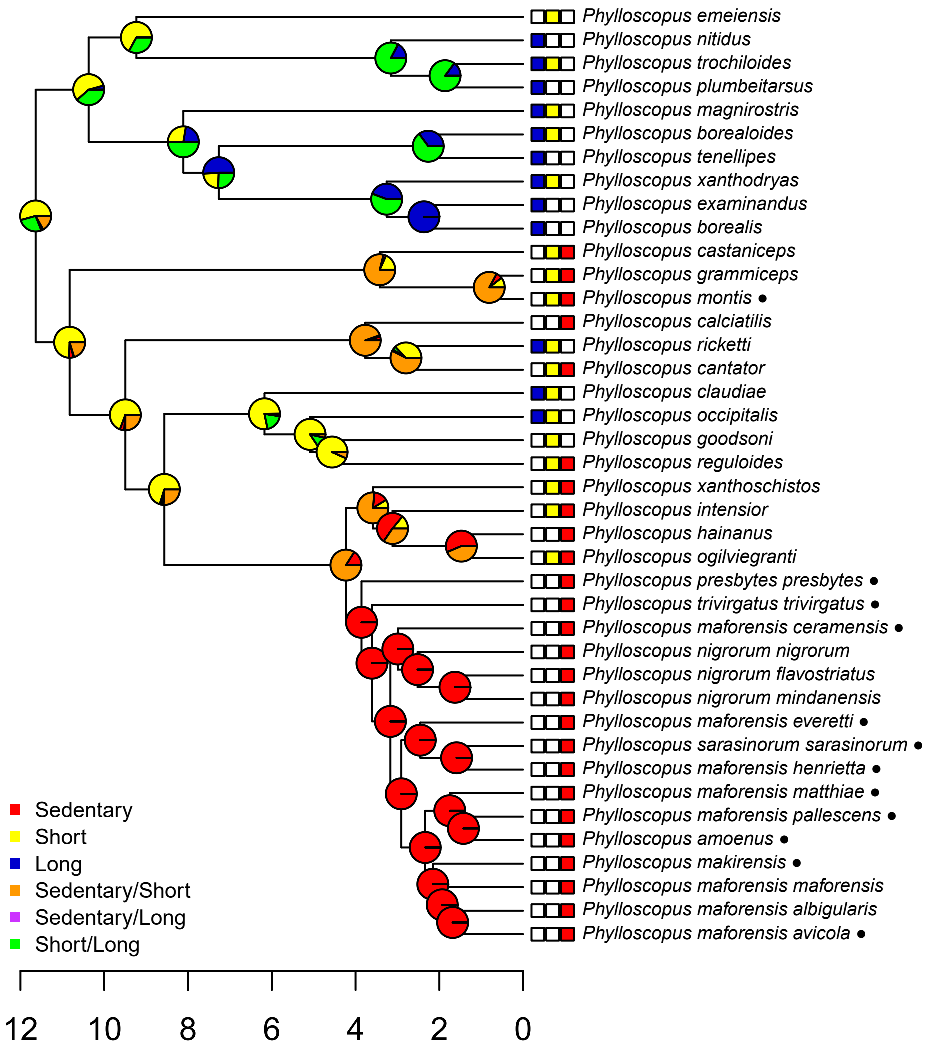

Phylloscopus emeiensis
Phylloscopus nitidus
Phylloscopus trochiloides
Phylloscopus plumbeitarsus
Phylloscopus magnirostris
Phylloscopus borealoides
Phylloscopus tenellipes
Phylloscopus xanthodryas
Phylloscopus examinandus
Phylloscopus borealis
Phylloscopus castaniceps
Phylloscopus grammiceps
Phylloscopus montis ●
Phylloscopus calciatilis
Phylloscopus ricketti
Phylloscopus cantator
Phylloscopus claudiae
Phylloscopus occipitalis
Phylloscopus goodsoni
Phylloscopus reguloides
Phylloscopus xanthoschistos
Phylloscopus intensior
Phylloscopus hainanus
Phylloscopus ogilviegranti
Phylloscopus presbytes presbytes ●
Phylloscopus trivirgatus trivirgatus ●
Phylloscopus maforensis ceramensis ●
Phylloscopus nigrorum nigrorum
Phylloscopus nigrorum flavostriatus
Phylloscopus nigrorum mindanensis
Phylloscopus maforensis everetti ●
Phylloscopus sarasinorum sarasinorum ●
Phylloscopus maforensis henrietta ●
Phylloscopus maforensis matthiae ●
Phylloscopus maforensis pallescens ●
Phylloscopus amoenus ●
Phylloscopus makirensis ●
Phylloscopus maforensis maforensis
Phylloscopus maforensis albigularis
Phylloscopus maforensis avicola ●
Sedentary
Short
Long
Sedentary/Short
Sedentary/Long
Short/Long
12
10
8
6
4
2
0

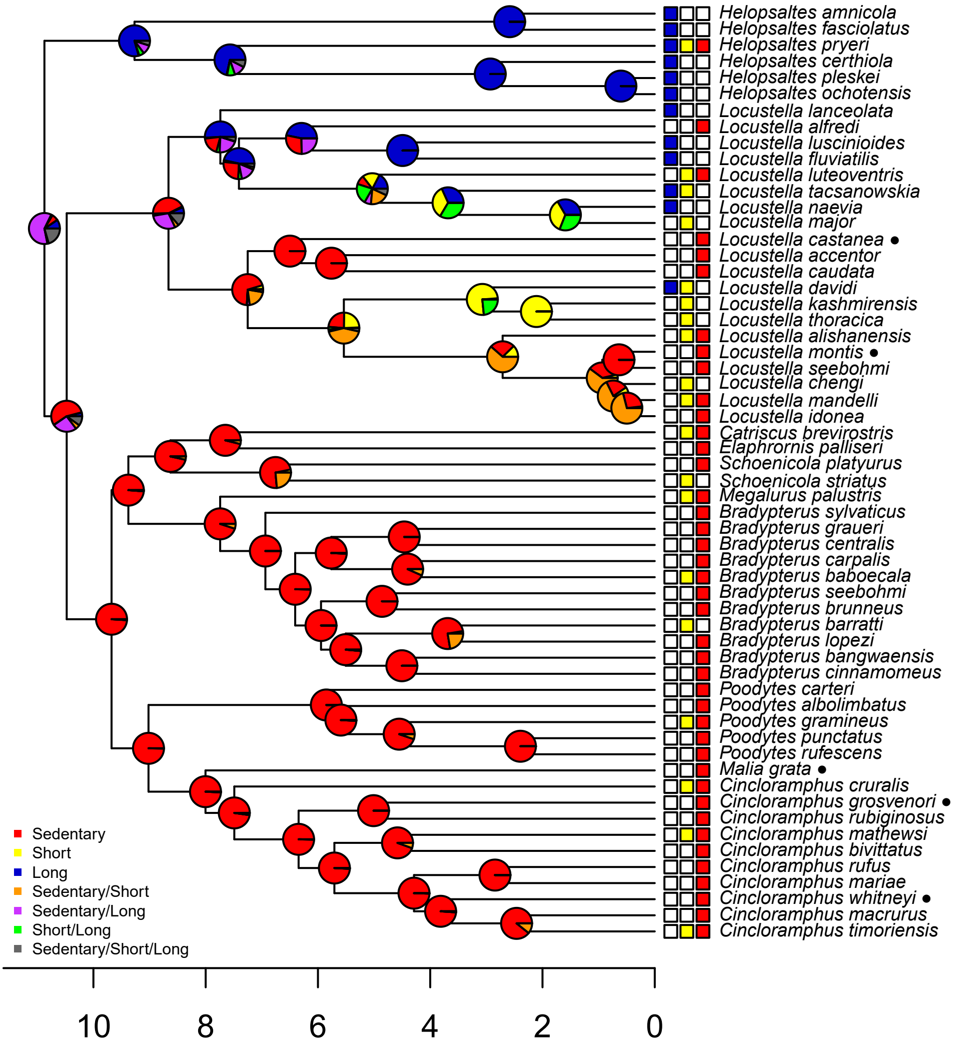

Helopsaltes amnicola
Helopsaltes fasciolatus
Helopsaltes pryeri
Helopsaltes certhiola
Helopsaltes pleskei
Helopsaltes ochotensis
Locustella lanceolata
Locustella alfredi
Locustella luscinioides
Locustella fluviatilis
Locustella luteoventris
Locustella tacsanowskia
Locustella naevia
Locustella major
Locustella castanea ●
Locustella accentor
Locustella caudata
Locustella davidi
Locustella kashmirensis
Locustella thoracica
Locustella alishanensis
Locustella montis ●
Locustella seebohmi
Locustella chengi
Locustella mandelli
Locustella idonea
Catriscus brevirostris
Elaphrornis palliseri
Schoenicola platyurus
Schoenicola striatus
Megalurus palustris
Bradypterus sylvaticus
Bradypterus graueri
Bradypterus centralis
Bradypterus carpalis
Bradypterus baboecala
Bradypterus seebohmi
Bradypterus brunneus
Bradypterus barratti
Bradypterus lopezi
Bradypterus bangwaensis
Bradypterus cinnamomeus
Poodytes carteri
Poodytes albolimbatus
Poodytes gramineus
Poodytes punctatus
Poodytes rufescens
Malia grata ●
Cincloramphus cruralis
Cincloramphus grosvenori ●
Cincloramphus rubiginosus
Cincloramphus mathewsi
Cincloramphus bivittatus
Cincloramphus rufus
Cincloramphus mariae
Cincloramphus whitneyi ●
Cincloramphus macrurus
Cincloramphus timoriensis
Sedentary
Short
Long
Sedentary/Short
Sedentary/Long
Short/Long
Sedentary/Short/Long
10
8
6
4
2
0

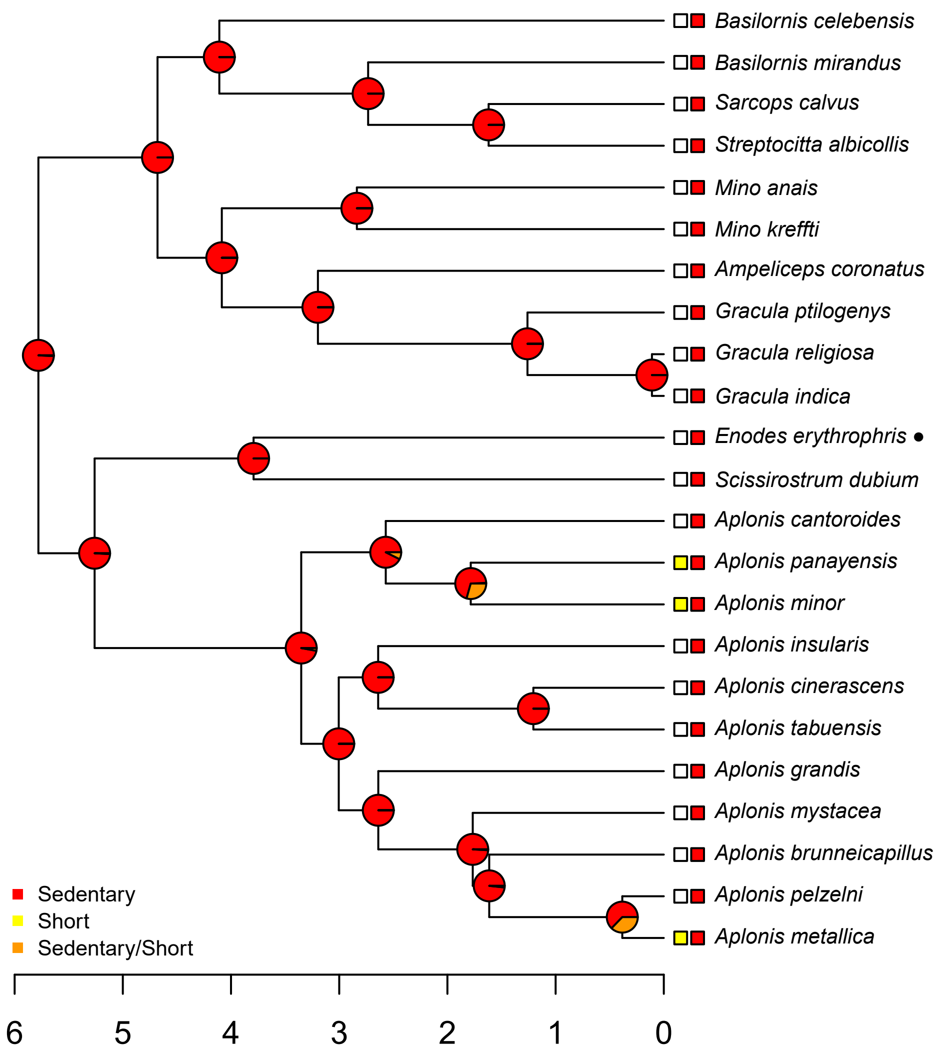

Basilornis celebensis
Basilornis mirandus
Sarcops calvus
Streptocitta albicollis
Mino anais
Mino kreffti
Ampeliceps coronatus
Gracula ptilogenys
Gracula religiosa
Gracula indica
Enodes erythrophris
Scissirostrum dubium
Aplonis cantoroides
Aplonis panayensis
Aplonis minor
Aplonis insularis
Aplonis cinerascens
Aplonis tabuensis
Aplonis grandis
Aplonis mystacea
Aplonis brunneicapillus
Aplonis pelzelni
Aplonis metallica
Sedentary
Short
Sedentary/Short
6
5
4
3
2
1
0

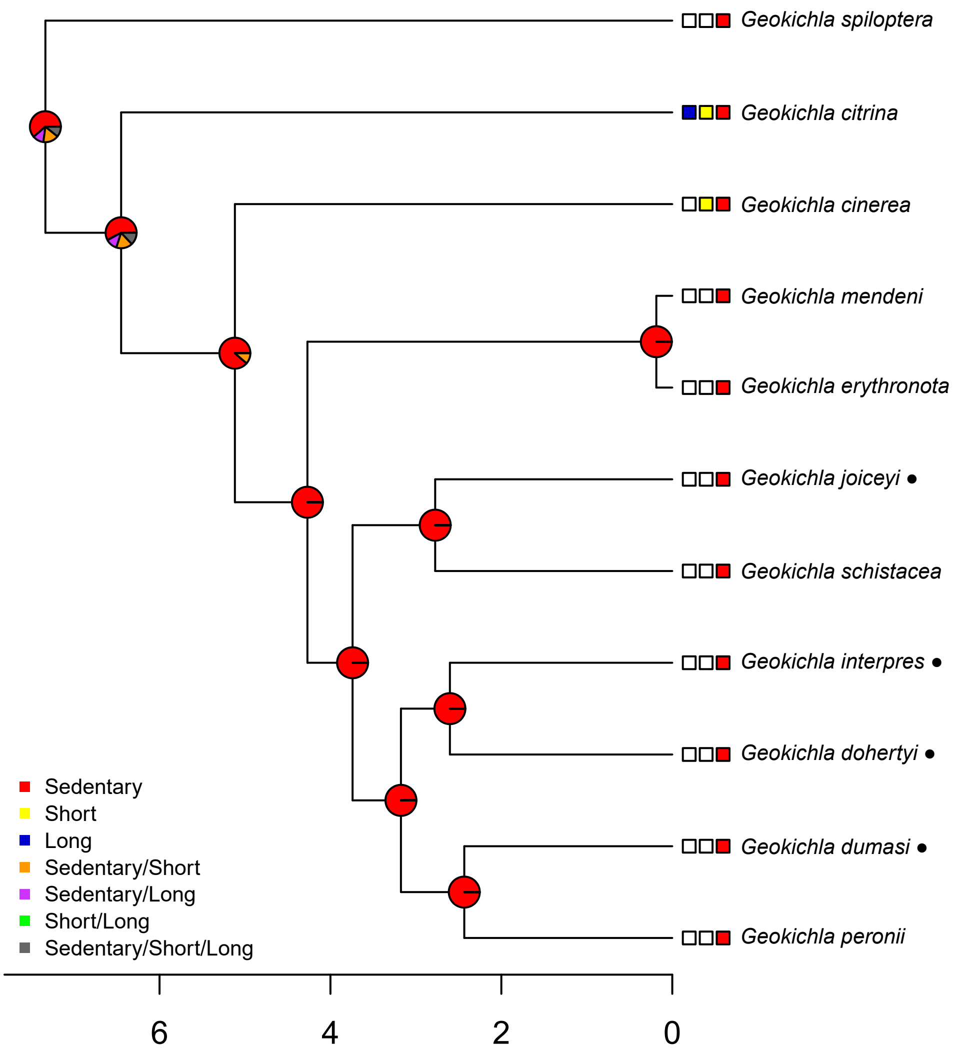

Geokichla spiloptera
Geokichla citrina
Geokichla cinerea
Geokichla mendeni
Geokichla erythronota
Geokichla joiceyi ●
Geokichla schistacea
Geokichla interpres ●
Geokichla dohertyi ●
Geokichla dumasi ●
Geokichla peronii
Sedentary
Short
Long
Sedentary/Short
Sedentary/Long
Short/Long
Sedentary/Short/Long
6
4
2
0

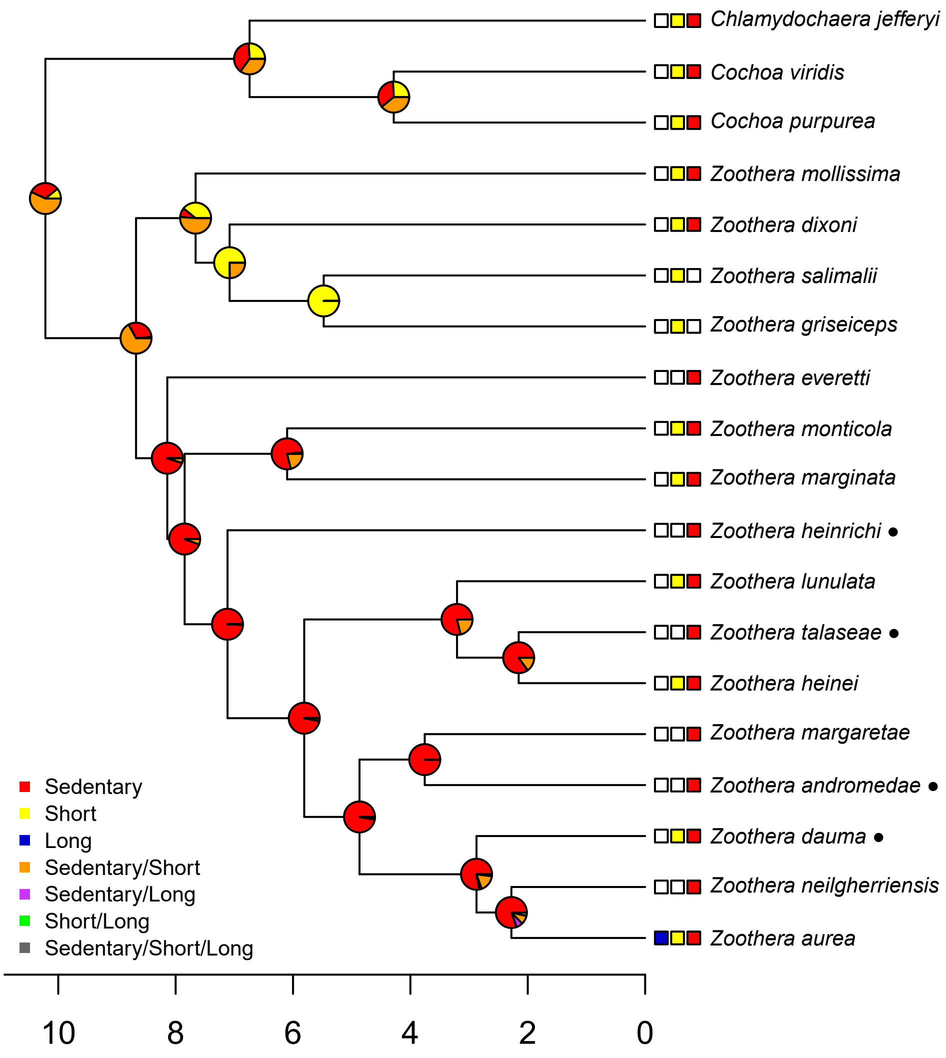

Chlamydochaera jefferyi
Cochoa viridis
Cochoa purpurea
Zoothera mollissima
Zoothera dixoni
Zoothera salimalii
Zoothera griseiceps
Zoothera everetti
Zoothera monticola
Zoothera marginata
Zoothera heinrichi ●
Zoothera lunulata
Zoothera talaseae ●
Zoothera heinei
Zoothera margaretae
Zoothera andromedae ●
Zoothera dauma ●
Zoothera neilgherriensis
Zoothera aurea
Sedentary
Short
Long
Sedentary/Short
Sedentary/Long
Short/Long
Sedentary/Short/Long
10
8
6
4
2
0

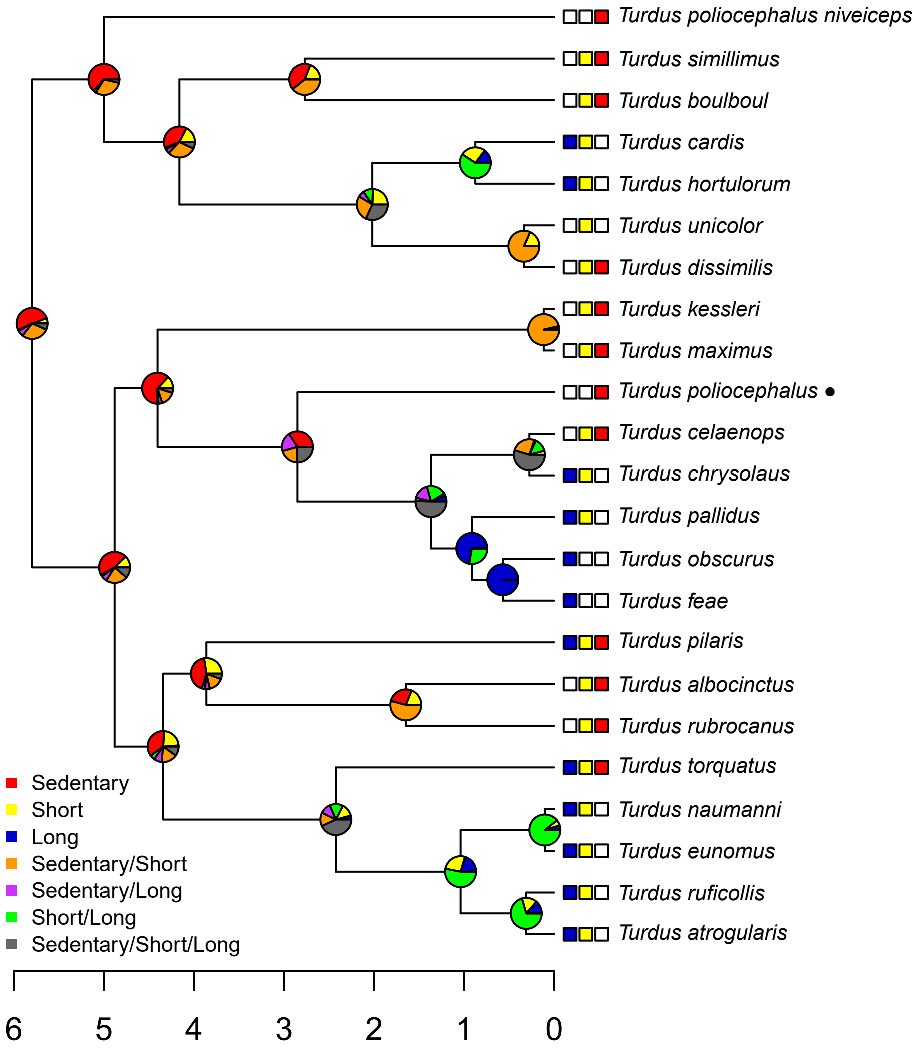

Turdus poliocephalus niveiceps
Turdus simillimus
Turdus boulboul
Turdus cardis
Turdus hortulorum
Turdus unicolor
Turdus dissimilis
Turdus kessleri
Turdus maximus
Turdus poliocephalus ●
Turdus celaenops
Turdus chrysolaus
Turdus pallidus
Turdus obscurus
Turdus feae
Turdus pilaris
Turdus albocinctus
Turdus rubrocanus
Turdus torquatus
Turdus naumanni
Turdus eunomus
Turdus ruficollis
Turdus atrogularis
Sedentary
Short
Long
Sedentary/Short
Sedentary/Long
Short/Long
Sedentary/Short/Long
6
5
4
3
2
1
0

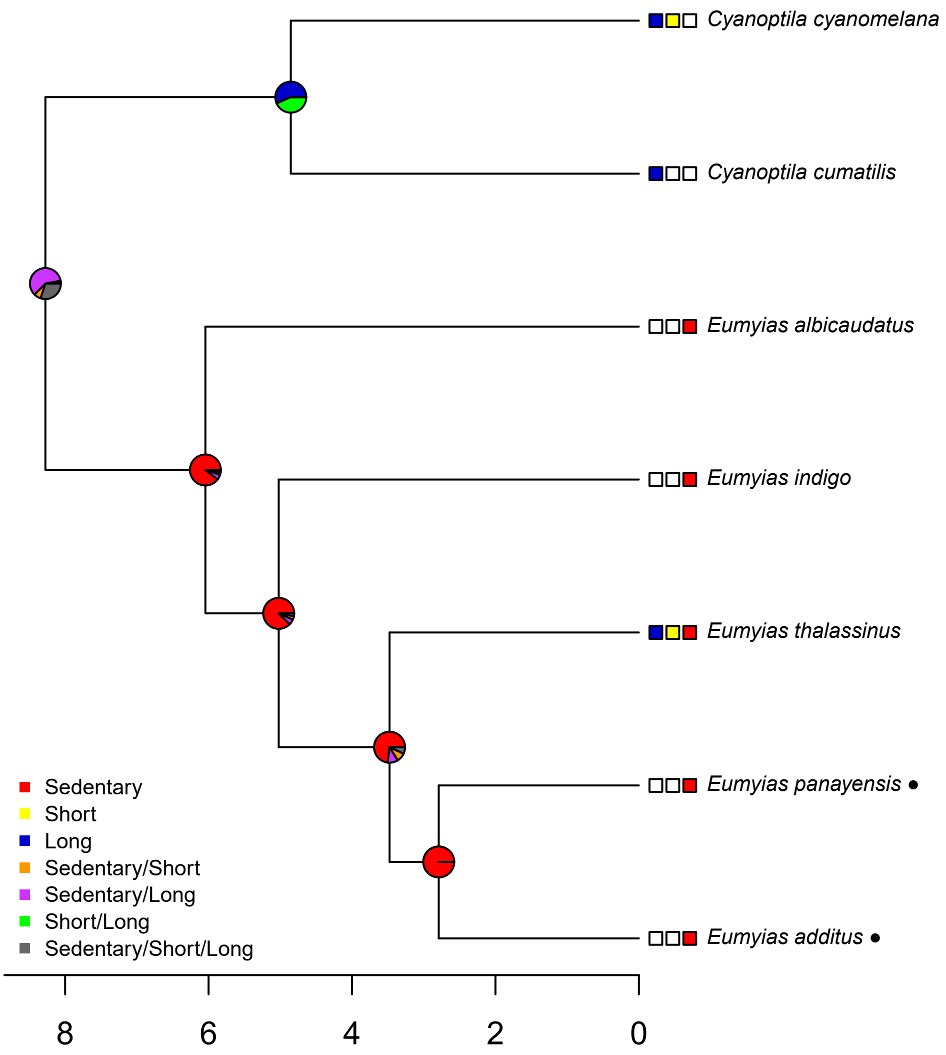

Cyanoptila cyanomelana
Cyanoptila cumatilis
Eumyias albicaudatus
Eumyias indigo
Eumyias thalassinus
Eumyias panayensis ●
Eumyias additus ●
Sedentary
Short
Long
Sedentary/Short
Sedentary/Long
Short/Long
Sedentary/Short/Long
8
6
4
2
0

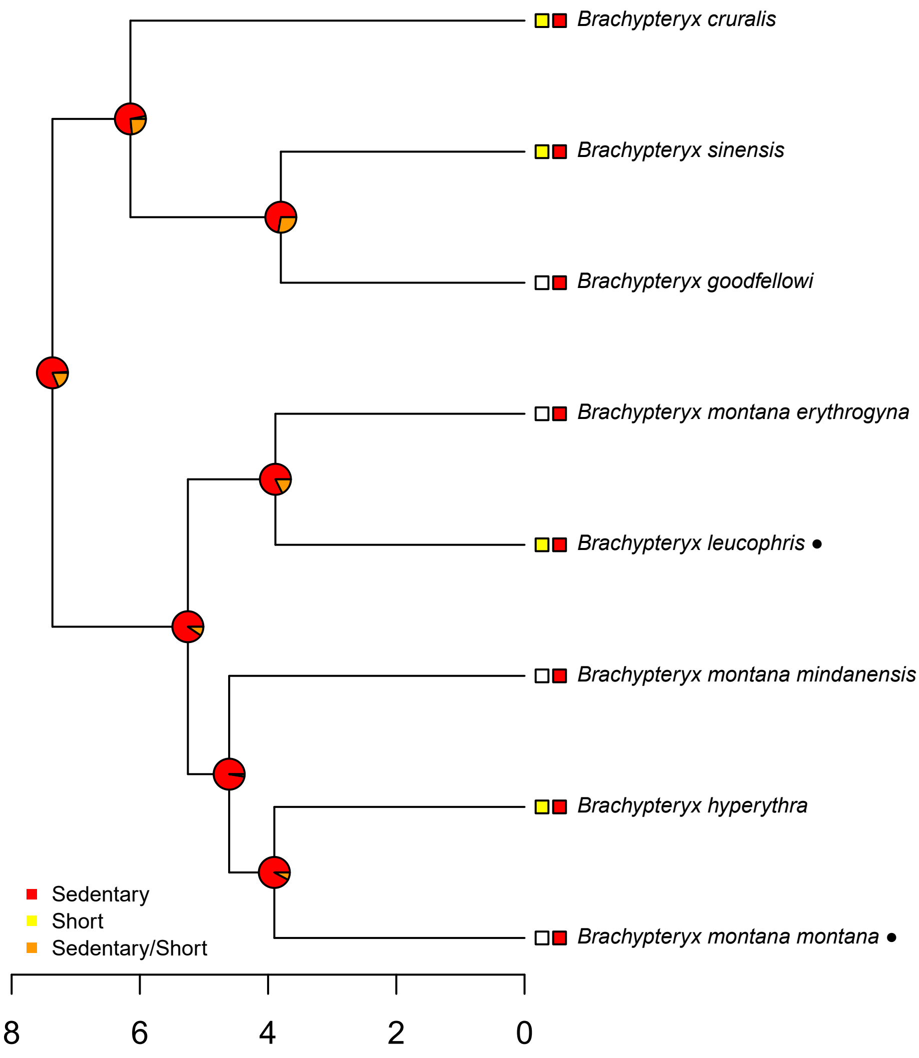
Brachypteryx cruralis
Brachypteryx sinensis
Brachypteryx goodfellowi
Brachypteryx montana erythrogyna
Brachypteryx leucophris ●
Brachypteryx montana mindanensis
Brachypteryx hyperythra
Brachypteryx montana montana ●
Sedentary
Short
Sedentary/Short
8
6
4
2
0

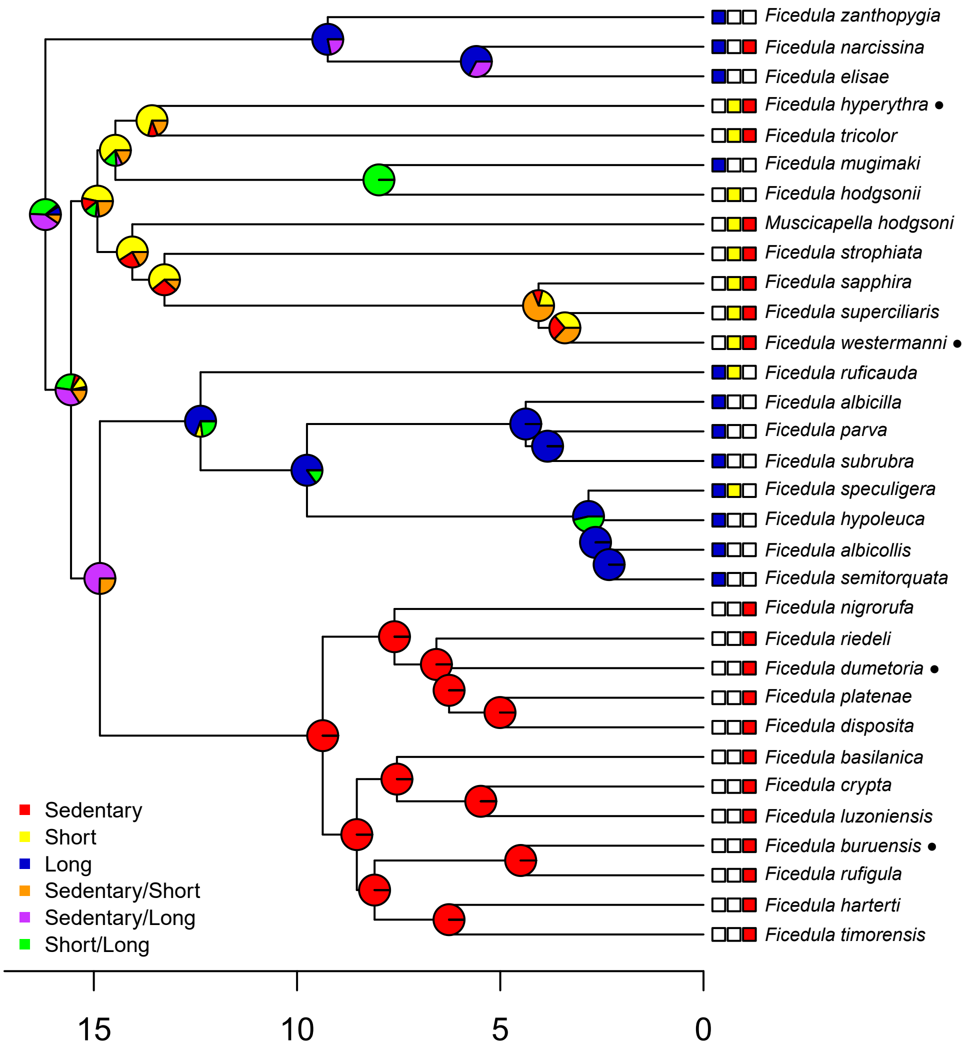

Ficedula zanthopygia
Ficedula narcissina
Ficedula elisae
Ficedula hyperythra ●
Ficedula tricolor
Ficedula mugimaki
Ficedula hodgsonii
Muscicapella hodgsoni
Ficedula strophiata
Ficedula sapphira
Ficedula superciliaris
Ficedula westermanni ●
Ficedula ruficauda
Ficedula albicilla
Ficedula parva
Ficedula subrubra
Ficedula speculigera
Ficedula hypoleuca
Ficedula albicollis
Ficedula semitorquata
Ficedula nigrorufa
Ficedula riedeli
Ficedula dumetoria ●
Ficedula platenae
Ficedula disposita
Ficedula basilanica
Ficedula crypta
Ficedula luzoniensis
Ficedula buruensis ●
Ficedula rufigula
Ficedula harterti
Ficedula timorensis
Sedentary
Short
Long
Sedentary/Short
Sedentary/Long
Short/Long
15
10
5
0

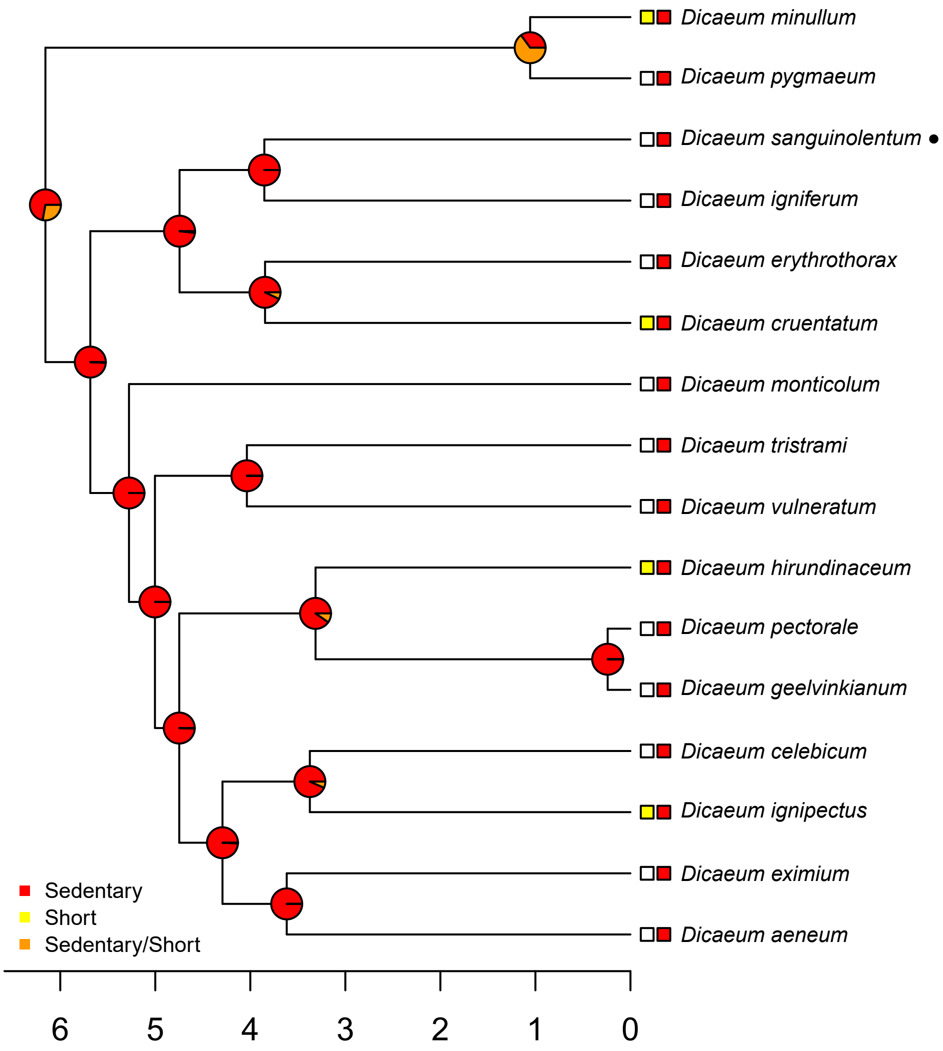

Dicaeum minullum
Dicaeum pygmaeum
Dicaeum sanguinolentum ●
Dicaeum igniferum
Dicaeum erythrothorax
Dicaeum cruentatum
Dicaeum monticolum
Dicaeum tristrami
Dicaeum vulneratum
Dicaeum hirundinaceum
Dicaeum pectorale
Dicaeum geelvinkianum
Dicaeum celebicum
Dicaeum ignipectus
Dicaeum eximium
Dicaeum aeneum
Sedentary
Short
Sedentary/Short
6
5
4
3
2
1
0

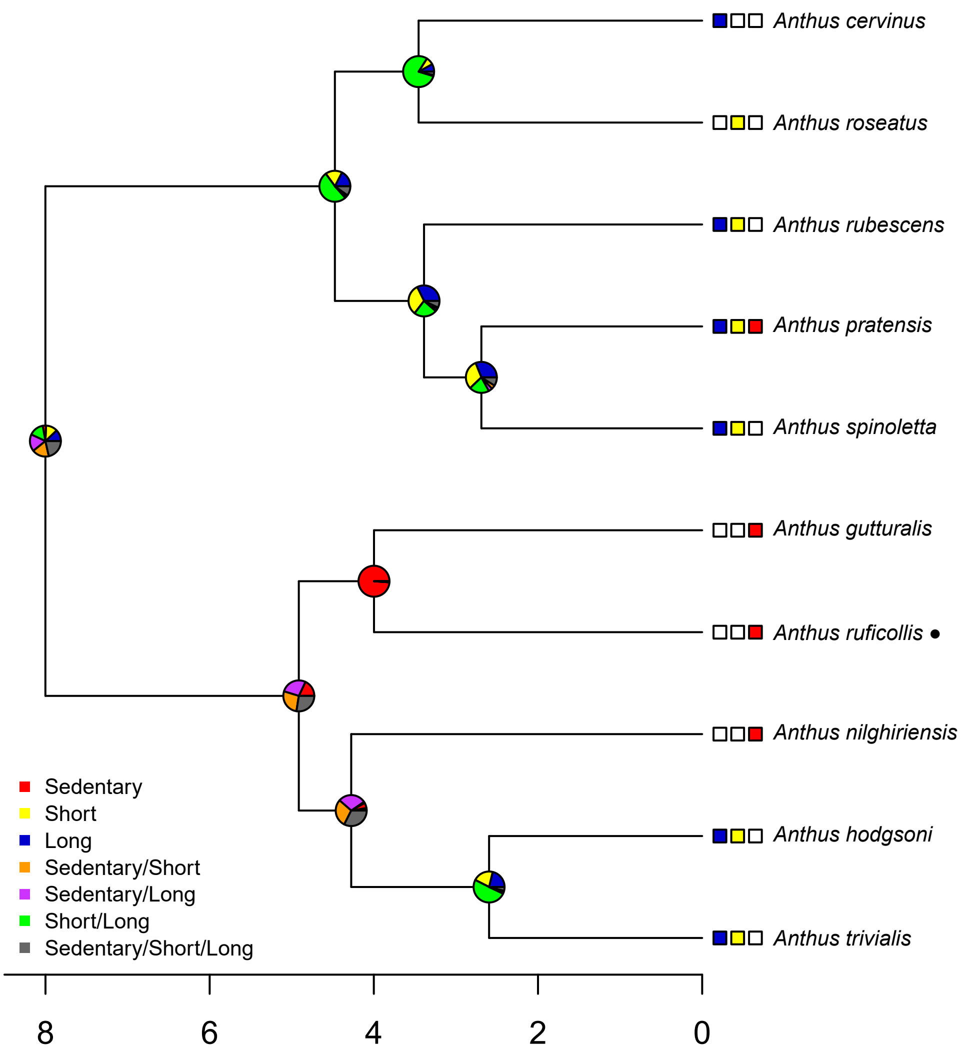
Anthus cervinus
Anthus roseatus
Anthus rubescens
Anthus pratensis
Anthus spinoletta
Anthus gutturalis
Anthus ruficollis •
Anthus nilghiriensis
Anthus hodgsoni
Anthus trivialis
Sedentary
Short
Long
Sedentary/Short
Sedentary/Long
Short/Long
Sedentary/Short/Long
8
6
4
2
0

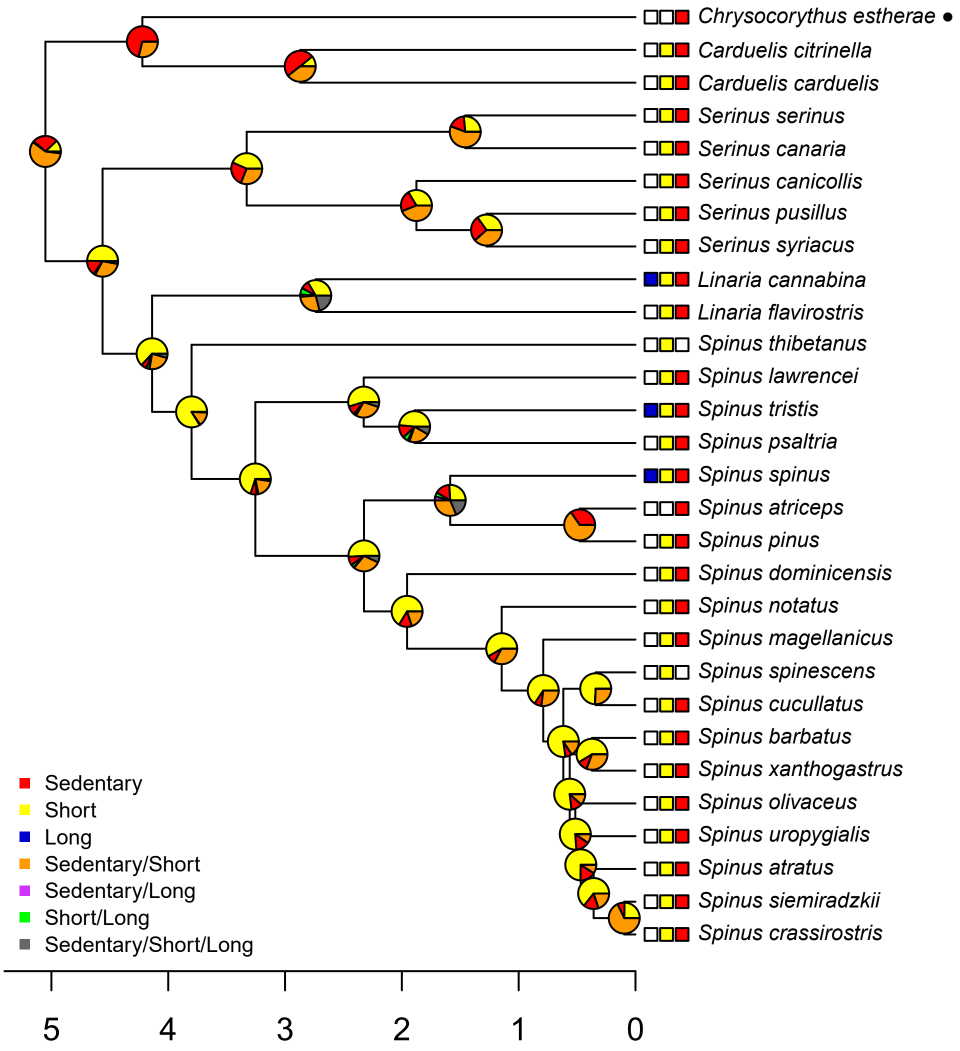
Chrysocorythus estherae •
Carduelis citrinella
Carduelis carduelis
Serinus serinus
Serinus canaria
Serinus canicollis
Serinus pusillus
Serinus syriacus
Linaria cannabina
Linaria flavirostris
Spinus thibetanus
Spinus lawrencei
Spinus tristis
Spinus psaltria
Spinus spinus
Spinus atriceps
Spinus pinus
Spinus dominicensis
Spinus notatus
Spinus magellanicus
Spinus spinescens
Spinus cucullatus
Spinus barbatus
Spinus xanthogastrus
Spinus olivaceus
Spinus uropygialis
Spinus atratus
Spinus siemiradzkii
Spinus crassirostris
Sedentary
Short
Long
Sedentary/Short
Sedentary/Long
Short/Long
Sedentary/Short/Long
5
4
3
2
1
0
