## Supplementary File 5 for "The formation of the Indo-Pacific montane avifauna"

**Supplementary File 5.** Ancestral geographic range curves for Eurasian-origin species. Curves link composite region scores for nodes in the ancestral geographic range reconstructions. Dark blue lines connect posterior probabilities of Palearctic occurrence, light blue lines Indomalayan occurrence.

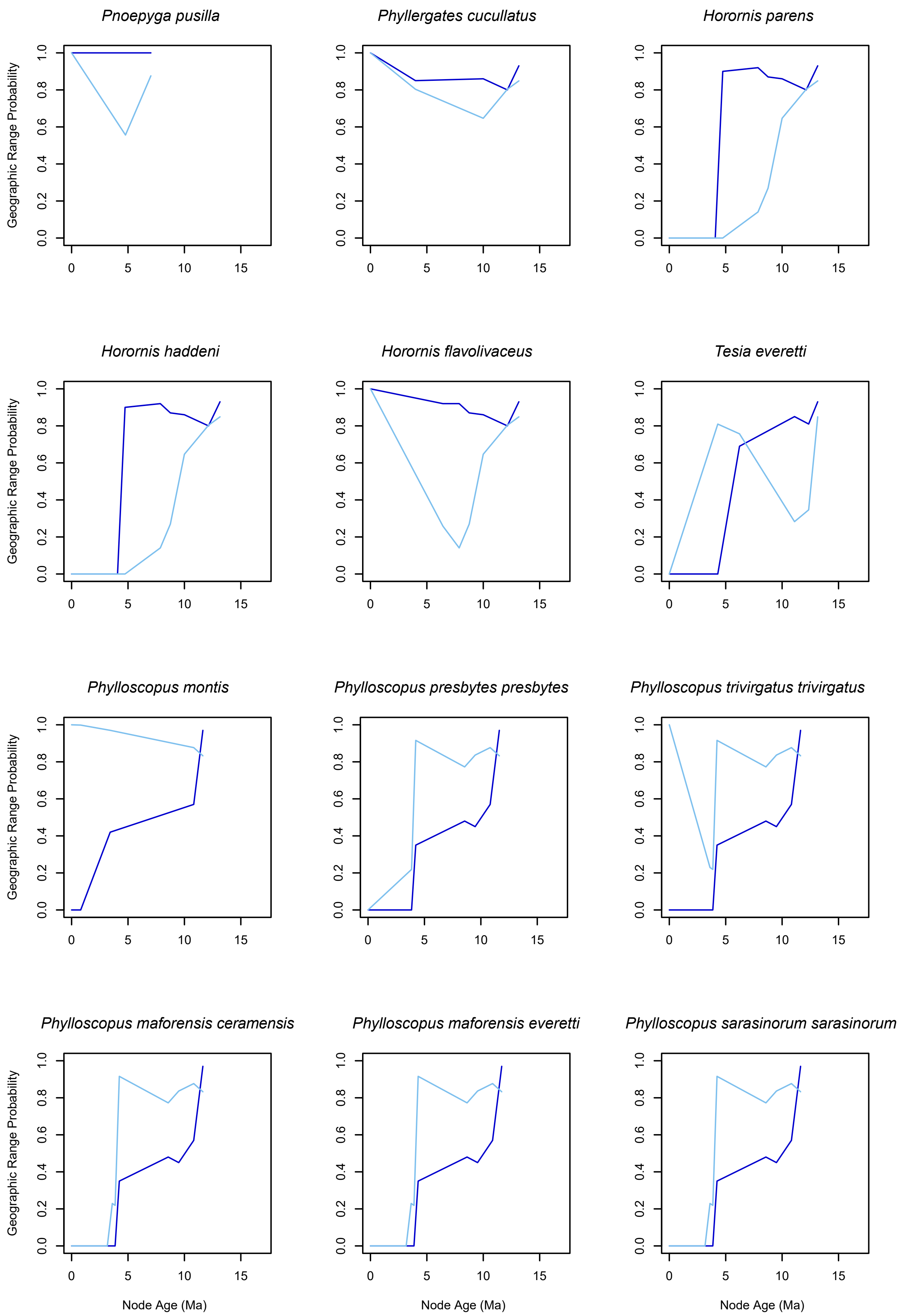
Pnoepyga pusilla
Phyllergates cucullatus
Horornis parens
Horornis haddeni
Horornis flavolivaceus
Tesia everetti
Phylloscopus montis
Phylloscopus presbytes presbytes
Phylloscopus trivirgatus trivirgatus
Phylloscopus maforensis ceramensis
Phylloscopus maforensis everetti
Phylloscopus sarasinorum sarasinorum
Geographic Range Probability
0.0
0.2
0.4
0.6
0.8
1.0
0
5
10
15
Node Age (Ma)

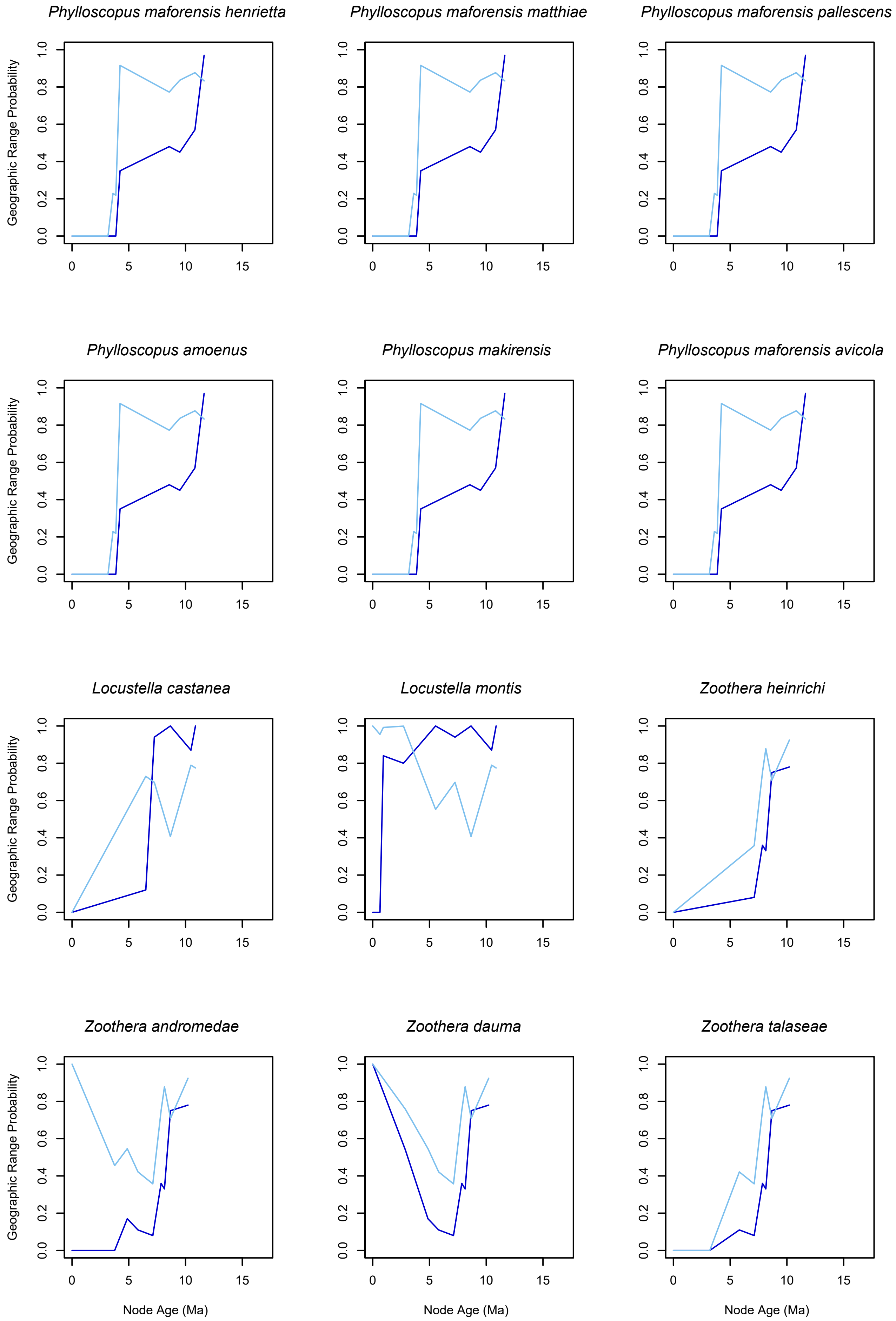
Phylloscopus maforensis henrietta
Phylloscopus maforensis matthiae
Phylloscopus maforensis pallescens
Phylloscopus amoenus
Phylloscopus makirensis
Phylloscopus maforensis avicola
Locustella castanea
Locustella montis
Zoothera heinrichi
Zoothera andromedae
Zoothera dauma
Zoothera talaseae
Geographic Range Probability
0.0
0.2
0.4
0.6
0.8
1.0
0
5
10
15
Node Age (Ma)

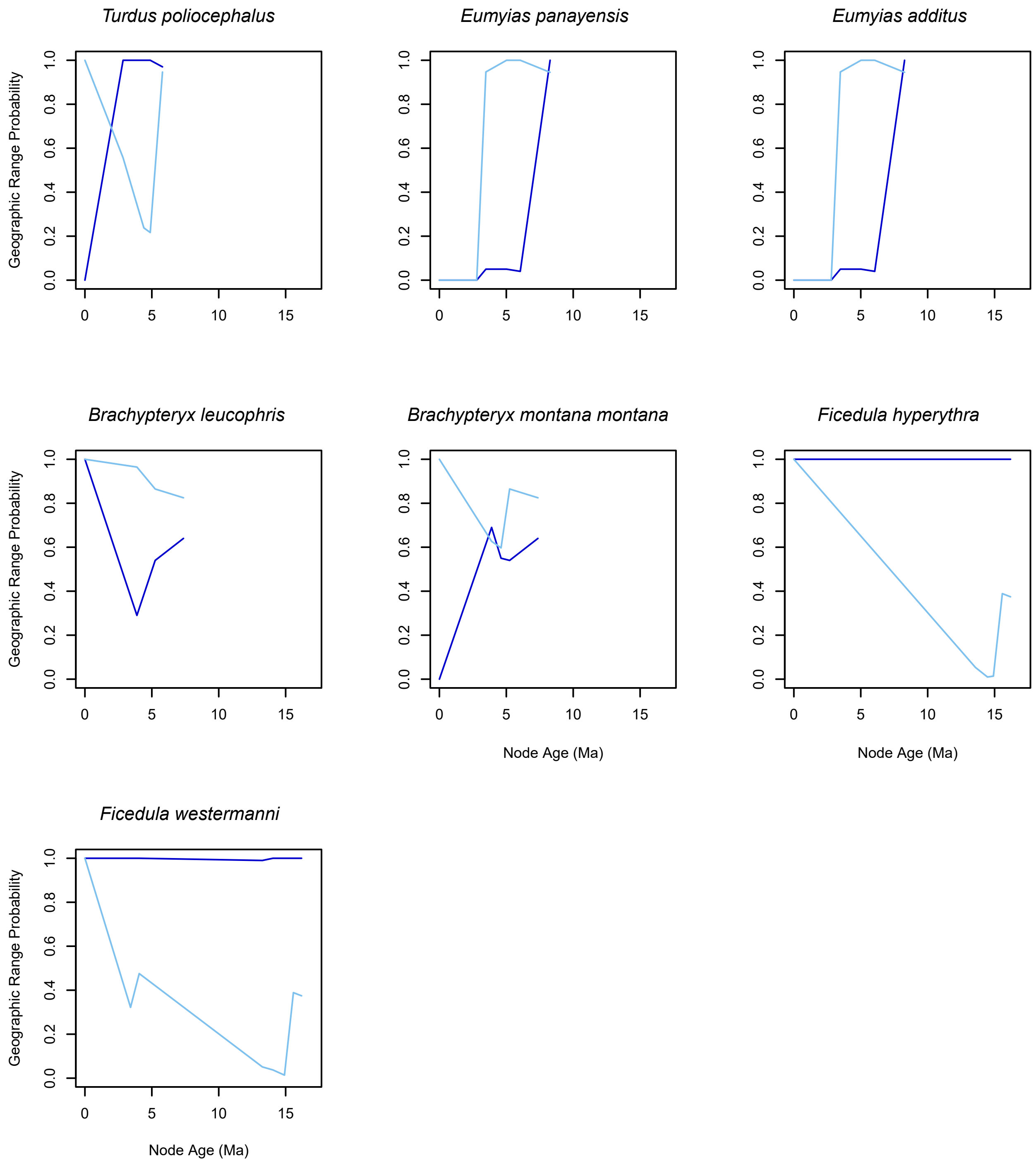
Turdus poliocephalus
Eumyias panayensis
Eumyias additus
Brachypteryx leucophris
Brachypteryx montana montana
Ficedula hyperythra
Ficedula westermanni
Geographic Range Probability
Node Age (Ma)
0.0
0.2
0.4
0.6
0.8
1.0
0
5
10
15
